# A Paracrine Dietary Lipid Axis Constrains Antitumor Immunity in Liver Cancer

**DOI:** 10.64898/2026.06.25.734592

**Authors:** Nicolae Ciobu, Reena Kumari, Jeevotham S. Kumar, Ugne Balaseviciute, Maria Iftesum, Jonathan Mitchell, Justin Ruiz, Sara Flowers, Kaila Nishikawa, Guillem Cano-Segarra, Anna Vila-Escoda, Yang Xiao, Aiden M. Phoebe, Raul Navaridas, Marcella Steffani, Durga P. Gannamedi, Jenny Jin, Bruno Cogliati, Michelle Saoi, Ritchie Ly, Joyce Ogidigo, Monica Rodriguez-Silva, Marta Pardo, Ella Pokrifka, Luis A. Almanza, Simoni Tiano, Erin C. Bush, Renu Nandakumar, Ghassan K. Abou-Alfa, Roser Pinyol, Mara Monetti, David B. Lombard, Defne Bayik, Dionysios C. Watson, Xiaoqiong Wang, Patricia D. Jones, Brent R. Stockwell, Robert F. Schwabe, James J. Galligan, Paul B. Romesser, Yael David, Manas R. Gartia, Josep M. Llovet, Viraj R. Sanghvi

## Abstract

Overnutrition-related liver dysfunction and cancer are increasingly prevalent and highly resistant to immunotherapy. While metabolic dysregulation is a hallmark of hepatocellular carcinoma (HCC), how nutrient overload impairs antitumor immunity remains unclear. Here, we show that short-term Western diet (WD) exposure drives near-complete loss of CD8⁺ T cell infiltration and antitumor function in HCC. We identify dietary linoleic acid (LA), the most abundant ω-6 fatty acid, as the dominant immunosuppressive driver. Cancer cell-restricted FADS2-mediated desaturation of LA to longer-chain ω-6 PUFAs drives their accumulation in the tumor interstitial fluid, suppressing infiltrating CD8⁺ T cells via lipid peroxidation. FADS2 inhibition restores CD8⁺ T cell function and sensitizes WD-driven HCC to PD-1-based immunotherapy. Further, the Parkinson’s disease-associated deglycase DJ-1 protects LA-handling proteins from methylglyoxal-mediated glycation, sustaining tumoral immunosuppressive PUFA production. Across multiple independent human MASLD-HCC cohorts, LA metabolic activity correlates with CD8⁺ T cell impairment, immune exclusion, and immunotherapy resistance. Overall, these studies identify a dietary lipid axis as a therapeutically actionable vulnerability in WD-associated HCC.

## Introduction

Hepatocellular carcinoma (HCC) is the most common and deadly form of primary liver cancer, accounting for nearly 10% of global cancer-related mortality^1^. The incidence of metabolic dysfunction–associated steatotic liver disease (MASLD)-driven HCC is rising at an alarming rate, largely driven by the widespread adoption of Western dietary (WD) patterns characterized by excessive intake of saturated and unsaturated fats, refined sugars, and processed carbohydrates, combined with increasingly sedentary lifestyles^1–4^. Despite this well-established link between WD and HCC, the molecular mechanisms by which excess dietary lipids promote HCC remain poorly understood. Previous studies have shown that WD-associated HCC exhibits key differences from other etiologies, most notably a profoundly immunosuppressive tumor microenvironment (TME)^5–7^. In particular, WD-associated HCC displays an immune-excluded phenotype, where CD8⁺ T cells are poorly activated, metabolically exhausted, and functionally reprogrammed^6,8,9^. This CD8⁺ T cell dysfunction facilitates persistent immune escape and underlies the purported failure of current immunotherapies in the context of steatotic liver disease^5,10,11^.

Tumor metabolism and immune function are tightly coupled, with nutrient partitioning in the TME dictating CD8⁺ T cell fate^12,13^. While several animal studies – often relying on subcutaneous transplants of melanoma and breast cancer cells in obese mice – have demonstrated that chronic overnutrition suppresses T cell function, the proposed mechanisms vary widely, ranging from impaired amino acid metabolism^14^, skewed lipid partitioning and lipid deficiency in CD8^+^ T cells^15^ and, conversely, excessive lipid uptake and accumulation in CD8^+^ T cells^16^. The extent and underlying mechanism of inhibition also depend on the dietary lipid formulation and the abundance of specific lipid species^17^. The liver is a uniquely immunoprivileged and metabolically active organ that exhibits pronounced metabolic zonation – a spatial organization of hepatocellular metabolic phenotypes that shapes the immunological tone of the hepatic microenvironment in ways that are fundamentally distinct from other tissues. Subcutaneous transplantation of tumor cells bypasses the liver’s intrinsic metabolic architecture, portal nutrient delivery, resident immune cell composition, and zonation-dependent signaling gradients that collectively define the immunosuppressive landscape of HCC. Such ectopic tumor models therefore are unlikely to faithfully recapitulate the unique hepatic niche. As such autochthonous models in which tumors arise within the liver parenchyma are essential to accurately interrogate the diet–immunity interaction in this disease. Moreover, the commonly used dietary models of MASLD-associated HCC reproduce key features of liver steatosis and fibrosis but are limited by weak and heterogeneous tumor immunogenicity and an inability to distinguish tumor-intrinsic from systemic effects^18^. A clearer understanding of how these mechanisms operate in the context of liver-specific TME is essential for identifying diet-immune interactions that underlie CD8^+^ T cell failure in tumor control.

A key molecular consequence of chronic glucose overnutrition is the excessive generation of methylglyoxal (MGO), a reactive glycolytic by-product that modifies proteins through non-enzymatic glycation, leading to the accumulation of advanced glycation end-products (AGEs)^19–21^. DJ-1 (encoded by *PARK7*), a conserved glyoxalase, counteracts MGO-mediated protein glycation and functions as a key metabolic stress sensor^22–24^. In contrast to enzymatic glycosylation, glycation is stochastic, often deleterious, and markedly elevated in hyperglycemia and metabolically stressed tissues such as liver and adipose^25,26^. MGO substrates include proteins involved in diverse cellular processes, such as chromatin regulation (histones)^19,27^, redox homeostasis^28^ (KEAP1), chaperone function (HSP90)^29^, and DNA repair (BRCA2)^30^. MGO is elevated in MASLD and MASLD-associated HCC, where it can profoundly influence both tumorigenesis and antitumor immune responses^25^. Despite the clear link between glycation, MASLD, and cancer risk, the role of DJ-1 in regulating metabolic stress, lipid handling, and hepatocarcinogenesis remains poorly defined.

## Results

### Western diet impairs neoantigen-driven antitumor immunity to accelerate HCC progression

We first established a hydrodynamic tail vein injection (HDTV)-based autochthonous mouse model of HCC that is responsive to WD and recapitulates key metabolic, transcriptional, and immunological features of MASLD-associated human HCC^31^. Briefly, we used HDTV to introduce *MYC* and sg-p53 (referred to as MP53 from here-on) into murine hepatocytes, as previously described^32^, followed by randomized exposure to isocaloric standard diet (SD) or obesogenic and steatogenic WD *ad libitum* **(Figure 1a; Figure S1a-d)**. As expected, multifocal MP53-driven HCC developed under both conditions; however, WD-fed mice exhibited markedly accelerated tumor growth and reduced overall survival in both female (median survival: SD – 30 d, WD – 24 d; *p* = 0.001) and male (median survival: SD – 56 d, WD – 40 d; *p* = 0.003) mice **(Figure 1b; Figure S1e)**; extended survival observed in males likely reflects a lower effective transposon dose delivered during injection. *Ex vivo* and histopathological analyses of time-matched livers revealed no major differences in tumor grade (Grade II (moderately differentiated) with marked nuclear atypia) **(Supplementary Table 1)**; however, WD-fed mice exhibited a marked increase in tumor burden and lesion size as well as demonstrated lipid droplet accumulation and mild fibrosis in tumor-adjacent tissue, consistent with features of the corresponding human disease^3,7^ **(Figure S1f-h)**. Longitudinal *in vivo* bioluminescence imaging of luciferase-expressing MP53 HCCs in both multifocal HDTV and focal electroporation-based models^33^ confirmed accelerated tumor growth under WD conditions **(Figure S1i-k)**. Gene set enrichment analysis (GSEA) of bulk transcriptomic data indicates that our model faithfully recapitulates deregulated gene expression programs characteristic of MASLD/MASH-HCC across murine models and human disease, especially metabolic and lipid-related programs^34–36^ **(Figure S2a, b, f, g)**. Consistent with lipid dysregulation and MASLD-like features, WD-fed HCCs exhibited increased intratumoral triglycerides **(Figure S8g)**. Importantly, similar WD-dependent disease acceleration and activation of these transcriptional programs were observed in an independent HCC model driven by MYC overexpression and steatosis-promoting *Pten* deletion^36^ (*MYC*/sg-Pten; referred to as MPn from here-on) **(Figure 1c; Figure S1l, m, 2c-e)**. Together, these data establish that WD accelerates HCC progression in HDTV-based mouse models that faithfully recapitulates key features of human MASLD-associated HCC.

**Figure 1:**
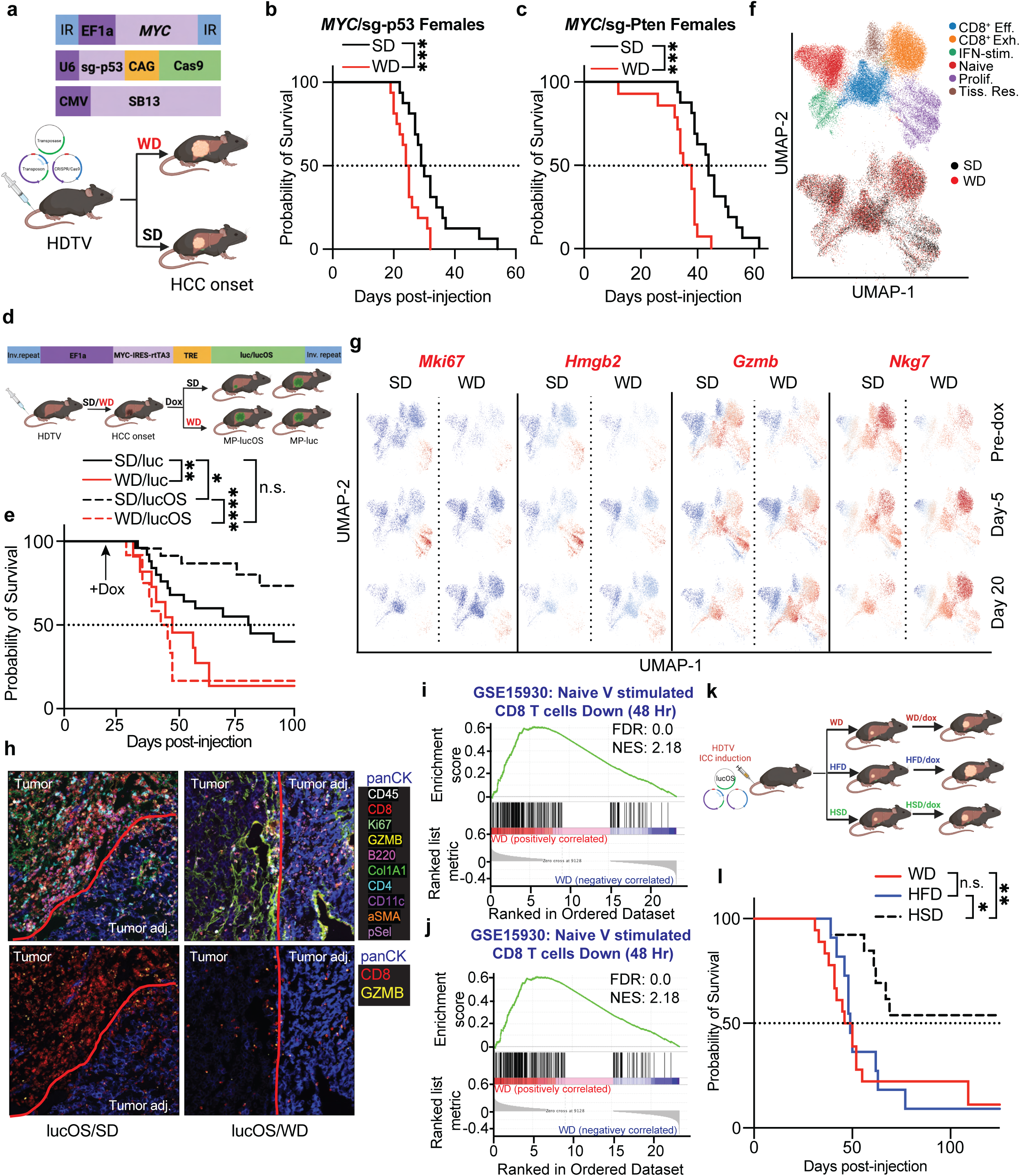
Western Diet drives T cell impairment in liver cancer *in vivo*. **(a)** Schematic of the HDTV-based murine model of SD– and WD-fed HCC; **(b & c)** Kaplan–Meier survival curves of female mice bearing MP53 (SD: n=16, WD: n=16) (b) and MPn (SD: n=16, WD: n=14) (c) HCCs; **(d)** Plasmid map of doxycycline (dox)-inducible luc or lucOS expression (top) and our experimental approach (bottom); **(e)** Kaplan-Meier survival curves of mice bearing luc– or lucOS-MP53 HCCs under SD or WD *ad libitum* (SD/Luc: n=25; WD/Luc: n=11; SD/lucOS: n=17; WD/lucOS: n=12); arrow indicates initiation of dox treatment (day 21 post-injection); **(f)** UMAP of tumour-infiltrating CD8^+^ T cells from MP53-lucOS HCCs colored by subtype (top) and diet (bottom) (SD: 20,200 cells, WD: 19,138 cells), Eff: Effector; Exh: Exhausted; Prolif: Proliferative; Tiss. Res: Tissue resident; **(g)** UMAP plots depicting expression of proliferative (*Mki67*, *Hmgb2*) and effector (*Gzmb*, *Nkg7*) marker genes in tumour-infiltrating CD8⁺ T cells across indicated dox timepoints; **(h)** Representative multiplexed immunofluorescence images of SD– and WD-fed MP53-lucOS tumours; red line delineates tumour border; **(i & j)** GSEA of tumour-infiltrating CD8⁺ T cells isolated from SD– and WD-fed MP53-lucOS HCCs (SD: n=4, WD: n=5); **(k & l)** Experimental scheme (k) and Kaplan–Meier survival curves of MP53-lucOS tumour-bearing mice under WD, HFD, or HSD (WD: n=9; HFD: n=8; HSD: n=8). ns: not significant, *p < 0.05, **p < 0.01, ***p < 0.001, ****p < 0.0001.

To dissect the immune-diet interaction, we profiled tumor-infiltrating CD45⁺ leukocytes by single-cell RNA sequencing (scRNAseq), revealing no broad compositional changes in immune cell populations between conditions (**Figure S3a; 4a)**. However, T cell subclustering revealed selective diet-dependent remodeling of the CD8⁺ T cell compartment, closely mirroring patterns described in human MASLD-HCC^7^ **(Figure S3b, c; 4b, c)**. In both models, WD shifted the intratumoral CD8⁺ T cell landscape from effector (*Gzmb⁺*, *Prf1⁺*, *Ifng⁺*) and IFN-stimulated (*Isg15⁺*) states toward naive (*Lef1⁺*, *Il7r⁺*) populations **(Figure S3c-e; 4c, d)**. Milo analysis confirmed these observations, identifying significant enrichment of effector neighborhoods under SD and naive neighborhoods under WD in both models **(Figure S3f; 4f)**. Feature plots, condition overlay, and Milo analysis collectively demonstrated that while effector molecule expression distributions, including *Gzmb*, *Prf1*, *Ifng*, and *Tnfa*, were preserved among infiltrating T cells, the relative abundance of effector molecule-high cells was markedly reduced under WD **(Figure S3f, g; 4e, f)**. Additionally, a trend toward increased exhaustion (*Pdcd1⁺*, *Tox^+^*) was observed in a subset of animals under WD, suggesting progressive but heterogeneous T cell dysfunction **(Figure S3c; 4c)**. Collectively, these data indicate that WD impairs early T cell activation and recruitment to the tumor, driving a compositional shift away from effector states toward naive exhausted phenotypes. In line with these, WD-imparted growth advantage is completely lost in immunocompromised NSG mice, and loss of CD8^+^ T cells under SD conditions phenocopied WD-driven tumor acceleration but conferred no additional growth advantage under WD in immunocompetent animals, suggesting that CD8⁺ T cells are already functionally incapacitated under calorigenic settings **(Figure S4g, h)**.

Both MP53 and MPn are known to be low-immunogenic models of HCC with minimal involvement of T cell-driven cytotoxicity^37,38^. To directly study the WD-mediated immune escape in a defined immunogenic context, we established a neoantigen-controlled MP53-HCC model. Using a modified Sleeping Beauty transposon system, we co-expressed MYC and reverse tetracycline transactivator (rtTA3) in a bi-cistronic vector along with either wild-type luciferase (Luc) or an immunogenic luciferase variant (lucOS) encoding three defined neoepitopes, including SIINFEKL from ovalbumin **(Figure 1d)**. When expressed constitutively, these epitopes have previously been shown to elicit robust CD8⁺ T cell–mediated responses in autochthonous lung and liver tumors^37,39^. We hydrodynamically initiated MP53-luc or highly immunogenic MP53-lucOS HCCs and maintained the animals on SD or WD. Doxycycline (dox) was administered post-tumor establishment to induce neoantigen expression. As before, WD accelerated disease progression and reduced survival (median survival: SD = 81 d; WD = 45 d; *p* < 0.004) in mice expressing weakly immunogenic wildtype luciferase **(Figure 1e)**. Under SD conditions, lucOS-expressing tumors were efficiently cleared by CD8⁺ T cells, as indicated by a progressive decline in bioluminescence, resulting in significantly improved survival (median survival: lucOS/SD-undefined, *p* = 0.01 v luc/SD) **(Figure 1e; Figure S5a)**. In contrast, mice not receiving dox supplementation displayed similar tumor progression, confirming minimal background expression and excluding confounding effects from potential transgene leakiness **(Figure S5b)**. Strikingly, however, this survival advantage was entirely lost under WD, indicating that overnutrition overrides neoantigen-driven antitumor immunity, even in the presence of a strong immunogenic driver (median survival: lucOS/WD: 43 d, *p* < 0.0001 v lucOS/SD and > 0.9 v luc/WD) **(Figure 1e; Figure S5c, d)**. Notably, this immune evasion occurred over a relatively short timescale, highlighting the rapid and dominant impact of metabolic stress on impairing CD8⁺ T cell–mediated tumor control. Importantly, this loss of immune protection only occurred when neoantigen expression was induced post-tumor establishment. In contrast, constitutive expression of lucOS from the time of tumor initiation preserved CD8⁺ T cell–mediated clearance as shown before^37^, even when mice were preconditioned with WD prior to hydrodynamic injection **(Figure S5e)**.

To characterize temporal immune dynamics following neoantigen induction, we performed scRNAseq on tumors harvested prior to and at days 5 and 20 post-doxycycline administration. Unsupervised clustering delineated major immune and stromal cell types, with WD-fed tumors exhibiting modest shifts in TME composition, such as increased macrophages and neutrophils, consistent with prior observations^40,41^ **(Figure S5f, g)**. Focused re-clustering of CD8⁺ T cells revealed six transcriptionally distinct subsets: naïve, proliferative, effector, exhausted, tissue-resident, and IFN-stimulated populations **(Figure 1f)**. Notably, at day 5 post-induction, SD-fed tumors showed a marked expansion of both effector and proliferative CD8⁺ T cells, indicative of robust antigen-driven activation – an effect that was substantially blunted in WD-fed tumors **(Figure 1g; Figure S5h)**. By day 20, CD8⁺ T cells in WD-fed MP53-lucOS tumors exhibited upregulation of both proliferative and effector programs alongside sustained exhaustion signatures, indicating that WD delays and dysregulates T cell activation, ultimately promoting dysfunctional immune responses **(Figure 1g; Figure S5h)**. Importantly, by this time point, tumors in SD-fed mice had already regressed or been cleared, whereas WD-fed mice progressed toward lethal and potentially irreversible disease, underscoring the profound immunosuppressive impact of dietary overnutrition **(Figure 1e; Figure S5a)**. Multiplex immunofluorescence, which maintains spatial context, revealed an immune-rich TME in SD-fed immunogenic MP53 HCCs but profoundly immune-depleted under WD. Notably, this difference was largely restricted to tumor tissue and not apparent in adjacent liver, indicating local WD-driven rewiring of the tumor microenvironment **(Figure 1h)**. Consistently, GSEA of bulk RNA-seq data from tumor-infiltrating CD8^+^ lymphocytes (TILs) of SD– and WD-fed animals revealed that even among T cells capable of infiltrating WD-fed tumors, transcriptional programs associated with T cell proliferation, metabolism, and activation were markedly suppressed compared to their SD counterparts **(Figure 1i, j; Figure S6a-d)**. WD-driven impairment of CD8⁺ T cell-mediated antitumor cytotoxicity was similarly observed in the MPn-lucOS model **(Figure S6e).** To identify the dietary macronutrient responsible for immune suppression, we subjected MP53-lucOS-injected animals to the WD with high fat and sugar content, the same high-fat diet (HFD) lacking sugar-water supplementation, or low-fat SD supplemented with sugar-rich drinking water (HSD). HFD phenocopied WD-induced immune suppression (median survival: WD — 48 d, HFD — 49 d, *p* > 0.8), whereas HSD showed extended survival compared to both lipid-rich diets (median survival — undefined, *p* < 0.005 v WD and *p* < 0.02 v HFD) **(Figure 1k, l)**. Collectively, these findings indicate that diet-supplied lipids are the dominant driver of immune exclusion and T cell impairment in HCC progression.

### Dietary linoleic acid fuels tumor ω-6 PUFA biosynthesis to drive immune evasion in HCC

To unravel the mechanism by which WD drives immune evasion, we performed untargeted metabolomics and lipidomics, revealing significant accumulation of linoleic acid (LA; C18:2 ω6) in WD-fed MP53 and MPn tumors, with minimal or inconsistent changes in saturated or monounsaturated fatty acids **(Figure 2a, b; Figure S7a)**. A similar increase in LA abundance was observed in a non-genetic hepatocarcinogenesis model driven by long-term WD feeding followed by CCl₄ treatment^42^ **(Figure S7b, c)**. Raman spectroscopy, which enables label-free detection of lipids through characteristic C–H (∼2850–2930 cm⁻¹) and C=C (∼1655 cm⁻¹) vibrational modes^43,44^, confirmed increased LA accumulation in WD-fed immunogenic tumors relative to SD-fed counterparts **(Figure 2c; Figure S7d)**. Given that LA is the predominant diet-derived ω-6 PUFA in the WD, and cannot be synthesized *de novo*, its selective accumulation in WD-fed tumors prompted us to investigate its role in immune suppression. Towards that, MP53-lucOS-injected mice were fed an alternative WD enriched in palmitic and oleic acids but deficient in LA, resulting in significantly prolonged survival compared to its non-immunogenic counterpart (median survival: lucOS/WD LA-depleted = 75 d; luc/WD LA-depleted = 47 d; *p* = 0.037) **(Figure 2d)**. Conversely, a custom calorie-matched diet in which fat calories were derived mostly from LA phenocopied the full-strength WD, confirming that LA is the dominant driver of WD-induced immune suppression (median survival: full WD – 49 d; LA-enriched WD – 45 d; *p* = n.s.) **(Figure 2e)**. LA is activated to linoleoyl-CoA in hepatocytes by long-chain acyl-CoA synthetases, such as ACSL1 and ACSL5^45–47^, enabling its mitochondrial β-oxidation or elongation and desaturation that generate longer-chain polyunsaturated fatty acids for incorporation into complex lipids, membrane biosynthesis, or release as free fatty acids following de-esterification by acyl-CoA thioesterases^48,49^ **(Figure S7f)**. Notably, untargeted metabolomics, targeted lipidomics and Raman scattering revealed accumulation of free LA-derived ω-6 PUFAs, including γ-linolenic acid (GLA, C18:3 ω-6), dihomo-γ-linolenic acid (DGLA; 20:3 ω-6), arachidonic acid (AA; 20:4 ω-6), docosatetraenoic acid (DTA; 22:4 ω-6), and docosapentenoic acid (DPA; 22:5 ω-6), in WD-fed MP53, MPn and WD/CCl₄ tumors, with corresponding enrichment in tumor interstitial fluid suggesting tumor-derived secretion **(Figure 2a-c; Figure S7a–e)**. To trace the fate of dietary LA within the TME *in vivo*, BODIPY-tagged LA was administered to WD-fed MP53 and MPn tumor-bearing mice and its uptake quantified across immune and non-immune cell populations. Cancer cells exhibited predominant LA incorporation, whereas lymphoid and myeloid compartments showed minimal uptake, establishing a tumor-intrinsic origin of LA-derived ω-6 PUFA production **(Figure S7g, h)**. Collectively, these data identify LA as a major immunosuppressive dietary lipid.

**Figure 2:**
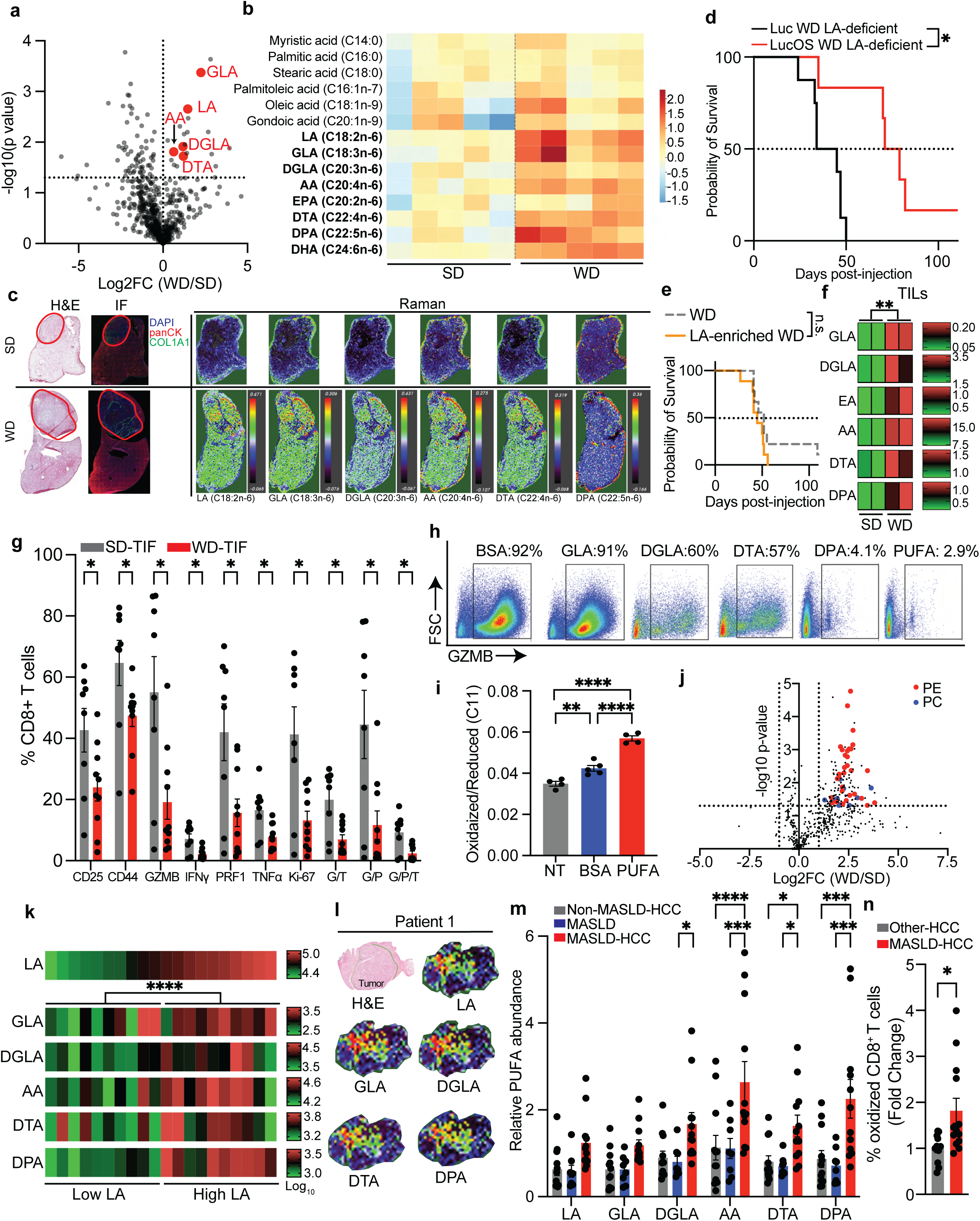
Dietary LA-derived ω-6 PUFAs suppress CD8⁺ T cell function in HCC. **(a)** Volcano plot of differentially abundant metabolites in SD– and WD-fed MP53 tumours by untargeted metabolomics (SD: n=3; WD: n=3); LA pathway metabolites highlighted in red; **(b)** Heatmap of targeted lipidomics showing indicated saturated, monounsaturated, and polyunsaturated fatty acids in MP53 HCCs under SD and WD (SD: n=5; WD: n=5). EPA: eicosadienoic acid (C20:2 n-6), DHA: docosahexaenoic acid (C24:6, n-6); **(c)** Spatial metabolomics of LA pathway metabolites by Raman spectroscopy; H&E staining and immunofluorescence provided for tumour localization (red circles); **(d)** Kaplan–Meier survival of MP53-Luc and –lucOS female mice under LA-deficient WD (Luc: n=8, lucOS: n=6); **(e)** Kaplan–Meier survival of MP53-lucOS-bearing female mice under WD or a matched LA-enriched WD (WD: n=12; LA-enriched WD: n=9); **(f)** Targeted lipidomics of LA-derived fatty acids in tumour-infiltrating CD8⁺ T cells from SD– or WD-fed MP53 HCCs (SD: n=2; WD: n=2); **(g)** CD8⁺ T cell activation marker expression following stimulation in the presence of TIF from SD– (n = 8) or WD-(n = 10) fed MP HCCs, G: granzyme B; P: perforin; T: TNFα; **(h)** Granzyme B expression in CD8⁺ T cells stimulated *ex vivo* in the presence of BSA or indicated ω-6 lipids (200 µM, 72 h), PUFA: equimolar mixture of all five ω-6 lipids (50 µM); **(i)** Oxidized:reduced C11-Bodipy in CD8⁺ T cells stimulated in the presence of BSA or ω-6 PUFA mix (50 µM each, 72 h, n≥4); NT: unstimulated control; NT: no stimulation; **(j)** Volcano plot of differentially abundant lipids in CD8⁺ T cells stimulated with SD– or WD-derived TIF (n=3); highlighted species contain AA, DTA, and DPA; PE: phosphatidylethanolamine; PC: phosphatidylcholine; **(k)** Heatmap of LA and LA-derived fatty acid abundance in HCC patient tumours (MSKCC cohort) (n=20) by targeted lipidomics, stratified by LA abundance; **(l)** Spatial metabolomics of LA pathway metabolites in MASLD-HCC patient tumour samples by Raman spectroscopy; H&E staining provided for tumour identification (green boundaries define tumour border); **(m)** Relative abundance of LA and derived PUFAs in serum from non-MASLD-HCC (n=11), MASLD (n=8), and MASLD-HCC (n=12) patients (University of Miami Cohort); **(n)** Fold change in lipid peroxidation of peripheral blood CD8⁺ T cells by C11-Bodipy staining in MASLD-HCC (n=13) relative to non-MASLD-HCC (n=11) patients. Results are shown as mean ± SEM. ns: not significant, *p < 0.05, **p < 0.01, ***p < 0.001, ****p < 0.0001.

We next asked whether these tumor-derived, LA intermediates contribute to immune suppression through a cancer-cell non-autonomous mechanism. To this end, although LA abundance in T cells did not show a consistent change with diet **(Figure S7i)**, TILs from WD-fed animals accumulated increased levels of LA-derived PUFAs **(Figure 2f)**. T cells lack expression of enzymes required for LA desaturation and elongation^50^, and consistent with this, incubation of T cells with GLA *in vitro* did not generate longer-chain desaturated PUFAs such as DTA and DPA, indicating that these lipids are acquired rather than synthesized by T cells **(Figure S7j)**. To further establish that tumor-derived LA metabolites are transferred to CD8⁺ T cells, activated CD8⁺ T cells were exposed to conditioned medium from alkyne-LA-pulsed MP53 murine HCC cells. Fluorescence signal in recipient lymphocytes confirmed transfer of LA-derived metabolites from cancer cells to CD8⁺ T cells **(Figure S7k)**.

To test whether ω-6 PUFA-rich WD-tumor interstitial fluids (TIFs) were immunosuppressants, CD8⁺ T cells were activated in the presence of TIF from SD– or WD-fed tumors. While SD-TIF supported robust activation (CD25, CD44, GZMB, IFNγ, TNFα, Ki-67), WD-TIF significantly attenuated T cell activation, cytokine production, and polyfunctionality **(Figure 2g)**. Direct exposure to individual LA-derived metabolites but not LA further confirmed these tumor-derived products as the key T cell suppressive agents **(Figure 2h; Figure S8a-d)**. Notably, we observed a progressive increase in T cell suppression correlating with both fatty acid chain length and degree of desaturation. GLA, containing 18 carbons and 3 double bonds, exhibited the weakest suppressive effect, whereas DPA, with 22 carbons and 5 double bonds, produced the most pronounced inhibition of T cell activation **(Figure 2h; Figure S8a, b)**. DGLA and DTA induced intermediate suppressive effects **(Figure 2h; Figure S8a, b)**. When combined, these metabolites exerted additive suppression, further attenuating CD8⁺ T cell cytokine production, proliferation, and polyfunctionality **(Figure 2h; Figure S8a, b)**. Moreover, etomoxir or thioridazine did not protect T cells from ω-6-mediated suppression, ruling out enhanced mitochondrial or peroxisomal oxidation as immunosuppressive mechanisms **(Figure S8e)**. Importantly, these desaturated ω-6 lipids also inhibited T cell activation by PMA/Ionomycin, ruling out impediment of proximal TCR signaling **(Figure S8f, g)**. Based on the structural features of these lipids – highly unsaturated and oxidizable – we next asked if they disrupt CD8⁺ T cell function through lipid peroxidation and mitochondrial impairment. Indeed, ω-6 lipid–treated T cells exhibited an elevated ratio of oxidized to reduced C11-BODIPY, indicative of heightened lipid peroxidation **(Figure 2i)**, and subsequent mitochondrial depolarization **(Figure S8h)**. Untargeted lipidomics revealed that T cells activated in WD-TIF exhibited altered membrane phospholipid composition, with increased PE and PC species containing the peroxidation-prone fatty acids AA, DTA, and DPA, whereas WD-influenced cancer cells exhibited reduced levels of these phospholipid species **(Figure 2j; Figure S8i)**. This divergent lipid redistribution suggests preferential accumulation of peroxidation-prone phospholipids in T cells within the WD TME. Consistent with this, activation of CD8⁺ T cells *ex vivo* in the presence of ω-6 PUFAs resulted in decreased cell viability, which was partially restored by pharmacological ferroptosis inhibition **(Figure S8j)**, suggesting that ferroptotic cell death contributes to PUFA-mediated CD8⁺ T cell loss in the WD-associated HCC microenvironment.

Human HCC samples exhibited significantly elevated ω-6 PUFA abundance that strongly tracked with LA content, consistent with dietary lipid exposure **(Figure 2k)**. Raman scattering further confirmed increased LA and PUFA accumulation in all steatotic HCC samples examined **(Figure 2l; Figure S9h, i)**. In keeping with this, transcriptional programs involved in LA and ω-6 PUFA metabolism were enriched in MASH-related HCC compared to viral– and alcohol-related etiologies **(Figure S2g)**. Targeted plasma lipidomics further revealed that LA-derived metabolites were selectively elevated in MASLD/MASH-HCC compared to both non-MASLD HCC and MASLD without carcinoma, indicating that their accumulation is associated with tumor presence rather than MASLD alone **(Figure 2m)**. Notably, matched peripheral CD8⁺ T cells from MASLD-HCC patients exhibited increased lipid peroxidation, consistent with increased exposure to ω-6 PUFA-derived lipids (**Figure 2n)**. Together, these data identify LA as the principal immunosuppressive component of the WD, which drives tumoral accumulation and transfer of elongated ω-6 PUFAs to CD8⁺ T cells, impairing their function in a cancer-cell non-autonomous manner.

### Targeting FADS2 unleashes antitumor immunity and establishes immunological memory

The rate-limiting step in the LA metabolic cascade is its conversion to GLA by fatty acid desaturase 2 (FADS2). We found elevated FADS2 expression in WD-fed cancer cells relative to both SD-fed counterparts and tumor-adjacent livers **(Figure S9a-d)**. Consistent with enhanced FADS2 expression, hepatic tumors in both MP53 and WD/CCl₄ murine models and 4/5 human MASLD-HCCs exhibited pronounced enrichment of LA-derived PUFA species compared to tumor-adjacent tissue, indicative of active LA flux through the desaturation cascade **(Figure 2c, l; Figure S9e-i)**. Tumor-cell-selective silencing of *Fads2* markedly reduced the release of GLA and downstream ω-6 PUFA species into the TME **(Figure S10a-c)** and rescued the T cell-suppressive effects of WD-derived TIF **(Figure S10d, e)**. Notably, whereas neither *Fads2* deletion nor αPD-1 monotherapy affected tumor progression in the lethal, multifocal MP53 model^51^, their combination extended survival by ∼35% in a manner fully dependent on CD8⁺ T cells (median survival: shCtrl/IgG – 29 d; shCtrl/αPD-1 – 23 d; shFads2/IgG – 29 d; shFads2/αPD-1 – 39 d (*p* < 0.05 v shCtrl/IgG, shCtrl/αPD-2, and shFads2/IgG); shFads2/αPD-1/αCD8 – 31 d (*p* = 0.8 v shFads2/IgG)) **(Figure S10f)**. Next, we tested whether pharmacologic inhibition of this rate-limiting enzyme^52^ could synergize with αPD-1 therapy in our immunogenic model. WD-fed MP53-HCCs were generated via hydrodynamic delivery, and upon tumor establishment, dox was administered to induce lucOS expression. Mice were then treated with either IgG control, the selective FADS2 inhibitor (FADS2i) sc-26196^52^, or their combination. As before, monotherapy with either sc-26196 or αPD-1 alone failed to confer any significant therapeutic benefit (median survival: PBS/IgG – 42 d, FADS2i/IgG – 46 d, PBS/αPD-1 – 37 d). In striking contrast, combined FADS2 inhibition and PD-1 blockade resulted in robust synergy with complete tumor regression observed in ∼25% of treated female (median survival: FADS2i/αPD-1 – 57 d, *p* < 0.005 v PBS/αPD-1) and ∼60% male animals (median survival: IgG/PBS – 56 d, FADS2i/αPD-1 – undefined, *p* = 0.03) **(Figure 3a, b)**. Multiplexed immunofluorescence and spectral profiling of the TME revealed that combination therapy restored an immunologically ‘hot’ state, characterized by increased immune cell infiltration **(Figure 3c-g)**. This included expanded CD45⁺ immune and CD8⁺ T cell populations, with infiltrating T cells exhibiting enhanced activation, as indicated by increased expression of CD25, CD107a, and co-expression of TNFα and granzyme B, reduced expression of the exhaustion marker TOX **(Figure 3c-g; Figure S10g-k)** and antitumor T cell expansion as evident from increased SIINFEKL-targeting TILs **(Figure S10l)**. Furthermore, bulk RNA sequencing of TILs revealed transcriptional signatures consistent with restored and enhanced activation in response to therapy **(Figure 3h, i)**. More importantly, analysis of livers and lymphoid tissue (spleen & lymph nodes) following tumor clearance revealed the presence of SIINFEKL-specific CD8⁺ T cells expressing effector, central, and resident-tissue memory-associated markers, indicating not only durable tumor eradication but also the establishment of long-term immunological memory **(Figure 3j; Figure S10m, n)**. Additionally, most SIINFEKL-targeting CD8⁺ T cells harvested from liver were CD69^+^/CD44^+^ tissue-resident memory cells, while those from the periphery consisted of effector (CD44^high^/CD62L^low^), central (CD44^high^/CD62L^high^), and precursor memory cells (CD127^high^/KLRG1^low^) **(Figure 3j; Figure S10m, n)**. A similar protective effect of FADS2i/αPD-1 combination, driven by CD8⁺ T cell revival, was observed in the immunogenic MPn model **(Figure S11A-d)**. Notably, the survival benefit of FADS2i/αPD-1 combination extended to the orthotopic Hep53.4 model, in which immune evasion is driven by infiltrating myeloid cells and stromal Notch activation^34,53^. While IgG and monotherapy controls succumbed to tumor burden by day 60 (median survival: IgG/DMSO – 44 d; IgG/FADS2i – 43 d; αPD-1/DMSO – 43 d), 1/11 of animals receiving FADS2i/αPD-1 combination remained alive with no observable carcinomas (median survival – 64 d; p < 0.01) **(Figure 3k)**. Similarly, FADS2i/αPD-1 combination treatment improved survival in mice intrasplenically transplanted with an immunotherapy-resistant β-catenin mutant murine HCC cells^37^, suggesting efficacy across mechanistically distinct contexts of checkpoint resistance (median survival: IgG/DMSO – 53 d; FADS2i/αPD – 78 d; p < 0.05) **(Figure S11e)**. To corroborate these findings in human HCC, we interrogated TCGA data, where high *FADS2* expression associated with significantly reduced T cell infiltration by CIBERSORT deconvolution and worse overall survival, independently substantiating a link between LA metabolic activity, immune exclusion, and poor clinical outcome **(Figure S11f, g)**. Reflecting the inverse relationship between FADS2-mediated LA metabolism and cytotoxic T cell infiltration, 6/7 patient samples in MASLD-HCC tissue microarray (TMA) with high CD8^+^ T cell infiltration exhibited low FADS2 expression, while almost all CD8^+^ T cell-depleted TMAs displayed high FADS2 **(Figure 3l; Figure S12)**. Together, these data establish FADS2-driven LA metabolism as a critical driver of immune exclusion in MASLD-HCC and demonstrate that its pharmacologic inhibition synergizes with PD-1 blockade to restore CD8⁺ T cell-mediated tumor control across multiple preclinical models and contexts of checkpoint resistance.

**Figure 3:**
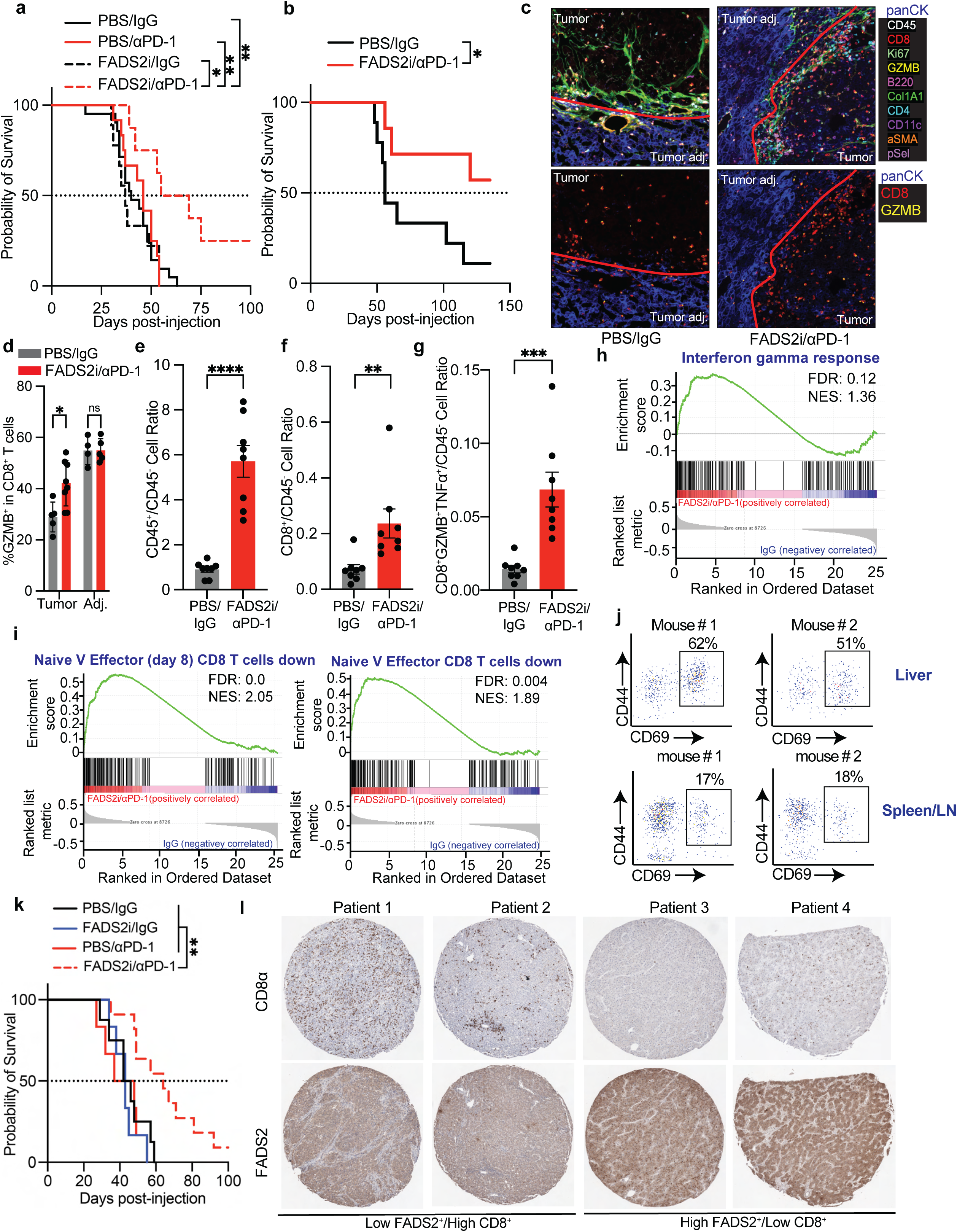
FADS2 inhibition resensitizes WD-driven HCC to immunotherapy. **(a & b)** Kaplan–Meier survival curves of female (a) and male (b) WD-fed mice bearing MP53-lucOS HCCs under indicated treatments (Females: PBS/IgG: n=21, FADS2i/IgG: n=12, PBS/αPD-1: n=9, FADS2i/αPD-1: n=8; Males: PBS/IgG: n=9, FADS2i/αPD-1: n=7); **(c)** Representative multiplexed immunofluorescence images of WD-fed MP53-lucOS tumours treated with PBS/IgG or FADS2i/αPD-1, red line delineates tumour border; **(d)** Frequency of granzyme B⁺ CD8⁺ T cells in tumour and adjacent tissue of WD-fed MP53-lucOS mice under indicated treatments (Tumour: PBS/IgG: n=5 tumours from 3 mice, FADS2i/αPD-1: n=8 from 3 mice; Adjacent: PBS/IgG: n=4 from 2 mice, FADS2i/αPD-1: n=5 from 3 mice); **(e-g)** Relative infiltration of CD45⁺ cells (e), CD8⁺ T cells (f), and GZMB⁺/TNFα⁺ CD8⁺ T cells (g) in WD-fed MP53-lucOS HCCs under indicated treatments (n=8 tumours from 3 mice); **(h & i)** GSEA of tumour-infiltrating CD8⁺ T cells from WD-fed MP53-lucOS treated with either PBS/IgG (n=5) or FADS2i/αPD-1 (n=6); **(j)** Representative dot plots of SIINFEKL-specific tissue-resident memory CD8⁺ T cells in liver (top) and spleen/lymph nodes (bottom) of long-term survivors on FADS2i/αPD-1 treatment; **(k)** Kaplan–Meier survival curves of WD-fed mice orthotopically implanted with Hep53.4 cells and treated as indicated (PBS/IgG: n=8; FADS2i/IgG: n=6; PBS/αPD-1: n=6; FADS2i/αPD-1: n=11); **(l)** Representative HCC tissue microarray stained for CD8α and FADS2 (Columbia University cohort). Results are shown as mean ± SEM. ns: not significant, *p < 0.05, **p < 0.01, ***p < 0.001.

### DJ-1 couples glycolytic flux to immune exclusion by protecting tumoral lipid uptake machinery

Glycation is a non-enzymatic post-translational modification in which reactive dicarbonyl metabolites generated during glycolysis, most prominently methylglyoxal (MGO), covalently modify arginine residues in proteins^21^ **(Figure 4a)**. MGO-derived advanced adducts, including carboxyethyl arginine (CEA) and methylglyoxal hydroimidazolone-1 (MG-H1), accumulate across glycolytic cancers^54^. The extent of glycation scales with glycolytic flux and is opposed by glyoxalase systems, such as Parkinson-associated deglycase 1^23,24^ (DJ-1; encoded by *PARK7*) **(Figure 4a)**. To define functional MGO targets in HCC, we performed chemoproteomic profiling using an alkyne-tagged MGO probe (alkMGO) in HepG2 liver cancer cells **(Figure 4b; Figure S13a)**. Beyond established substrates such as histone H3, this analysis surprisingly revealed a prominent enrichment of proteins involved in lipid uptake and metabolism, including the long-chain fatty acid transporter SLC27A2 (FATP2), the cytosolic fatty acid-binding proteins FABP1 and FABP5, and the lipid droplet-associated protein PLIN3 **(Figure 4c; Figure S13b)**. Among the most extensively glycated species were CYB5B and CYB5R1, which support electron transfer to mitochondrial desaturases and elongases, thereby promoting lipid desaturation and sterol biosynthesis. CLPTM1L, a lipid scramblase implicated in glycosylphosphatidylinositol biosynthesis, was additionally identified as a DJ-1–sensitive glycation target **(Figure S13b; Supplementary Table 2)**. Notably, lipid pathway was not enriched in MCF7 breast cancer cells, consistent with their lower expression in this cell type^50^ **(Figure S13c, d)**.

**Figure 4:**
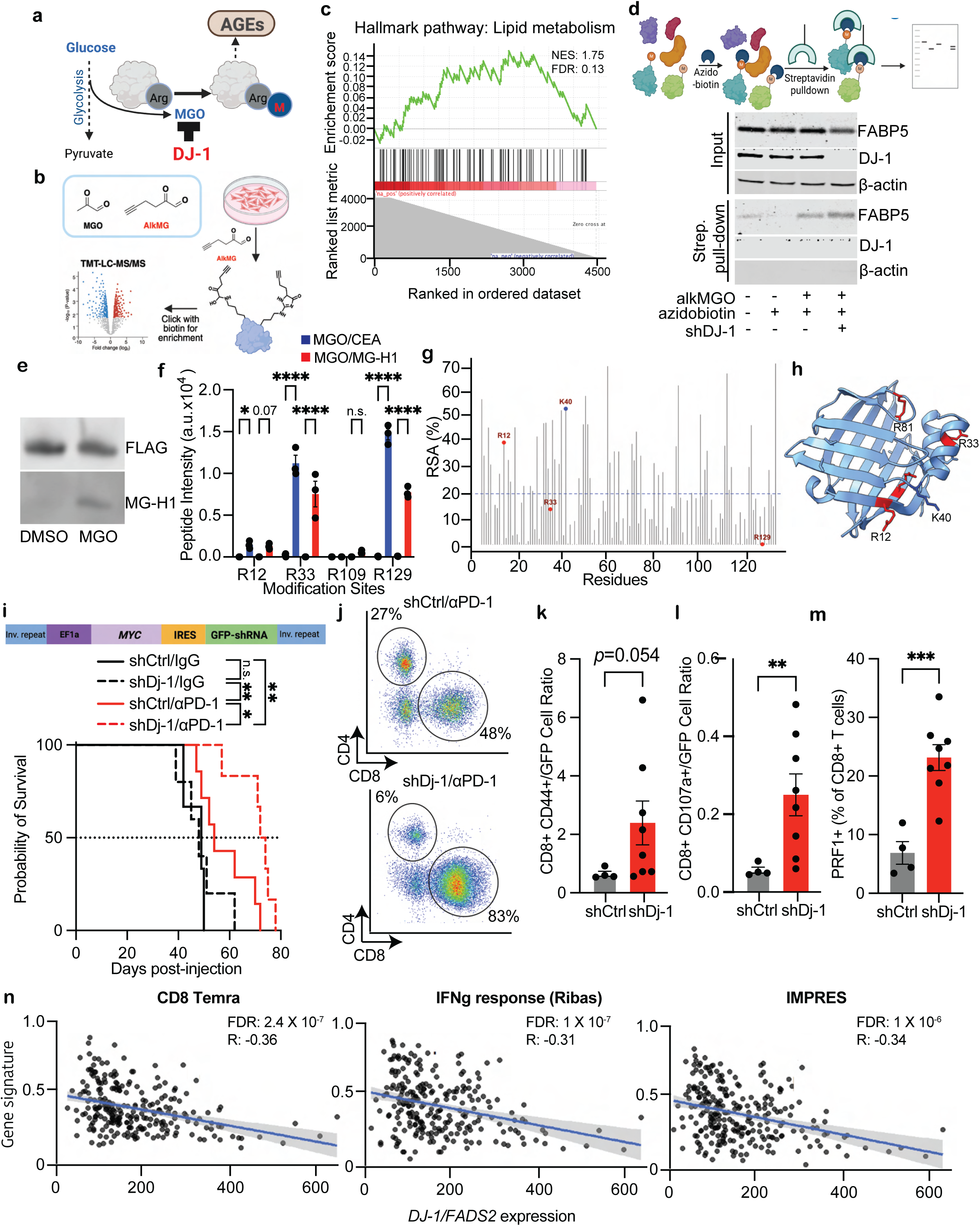
DJ-1-sensitive glycation links LA metabolism to CD8⁺ T cell-mediated antitumor immunity. **(a)** Schematic of MGO-mediated arginine glycation and adduct formation; **(b)** Chemoproteomic workflow for profiling MGO glycation targets using an alkyne-tagged MGO probe; **(c)** GSEA enrichment plot for lipid metabolism proteins among MGO-glycated targets in HepG2 cells; **(d)** Schematic of biotin–streptavidin pull-down strategy (top) and immunoblot of streptavidin-enriched fractions from *DJ-1*–proficient or –deficient HepG2 cells treated with alkMGO (3 mM, 5 h) (bottom); **(e)** Anti-FLAG immunoprecipitates from FABP5-expressing HEK293T cells treated with DMSO or MGO (3 mM, 4 h) probed with MG-H1 antibody; **(f)** Relative peptide intensity of indicated glycation adducts in FABP5-expressing HEK293T cells under DMSO or MGO (3 mM, 4 h) treatment; **(g)** SASA analysis of human FABP5; **(h)** Structural mapping of MGO glycation sites on FABP5; **(i)** Schematic of bicistronic transposon construct (top) and Kaplan–Meier survival curves of WD-fed male mice bearing *Dj-1*–proficient or –deficient MP53 HCCs under indicated treatments (shCtrl/IgG: n=5, shDj-1/IgG: n=5, shCtrl/αPD-1: n=7, shDj-1/αPD-1: n=6) (bottom); **(j – m)** Representative flow cytometry plots of tumour-infiltrating CD4⁺ and CD8⁺ T cells (j), and relative infiltration of CD44⁺ (k), CD107a⁺ (l), and PRF1⁺ (m) cytotoxic T cells in αPD-1–treated WD-fed *Dj-1*–proficient or –deficient MP53 HCCs (shCtrl/αPD-1: n=4 tumours from 2 mice; shDj-1/αPD-1: n=8 tumours from 3 mice); **(n)** Correlation of *DJ-1* and *FADS2* expression with CD8⁺ T cell cytotoxicity– and immunotherapy-response predictive gene signatures. Results are shown as mean ± SEM. ns: not significant, *p < 0.05, **p < 0.01, ***p < 0.001, ****p < 0.0001.

Lipid-related DJ-1 clients are involved in LA metabolism; FATP2 facilitates cellular LA import, while FABP1 and FABP5 direct intracellular LA trafficking to the endoplasmic reticulum for FADS2-mediated desaturation into longer-chain ω-6 PUFAs and signaling via activation of the mechanistic target of rapamycin complex I (mTORC1)^55^. To establish these proteins as *bona* fide DJ-1 substrates, we compared alkMGO labeling in *DJ-1*-proficient and *DJ-1*-deficient HepG2 cells using azido-biotin conjugation and streptavidin enrichment **(Figure 4d)**. FABP1 and FABP5 were selectively enriched in DJ-1–deficient pull-downs despite reduced total protein abundance, indicating enhanced glycation and impaired adduct repair in the absence of DJ-1 **(Figure 4d; Figure S13e)**. To validate intracellular MGO adduction sites, FLAG-tagged *FABP5* was expressed in HepG2 cells and treated with MGO (3 mM, 4 h). FLAG immunoprecipitation followed by MG-H1 immunoblotting confirmed MGO-dependent glycation of FABP5 **(Figure 4e)**, and subsequent mass spectrometry identified R12, R33, and R129 as the major MGO-susceptible arginine residues, bearing both CEA and MG-H1 adducts **(Figure 4f; Figure S14-16)**. R109 emerged as a minor site carrying MG-H1 modifications exclusively, while K40 was inconsistently detected across replicates and likely represents a low-occupancy site **(Figure 4f; Figure S14c)**. Solvent-accessible surface area (SASA) analysis^56,57^ confirmed the full surface accessibility of R12 and K40 and partial accessibility for R33 **(Figure 4g, h)**. Notably, despite its burial within a β-sheet, R129 **(Figure 4h)** was identified as a glycation-prone residue both in this study and independently in a recent chemoproteomics screen employing a more reactive phenylglyoxal –based probe^56^, supporting its intrinsic “glycatibility”. In line with this, *DJ-1* deficiency reduced steady-state and lipid-induced levels of FABP1, FABP5, and FATP2, with cycloheximide chase confirming accelerated client protein degradation **(Figure S13f, g)**. *DJ-1* loss additionally reduced cell surface CD36 levels, though CD36 was not itself identified as a chemoproteomic MGO target, suggesting that DJ-1-dependent regulation of the lipid uptake machinery extends beyond detected substrates **(Figure S13h)**. Consistent with DJ-1-dependent maintenance of LA import machinery, *DJ-1/Dj-1* loss suppressed LA uptake both *in vitro* **(Figure S13i)** and in MP53 tumors *in vivo* **(Figure S13j)**; an effect recapitulated by MGO treatment alone and rescued by concurrent DJ-1 overexpression **(Figure S13k)**. To determine whether impaired LA uptake translates to reduced PUFA transfer to CD8⁺ T cells, we pulsed *Dj-1*-proficient and *Dj-1*-deficient murine HCC cells with an alkyne-tagged LA probe and collected conditioned medium for transfer to CD8⁺ T cells. Consistent with reduced LA uptake, DJ-1-deficient cells exhibited markedly diminished secretion of PUFA-containing lipid species, as evidenced by reduced alkyne-lipid transfer to CD8⁺ T cells **(Figure S13l)**. Taken together, these data identify DJ-1 as a critical regulator of LA metabolism in HCC, maintaining the stability of key lipid uptake and trafficking proteins through prevention of MGO adductions, thereby sustaining LA import, intracellular trafficking, and ultimate transfer of immunosuppressive ω-6 PUFAs to CD8⁺ T cells.

These findings establish that DJ-1 controls detoxification of glycolysis-derived MGO and stabilizes LA-handling proteins, raising the question of whether DJ-1 loss phenocopies pharmacological FADS2 inhibition in suppressing PUFA-mediated CD8⁺ T cell immunosuppression. To address this, we examined DJ-1 expression and MGO adduct burden *in vivo*. WD-fed MP53 and MPn HCCs exhibited elevated glucose uptake and DJ-1 expression **(Figure S17a-e)**, and selective tumoral *Dj-1* loss via transposon-encoded shRNA **(Figure S17l)** (schematic shown in Figure 4i) increased MGO levels and MGO-derived glycation adducts, most prominently MG-H1 and, to a lesser degree, CEA, without affecting lysine-directed glycation **(Figure S17f, g)**. Consistent with glycation-driven client protein destabilization, WD feeding elevated expression of several lipid handling proteins in *Dj-1*-proficient MP53 HCCs (e.g., FABP1, FATP2, CD36), and these proteins were selectively reduced upon *Dj-1* silencing **(Figure S17h, i)**. *Dj-1* loss did not impact cellular redox state, and GLO1 protein levels were unchanged **(Figure S17j, k)**, excluding compensatory upregulation of the canonical glyoxalase detoxification system^58^. Consistent with uncompensated MGO accumulation, targeted ω-6 lipidomics of WD-fed *Dj-1*-silenced MP53 tumors revealed a selective and marked reduction in LA-derived PUFA species relative to *Dj-1*-sufficient counterparts, while untargeted tumoral lipidomics showed no major global alterations in lipid composition, indicating that DJ-1 loss impairs substrate flux specifically through the LA desaturation cascade rather than broadly disrupting lipid homeostasis **(Figure S17m, n)**. Selective tumoral *Dj-1* loss did not affect tumor growth as a monotherapy, mirroring the phenotype observed with FADS2 inhibition, yet, it conferred a significant survival benefit in combination with anti-PD-1 under Western diet conditions (Median survival: Males: shCtrl/IgG – 49 d, shDj-1/IgG – 48 d, shCtrl/αPD-1 – 54 d, shDj-1/αPD-1 – 73 d) **(Figure 4i; Figure S18a)**. Notably, this protection is fully negated by co-depleting CD8^+^ T cells **(Figure S18a)**. Immune phenotyping of the TME confirmed a shift toward an immunologically hot milieu, with increased T cell ingress and activation markers consistent with productive anti-tumor immunity **(Figure 4j-m; Figure S18c-i)**. This synergy was absent in standard diet-fed animals **(Figure S18b)**, consistent with a metabolic dependency in which elevated glycolytic flux and lipid substrate availability are required for DJ-1 to constitute a rate-limiting determinant of LA uptake and tumoral PUFA production. Corroborating these findings clinically, TCGA analysis revealed a significant inverse correlation between *DJ-1* mRNA expression and T cell infiltration, and high *DJ-1* expression associated with worse overall survival, recapitulating the prognostic relationship observed for FADS2 **(Figure S19a-c)**. TCGA analysis further revealed that *DJ-1* and *FADS2* upregulation are largely mutually exclusive across HCC tumors **(Figure S19d)**. Consistent with these observations, both *DJ-1* and *FADS2* expression showed a significant but modest inverse association with the IMPRES immunotherapy response signature^59^ and T cell activation score^60^; however, their combined expression strengthened these inverse associations **(Figure 4n; Figure S19e, f)**. These findings establish DJ-1 as a diet-dependent, rate-limiting regulator of tumoral LA metabolism whose loss, like FADS2 inhibition, selectively disrupts ω-6 PUFA production and synergizes with PD-1 blockade to restore CD8⁺ T cell-mediated antitumor immunity.

### LA metabolic axis defines immune-excluded HCC and predicts resistance to checkpoint therapy

To assess the clinical relevance of these findings, we interrogated transcriptomic data (n=247) from the phase 1b (GO30140) and phase III (IMbrave150) trials evaluating atezolizumab plus bevacizumab in advanced HCC^61–63^. This analysis revealed significant enrichment of lipid activation, elongation, and metabolism-related pathways along with lipid and linoleic acid-related signatures in non-responders **(Figure 5a)**. Strikingly, lipid and LA-related metabolic signatures were specifically enriched in partial and non-responder patients **(Figure 5b)**. This enrichment was particularly evident in non-inflamed tumors, specifically those with an immune-excluded phenotype **(Figure 5c),** as predicted by our causation studies in mice. These results implicate LA metabolic reprogramming as a driver of immunosuppressive TME remodeling and resistance to checkpoint-based therapy in HCC.

**Figure 5:**
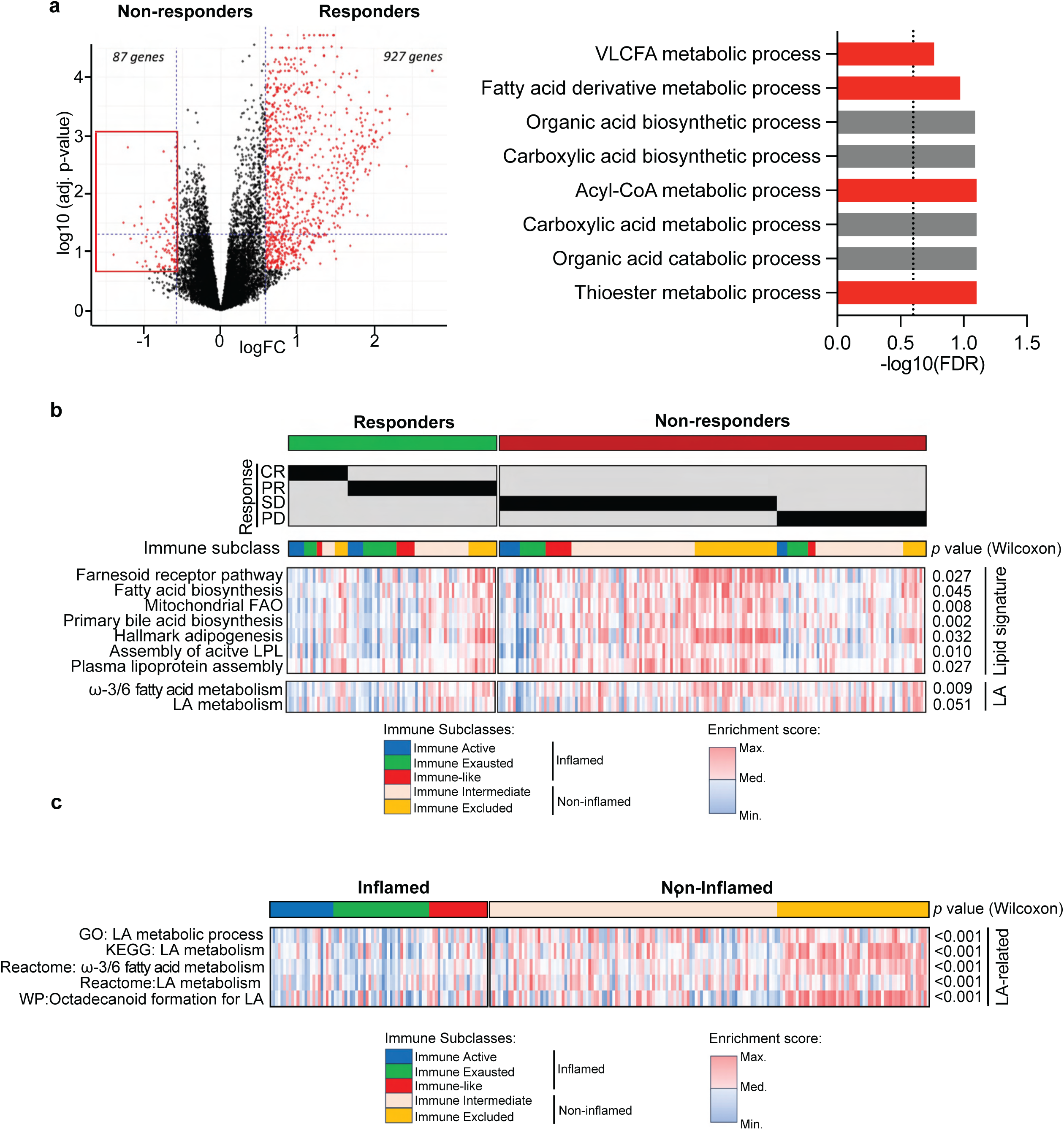
LA-metabolic pathways correlate with poor response to immunotherapy in advanced HCC. **(a)** Volcano plot (left) and GSEA (b) of differentially expressed genes between non-responders (n=166) and responders (n=81) to atezolizumab plus bevacizumab in HCC, significantly enriched genes in non-responders highlighted in red; **(b)** Heatmap comparing lipid– and LA-related pathway enrichment between responders and non-responders to atezolizumab plus bevacizumab treatment; p-values calculated by Wilcoxon signed-rank test; **(c)** Heatmap showing enrichment of LA-related pathways in the immune-excluded tumour phenotype in atezolizumab plus bevacizumab-treated HCC patients.

Collectively, these data establish that direct lipid desaturation and glycolytic MGO accumulation represent convergent but mechanistically distinct routes to LA-mediated immune exclusion, and identify FADS2 and DJ-1 as therapeutically actionable targets for restoring immunotherapy responsiveness in WD-associated HCC **(Figure 6)**.

**Figure 6:**
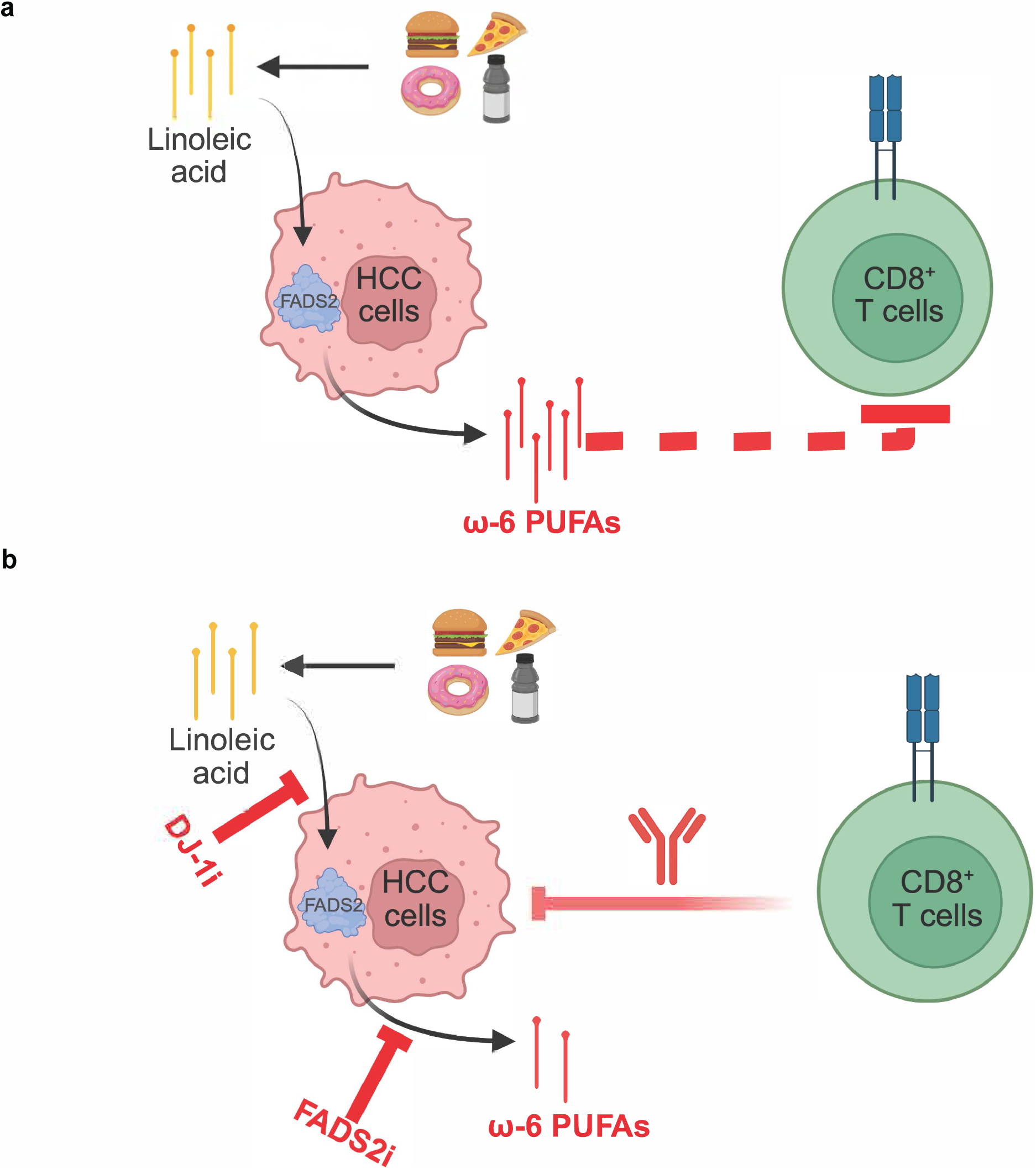
FADS2 and DJ-1 drive LA-mediated immune evasion in WD-associated HCC. **(a & b)** Proposed mechanism by which FADS2 and DJ-1 cooperate to drive LA-mediated immune suppression in WD-associated HCC and its therapeutic reversal by FADS2 or DJ-1 inhibition.

## Discussion

The rising incidence of HCC linked to WD overconsumption and MASLD underscores the urgent need to define how nutrient overload shapes tumor metabolism and immune evasion. The liver is inherently immunosuppressive, a feature further compounded by overnutrition, which likely underlies the repeated failure of immunotherapies in HCC^64,65^. As the mechanisms underlying immunotherapy resistance in MASLD-associated HCC have remained poorly understood, druggable targets that may improve ICI-based therapies in this relevant disease context have not yet been identified. Although previous studies have pointed towards MASLD-associated CD8⁺ T cell dysfunction driven by lipid deposition and metabolic disruption^9,12^, the identity of specific immunosuppressive dietary metabolites remains elusive^14,15^. Here, we address this gap by developing a conditionally immunogenic autochthonous mouse model that integrates oncogenic driver expression, dietary overnutrition, and temporal neoantigen activation — enabling, for the first time, direct interrogation of how diet-driven metabolic reprogramming shapes antitumor immunity in an anatomically relevant context. Using this system, we identify a paracrine and druggable dietary lipid axis in which tumor-intrinsic LA metabolism generates an immunosuppressive microenvironment that constrains CD8⁺ T cell antitumor function and contributes to immunotherapy resistance.

Several recent studies have examined the effects of dietary sugars and lipids on T cell activity, demonstrating highly context-specific outcomes that vary with tumor type, anatomical site, and experimental model^15,17^. Liver-related studies have relied on prolonged dietary regimens, suggesting that observed T cell effects arise primarily from chronic nutrient exposure^5,6^. Our HDTV-based models, which disentangle tumor-specific effects from those of chronic WD exposure, reveal an immediate and dominant impairment of CD8⁺ T cells following WD exposure — effects independent of the systemic consequences of chronic dietary exposure and instead reflecting tumor-specific responses to overnutrition. Importantly, despite the presence of strong neoantigens, even short-term WD exposure completely abrogates the T cell response, mirroring the failure of immunotherapy observed in our mouse models and human MASLD-HCC patients^9^. Unlike in melanoma, where T cell dysfunction stems from lipid deprivation^15^, in the hepatic setting, it is not the loss of lipid nutrients but rather tumor-derived secondary oncometabolites generated from LA that drive immune suppression — a distinction that reframes our understanding of how diet shapes antitumor immunity in the context of metabolic liver disease.

Linoleic acid is the most abundant fatty acid in Western diets, constituting up to 90% of total ω-6 PUFA intake driven largely by widespread consumption of LA-rich vegetable oils^66–68^. While LA is canonically recognized as a precursor for pro-inflammatory eicosanoids, our findings identify a mechanistically distinct immunosuppressive role for its longer-chain ω-6 PUFA derivatives, produced exclusively within tumor cells via FADS2, that operates through direct suppression of CD8⁺ T cell function. Previous studies have shown that while *in vitro* LA exposure enhanced CD8⁺ T cell activation^69^, chronic consumption of an LA-rich diet *in vivo* impairs it^70^ — a paradox resolved by our demonstration that LA is exclusively consumed and metabolized by cancer cells, shielded from immune cells, and converted to a spectrum of longer-chain ω-6 PUFAs that potently suppress CD8⁺ T cells in a chain length– and desaturation-dependent manner. Minimal inhibition is seen with shorter-chain species such as GLA (C18:3), whereas maximal suppression occurs with highly unsaturated long-chain species such as DPA (C22:5). These lipids are detected in the TIF of WD-fed murine HCC models and in MASLD-HCC patients^71,72^, and are likely internalized by T cells, likely via CD36^73^ or passive diffusion, where they increase lipid peroxidation and drive mitochondrial depolarization to additively suppress CD8⁺ T cell function.

Warburg tumors generate toxic MGO that must be neutralized^27,74–76^. DJ-1 is canonically recognized as a glyoxalase that protects histones^27^ and nucleic acids^77^. Our findings reveal an unrecognized role in regulating hepatocellular lipid uptake and immune evasion under metabolic stress. Chemoproteomic profiling revealed that DJ-1 prevents MGO adduction of a network of lipid importers and cytoplasmic chaperones — including FATP2, FABP1, FABP5, and the desaturase cofactors CYB5B and CYB5R1 — positioning it as a central node enabling tumor cells to preferentially acquire and metabolize immunosuppressive lipids in a nutrient-competitive TME. Among these substrates, FABP5 is of particular interest — a cytoplasmic chaperone known to shuttle LA to the endoplasmic reticulum and activate mTORC1 signaling^55,78^. Structural analysis of the FABP5 binding pocket identifies R109 and R129 as key LA-coordinating residues, with R129 forming a salt bridge with the carboxyl head group of LA and R109 engaging LA via a hydrogen bond through an ordered water molecule^78^.

From a therapeutic standpoint DJ-1 or FADS2 inhibition synergized with anti-PD-1 to markedly prolong survival by counteracting the oncogenic effects of LA metabolism in an immunogenic context. Notably, neither DJ-1 nor FADS2 inhibition alone affected tumor growth, akin to anti-PD-1 monotherapy, highlighting the multifactorial nature of WD-induced T cell blockade. Although both FADS2^51^ and DJ-1^27^ have been implicated as anticancer targets in other metabolic settings, our study uniquely identifies their cooperative role in a linoleic acid–driven axis that links tumor lipid metabolism to CD8⁺ T cell suppression and immunotherapy resistance. Beyond their effect on tumor immune suppression, these inhibitors, especially DJ-1, may also have application in other metabolic disorders that are caused by excessive lipid uptake and energy imbalance, such as obesity, cardiovascular diseases, and MASLD^79^.

While our findings support the combination of LA-metabolism inhibitors with immunotherapy in MASLD/overnutrition-driven HCCs, they also raise the possibility that diet–immunotherapy interactions broadly influence treatment outcomes across MASLD-dependent and –independent cancers, potentially contributing to patient-to-patient variability in response. Finally, dietary intervention-based adjuvant cancer therapies, such as intermittent fasting, caloric restriction, ketogenic diets, and plant-based or other specially formulated regimens are gaining popularity, yet their efficacy remains highly variable, influenced by tumor stage, anatomical location, driver mutations, sex, and other environmental factors^80,81^. Our findings further support that restricting dietary LA can be an effective strategy to restore T cell function and enhance responses to immunotherapy.

## Author contributions

N.C., R.K. and J.S.K. designed and performed experiments, analyzed data, and edited the manuscript. U.B., G.C.S and A.V.E performed the bioinformatic analysis on human HCC data. M.I. performed the Raman spectroscopy experiments. J.M. and J.R. performed the PBMC lipid peroxidation staining experiments. S.F. designed, performed, and analyzed multiplexed immunofluorescence. Y.X. and K.N. designed, performed, and analyzed the MGO-alkyne chemoproteomics experiments. A.M.*P*. performed MGO/AGE quantification. R.N. and M.S. helped with animal experiments involving surgery. D.P.G., L.A.A., J.J., M.S. and R.L. performed lipidomics experiments. J.O., M.R., M.P., J.S. and E.P. provided experimental assistance. B.C. and X.W. helped with histopathological analyses of murine and human samples, respectively. S.T. and E.B. helped with scRNAseq data acquisition and analyses. R.N. supervised metabolomic experiments. M.M. supervised proteomics and metabolomics experiments. D.L. supervised the targeted lipidomic experiments and provided critical reagents. D.B. and D.C.W. supervised and provided critical reagents for the patient PBMCs staining experiments. M.R.G. supervised the Raman spectroscopy experiments and provided critical reagents. B.R.S. provided critical reagents for targeted lipidomic experiments. R.F.S. supervised animal experiments, contributed with critical reagents, human data analysis, and assisted with manuscript editing. J.J.G. supervised MGO/AGE quantification. *P*.D.J. provided access to patient PBMCs and serum samples. *P*.B.R. provided access to HCC patient samples, provided critical reagents, and supervised the multiplexed immunofluorescence experiments. Y.D. provided critical reagents and supervised chemoproteomics experiments. R.P. supervised patient HCC bioinformatic analysis. J.M.L. supervised patient HCC bioinformatic data interpretation. V.R.S. conceived and supervised the study and wrote the manuscript. All authors read and approved the manuscript.

## Acknowledgments

We acknowledge Drs. Christine Chio (Columbia University), Scott Friedman (Mt. Sinai), Anil K. Rustgi (Columbia University), and Priyamvada Rai (University of Miami) for providing reagents, protocols, and/or experimental discussion. We thank Peter A. Sims for assistance with scRNAseq bioinformatic analysis. We thank Caleb Porter, Arsalan Hashmi, and Zhuoning Li (MSKCC) for their assistance with proteomics and metabolomics experiments. We sincerely thank members of the following core facilities at Columbia University: Oncology Precision Therapeutics and Imaging Core, Molecular Pathology, and Flow Cytometry Core by the Herbert Irving Comprehensive Cancer Center (HICCC); Bioinformatics and Single Cell Analysis Core by Digestive and Liver Diseases Research Center; and Flow Cytometry Core by Columbia Stem Cell Initiative. We thank the members of the Institute of Comparative Medicine for animal monitoring and husbandry. N.C. was supported by a Fulbright scholarship, a postgraduate fellowship from La Caixa Foundation, a Trainee Associate Member Predoctoral Award from HICCC, and internal funds from Sylvester Comprehensive Cancer Center at the University of Miami. J.O. was supported by the Trainee Associate Member Postdoctoral Award from HICCC. G.C.S was supported by Ministerio de Ciencia, Innovación y Universidades with a FPU fellowship (FPU24/01471). A.V.E was supported by “La Caixa” Foundation (ID 100010434), under grant code LCF/BQ/DFR25/12000106. L.A.A. is an American Cancer Society Post-Baccalaureate fellow and was supported in part by the University of Miami, Miller School of Medicine, and the Sylvester ACS Post-Baccalaureate Program award DICR POST-BACC-23-1161831. B.C. was supported by São Paulo Research Foundation (FAPESP; 2022/02175-1). R.P. was supported by the Fundació de Recerca Clínic Barcelona – IDIBAPS and by a grant from the Spanish National Health Institute (MICINN, PID2022-139365OB-I00, funded by MICIU/AEI/10.13039/501100011033 and FEDER). D.B.L. is supported by the following grants: R01CA253986, R33AG077856, I01BX006593. This work is also supported by V Scholar Award V2024-033 (D.B.), NIND 1R35GM162083-01 (D.B.), and Sylvester Comprehensive Cancer Center (SCCC) Start-up Funds (D.B., D.C.W.). B.R.S. is supported by the National Cancer Institute (P01CA291697) and a Cancer Center Support Grant (P30CA008748). J.J.G. is funded by NIH/NIGMS R35 GM137910 and NIH/NIDDK R01 DK133196. *P*.B.R is supported by the Memorial Sloan Kettering Cancer Center (MSK) Support Grant [P30 CA008748], an NIH/NCI grant [K08 CA255574] and a NIH/NCI grant (R37 CA304010). Y.D. is supported by the Josie Robertson Foundation, the Pershing Square Sohn Cancer Research Alliance, the NIH (CCSG core grant P30 CA008748, MSK SPORE P50 CA19HEK293T7, and R21 DA044767), the Parker Institute for Cancer Immunotherapy (PICI), and the Anna Fuller Trust. In addition, the Y.D. lab is supported by Mr. William H. Goodwin and Mrs. Alice Goodwin, the Commonwealth Foundation for Cancer Research, and the Center for Experimental Therapeutics at Memorial Sloan Kettering Cancer Center. M.R.G. is funded by an MIRA grant (R35GM150564) from NIH/NIGMS. M.R.G and M.I. were partially supported by the National Science Foundation (NSF CAREER award number: 2045640). M.R.G. acknowledges LSU Advanced Microscopy and Analytical Core (AMAC) facility for the Raman imaging. J.M.L. was supported by grants from the European Commission (Horizon Europe-Mission Cancer, THRIVE, Ref. 101136622), the NIH (R01-CA273932-01, R01DK56621 and R01DK128289); the Samuel Waxman Cancer Research Foundation; the Spanish National Health Institute (MICINN, PID2022-139365OB-I00, funded by MICIU/AEI/10.13039/501100011033 and FEDER); by an Accelerator Award from Cancer Research UK, Fondazione per la Ricerca sul Cancro (AIRC) and Fundación Científica de la Asociación Española Contra el Cáncer (FAECC) (HUNTER, Ref. C9380/A26813); by grants from the Fundación Científica de la Asociación Española Contra el Cáncer (FAECC)(Proyectos Generales, Ref. PRYGN223117LLOV; Reto AECC 70% Supervivencia, Ref. RETOS245779LLOV and AECC-IDIBAPS Excellence Program, Ref. EPAEC246711CLIN), the Generalitat de Catalunya/AGAUR (2021 SGR 01347) and “la Caixa” Foundation under the agreement LCF/PR/SP23/52950009, Cancer Research Group Support Program (IDIBAPS). V.R.S is a recipient of the Department of Defense’s Concept Award (W81XWH2010549) and Career Development Award (W81XWH2110377), Rally Foundation’s Independent Scientist Award (20IN31), Glaser Foundation’s New Investigator Research Award (UM SJG 2022-24). V.R.S is also supported by American Cancer Society’s Research Scholar Grant (RSG-22-071-01-TBE) and MIRA (R35GM147497) from NIH/NIGMS. Additionally, V.R.S was supported by faculty development start-up package by the University of Miami and Sylvester Comprehensive Cancer Center and is supported by faculty development funds from the HICCC. Research reported in this publication was performed in part at the Flow Cytometry Shared Resource (FCSR; RRID: SCR_022501) of the SCCC at the University of Miami Miller School of Medicine, which is supported by the National Cancer Institute Cancer Center Support Grant (CCSG) P30-CA240139. Lipidomics from D.B.L lab (Figure 2m) was supported by Sylvester Comprehensive Cancer Center. Part of the lipidomics experiments was supported by the Core grant P30 CA008748 and the Proteomics and Metabolomics Core, RRDI: SCR_027811. This research was funded in part through the NIH/NCI Cancer Center Support Grant P30CA013696 and Columbia University Digestive and Liver Disease Research Center (CU-DLDRC) grant P30DK132710, and using the CU-DLDRC BSCAC, BIC, and OCCC cores.

## Methods and Materials

### Statistical analysis

Graphs and statistical analyses were performed using GraphPad Prism unless otherwise stated. For all experiments, n indicates the number of independent biological replicates. Specifically, for survival curves, n represents the number of mice; for flow cytometry analysis of immune infiltrates, n represents the number of tumors harvested from at least two independent mice, except for Figure 1l-n and Figure S3d-f,h,j, where one mouse was used. For MP53 and temporal scRNA-seq, each experimental group consisted of two independent mice, with 3–4 tumor nodules pooled per group, except for MP53-lucOS WD day 20, where four mice were used. For *in vitro* CD8⁺ T cell and HepG2 experiments, n represents independent wells, repeated in at least two independent experiments. Two-tailed unpaired Student’s t-tests were applied to Figures 1j-l, 4c-g, and Figure S3a,f,h, 4a-c,j,m, 5f,g, 6a,d, and 8b,c. Survival curve differences (Figures 1b,c,h,m 3b, 4a,b,i and Figure S1e,f, 3b,c, 8a, 9a,d, 12a were assessed using the log-rank (Mantel–Cox) test. Multiple t-tests with Holm–Sidak correction were applied to Figure 2c and Figure S3d,e,j, 5e, and 9b. One-way ANOVA followed by Sidak’s multiple comparison test was applied to Figures 1n, 2f, 4j, and Figure S4k,n, 5a–c, 7a–e, 8d,e, and 13c. Linear regression was used to analyze Figures 1g, 3e and Figure S2g. For Figure S13a, mutual exclusivity was assessed in cBioPortal using Fisher’s exact test. Randomization was used to allocate mice to diet and treatments. For experiments shown as representative, results were reproduced in at least two independent experiments with consistent outcomes.

### Animal studies

All animal procedures were approved by the Columbia University Institutional Animal Care and Use Committee (protocol #AABV8661). Mice were housed in the Columbia University Russ Berrie Medical Science Pavilion, under a 12 h light-dark cycle, constant temperature of 21-24 °C, and 50-60% of humidity. Animals were fed with *ad libitum* water and regular mouse chow (PicoLab Rodent Diet 20, #5053) during breeding and non-experimental maintenance. For experimental procedures, mice under Standard Diet (SD) were fed *ad libitum* with #D12450K (Research Diets, Inc., 10% of total kcal from fat, and 4.7% from LA) and regular water, while Western Diet (WD) was defined as #D12492 (Research Diets, Inc.; 60% of total kcal from fat, and 18.6% from LA) supplemented with high-fructose/high-glucose water (23.1 g/L D-fructose + 18.9 g/L D-glucose). High-fat diet (HFD) consisted of WD without high-fructose/high-glucose water supplementation, whereas high-sugar diet (HSD) consisted of SD supplemented with high-fructose/high-glucose water. For the LA-deficient diet, #TD.88137 (Envigo, 42% of total kcal from fat, and less than 1% from LA) was used. LA-enriched WD was a custom-calorie– and nutrient-matched WD in which fat components were modified to make LA the primary source of fat-derived calories (#D25122401; Research Diets, Inc.; 60% of total kcal from fat, and 41% from LA). The doxycycline versions of the same diets had 625 mg/kg of doxycycline hydrochloride added. 8–10-week-old C57BL/6J (JAX, #000664), NSG (JAX, #005557), and DJ1^-/-^ (JAX, #006577) male and female mice were obtained from The Jackson Laboratories.

Hydrodynamic tail vein injections were performed as described previously^31,32,82^. To generate *MYC*/sg-p53 (MP53) and *MYC*/sg-PTEN (MPn) HCC tumors, 8-10 week-old C57BL/6 mice were injected into the lateral tail vein with a plasmid mix consisting of pT3-EF1α-*MYC* transposon backbone (10 μg), Sleeping Beauty transposase SB13 (2 μg), and CRISPR/Cas9-containing PX330 plasmid expressing a single guide RNA against *Trp53* (sg-p53) or *Pten* (sg-PTEN) (10 μg) diluted in 2 mL of sterile 0.9% NaCl saline per mouse. For experiments involving IVIS imaging, the pT3-EF1α-*MYC*-IRES-Luciferase plasmid was used. For the constitutive expression of small hairpin RNAs (shRNAs), Dox-inducible luc/lucOS, and constitutive expression of luc/lucOS experiments, 10 μg of EF1α-*MYC*-IRES-GFP-shRNAs, EF1α-*MYC*-rtTA3-TRE-luc/lucOS, or EF1α-*MYC*-IRES-luc/lucOS transposon plasmids were used, respectively. The targeting sequences for the shRNAs were: *Dj-1* (5’– GGGTGCACAGAATTTATCTGAT-3’), *Fads2 (*5’-ACCCTGGTTTTCCTCAACTTTAT-3’), and Luciferase-targeting used as shCtrl (5’-AAGGTATATTGCTGTTGACAG-3’). Plasmid vectors were isolated using the QIAGEN PlusMaxi kit following the manufacturer’s instructions and confirmed via sequencing. Animals were randomized 5 days post-injection for diet assignment and switched from regular chow to the appropriate experimental diet. For Dox-inducible luc/lucOS experiments, animals were switched to the Dox-containing equivalent of the diets on day 18-21 post-injection. For the constitutive expression of the lucOS experiment (Figure S5e), mice were pre-fed for 8 weeks in advance of HDTV. Tumor initiation was confirmed by liver palpation and ultrasound imaging before dox-containing diet switch or treatment initiation. Positron emission tomography (PET) scans were acquired using an Inveon microPET scanner (Siemens, Germany) according to standard procedures, and images were analyzed using VivoQuant version 4 (Invicro, MA, USA). For bioluminescence imaging, the abdominal area was first depilated using Nair hair removal lotion. Mice were administered D-luciferin (150 mg/kg, intraperitoneally) and anesthetized with 2% isoflurane for 10 minutes. Images were acquired using the Ami HTX optical imaging system (Spectral Instruments Imaging), and signal intensity was quantified as total photon flux (photons/sec) within regions of interest (ROIs) encompassing the liver area using Aura Imaging Software (Spectral Instruments Imaging).

#### Liver electroporation

Mice were anesthetized with 2% isoflurane in an induction chamber, and the abdominal area was shaved and disinfected with 70% ethanol. Following laparotomy, the left liver lobe was pulled out and injected with a 50 µL plasmid mixture containing pT3-EF1α-*MYC*-IRES-Luciferase (15 µg), SB13 transposase (3 µg), and PX330-sg-p53 (10 µg) diluted in sterile 0.9% NaCl. Electroporation was performed using tweezer electrodes connected to an *in vivo* electroporator (NEPA21 Type II Electroporator, Nepa Gene) by applying two pulses of 70 V for 35 ms at 500 ms intervals around the injection site. After electroporation, the peritoneal cavity was sutured, and the skin was closed with surgical staples. Mice were maintained at 37°C during recovery until fully awake. Postoperative analgesia was provided with buprenorphine and/or meloxicam for 3 consecutive days following surgery. Tumor progression was monitored longitudinally by bioluminescence imaging.

#### Orthotopic liver injection

8-week-old C57BL/6 mice were pre-fed in WD for 2 weeks and orthotopically injected in the liver with a total of 0.6 × 10^6^ Hep53.4 syngeneic cells in 50 μl of PBS per mouse, previously described as an αPD-1-resistant MASH-HCC model.

#### Liver metastasis by intrasplenic injection

Mice were pre-fed with WD for two weeks *ad libitum*, anesthetized with 2% isoflurane, and a laparotomy was performed in the left abdominal flank. A total of 0.5 × 10^6^ cells of the immunotherapy-resistant *MYC*/sg-p53/β-catenin^Δ90^ mHCC cell line were injected into the spleen in 100 μl of PBS. After 90 s, partial splenectomy was performed.

#### Antibody and drug treatments

For depletion and blockade antibody treatments, mice were injected intraperitoneally twice a week with isotype control IgG2b (IgG, clone LTF-2, BioXCell), anti-PD-1 (αPD-1, clone 29F.1A12, BioXCell) or anti-CD8a (αCD8a, clone 2.43, BioXCell). Animals were treated with 3 initial doses of 200 µg/mouse and followed with 100 µg/mouse injections during the rest of the experiment. FADS2 inhibitor (sc-26196^52^) was dissolved in a DMSO:PEG4000:saline (1:7:12) solution, vortexed at 50 °C for 10 minutes, and delivered via oral gavage (50 mg/kg, 3 times/week). For *in vivo* LA uptake experiments, mice were fasted for 4 h and injected intravenously (i.v.) with 20 µg/mouse of LA-Bodipy at a concentration of 0.1 mg/mL diluted in 5% BSA. After 45 minutes, tumors were harvested and processed for flow cytometry analysis.

#### Tissue harvest and processing

Mice were euthanized by CO₂ asphyxiation following early tumor development or at humane endpoint. Tumor nodules were harvested and either stored in RNAlater for subsequent RNA analysis or flash-frozen, while whole liver tissues were fixed in 10% neutral buffered formalin. For TIF collection, 3–4 tumor nodules were excised, rinsed in PBS, blotted dry, and placed in a 40-µm cell strainer in a 50-mL Falcon tube. The samples were centrifuged at 110 × g for 5 min, and the flowthrough supernatant was collected and stored at –80 °C. For plasma collection, blood was obtained by cardiac puncture and transferred to a tube containing 10 µL of 0.5 M EDTA, followed by incubation on ice for 15 min and centrifugation at 500 × g for 10 min at 4 °C. The resulting plasma supernatant was collected and stored at –80 °C.

#### Mouse liver histology and scoring

Hematoxylin and eosin (H&E) and Sirius red staining were performed on formalin-fixed, paraffin-embedded (FFPE) sections. After fixation, liver samples were paraffin-embedded, sectioned at 5 µm, and mounted on charged slides. Following deparaffinization and rehydration, sections were stained and evaluated by a veterinary pathologist blinded to group allocation. For the tumor-adjacent liver tissue, lesions were graded according to the rodent MASLD scoring framework^83^. For each animal, an overall steatosis score was calculated by adding the scores for macrovesicular steatosis, microvesicular steatosis, and hepatocellular hypertrophy. Each parameter was graded from 0 to 3 according to the percentage of affected parenchyma: <5%, 5–33%, 33–66%, or >66%, respectively. Lobular inflammation was assessed by counting inflammatory foci per field and classified as 0 (<0.5 foci/field), 1 (0.5–1.0 foci/field), 2 (1.0–2.0 foci/field), or 3 (>2.0 foci/field). The liver tumors were graded according to the Edmondson-Steiner grading system. Tumors were classified as grade I when well differentiated, grade II when moderately differentiated, grade III when poorly differentiated, and grade IV when undifferentiated or anaplastic. Nuclear atypia was scored separately on a 1–4 scale, corresponding to minimal, moderate, marked, or extreme atypia, respectively. Mitotic activity was evaluated in tumor hot spots using high-power fields (HPF) with a 40× objective. The mitotic index was categorized as low proliferative when 0–1 mitoses/HPF were observed, moderate when 2–5 mitoses/HPF were present, high when 6–10 mitoses/HPF were present, and very high when more than 10 mitoses/HPF were identified. The extent of necrosis and hemorrhage was scored independently using a 0–4 scale according to the percentage of affected tumor area: 0, absent; 1, 1–25%; 2, 26–50%; 3, 51–75%; and 4, greater than 75%.

### Human specimens

Frozen tumor biospecimens from 20 deidentified patients diagnosed with HCC were obtained from the Memorial Sloan Kettering Cancer Center Biobank and processed for targeted lipidomics analysis as described below. Formalin-fixed paraffin-embedded (FFPE) tissue microarrays (TMAs) containing HCC tumor samples (three cores per patient) were obtained from the Herbert Irving Comprehensive Cancer Center Molecular Pathology Shared Resource at Columbia University. Immunohistochemical staining was performed following standard protocols using anti-FADS2 (Invitrogen, PA5-48353) and anti-CD8 (CST, #70306T) antibodies. OCT-embedded tumor and adjacent liver tissue biospecimens from patients diagnosed with MASLD-associated HCC were obtained from the Herbert Irving Comprehensive Cancer Center Molecular Pathology Shared Resource. Histopathological evaluation and annotation of tumor and adjacent tissue regions were performed by liver pathologist X.W. Raman spectroscopy–based spectral lipidomics was subsequently performed as described below. Matched peripheral blood mononuclear cell (PBMC) and plasma samples were obtained from the Sylvester Comprehensive Cancer Center Biospecimen Shared Resource at the University of Miami from patients diagnosed with MASLD, MASLD-associated HCC, or non-MASLD-associated HCC. CD8⁺ T cell isolation and C11-Bodipy staining from PBMCs, and targeted lipidomics analyses of plasma samples were performed as described below. All human biospecimens were collected under protocols approved by the corresponding institutional review boards, and all samples were deidentified prior to analysis.

### Flow cytometry and immune profiling

Single-cell suspensions from liver tumors were generated following modified protocols^84^. Livers were harvested, washed in PBS, and tumor nodules were excised, finely minced, and digested in RPMI 1640 containing Dispase II (2.5 mg/mL), DNase I (1 mg/mL), Collagenase IA (0.365 mg/mL), Collagenase IV (0.75 mg/mL), and Hyaluronidase (0.125 mg/mL). Tissue fragments were incubated with agitation (250 rpm) for 30 min at 37 °C. Digestion was quenched with cold PBS, and cell suspensions were filtered through a 40 µm nylon strainer. The filtrate was centrifuged at 750 × g for 8 min at 4 °C, and the pellet was resuspended in RBC Lysis Buffer (BioLegend) for 3 min at 37 °C to lyse erythrocytes. The reaction was stopped with PBS, followed by centrifugation and resuspension in PBS supplemented with 3% FBS to obtain single-cell suspensions for antibody staining. For flow cytometry, 5 × 10^6 cells per sample were stained with LIVE/DEAD Fixable Viability Dye (Invitrogen, 1:1000) for 30 min on ice. Fc receptors were blocked with anti-CD16/32 TruStain FcX (BioLegend, 1:50). When indicated, Pentamer-SIINFEKL (ProImmune, 1:10) staining was performed at room temperature (RT) for 15 min. Surface antibody staining was carried out for 30 min on ice, protected from light. The following surface antibodies were used: CD3e (BioLegend, KT3.1.1, 1:25), CD4 (BD Biosciences, GK1.5, 1:25), CD8 (BioLegend, QA17A07, 1:50), CD11b (ThermoFisher, M1/70, 1:400), CD11c (BD Biosciences, N418, 1:50), CD19 (ThermoFisher, eB1o1D3, 1:50), CD25 (BD Biosciences, PC61, 1:25), CD44 (BD Bioscience, IM7, 1:50), CD45 (BD Biosciences, 30-F11, 1:400), CD62L (BioLegend, MEL-14, 1:50), CD69 (ThermoFisher, H1.2F3, 1:50), CD107a (BD Biosciences, 1D4B, 1:50), CD127 (BD Biosciences, SB/199, 1:50), KLGR1 (BD Biosciences, 2F1, 1:50), Ly6G (BD Biosciences, 1A8, 1:50), MHC-I-bound SIINFEKL (BioLegend, 25-D1.16, 1:25), MHC-II (BioLegend, M5/114.15.2, 1:200), NK1.1 (BioLegend, S17016D, 1:25), F4/80 (ThermoFisher, BM8, 1:50) and CD36 (BioLegend, HM36, 1:50). For intracellular staining, cells were fixed and permeabilized using the Foxp3 Transcription Factor Staining Buffer Set (Invitrogen) for 30 min, followed by incubation with intracellular antibodies for 30 min at RT, protected from light. The following intracellular antibodies were included: Granzyme B (BioLegend, QA16A02, 1:25), IFNγ (ThermoFisher, XMG1.2, 1:25), Perforin (BioLegend, S16009A, 1:25), Ki67 (ThermoFisher, SolA15, 1:100) and TNFα (BD Biosciences, MP6-XT22, 1:25). Samples were acquired on a Sony ID7000 spectral cytometer at the Columbia Stem Cell Initiative Flow Cytometry Core, and data were analyzed using FlowJo v10 software (Tree Star).

### Multiplexed immunofluorescence

Tissue slides were baked at 64°C for 30 min and washed three times with 1× PBS (pH 7.4) under gentle rocking conditions. Tissue permeabilization was performed using 0.3% Triton X-100 for 15 min at RT, followed by two PBS washes. Samples were then blocked for 1 h at RT using blocking buffer consisting of 1% BSA in PBS supplemented with mouse BD Fc Block. Nuclear staining was performed with DAPI diluted 1:1000 in PBS for 5 min at RT, followed by three PBS washes. Slides were mounted in 50% glycerol/PBS mounting medium and loaded into the Leica Cell DIVE platform for region-of-interest selection and imaging acquisition. Sequential multiplex immunofluorescence imaging was performed using DAPI, FITC, Cy3, Cy5, and Cy7 channels on the Leica Cell DIVE system. For primary antibody staining, antibodies were diluted in 3% BSA in PBS, and 300 µL of staining solution was added per slide, followed by overnight incubation at 4°C on a rotator. Slides were washed three times using PBS containing 0.01% Tween-20. For staining rounds requiring secondary antibodies, secondary antibodies were diluted in 3% BSA in PBS, incubated for 1 h at RT, and washed three times afterward. Following image acquisition, fluorophores were chemically inactivated using 0.1 M Na₂CO₃/3% H₂O₂ solution for 15 min at RT. Autofluorescence scans were reacquired between staining cycles for background subtraction. Image analysis and quantification were done using HALO v4.2 software (Indica Labs). Cell segmentation was performed using a trained segmentation algorithm, and positivity thresholds were defined individually for each marker. Tissue artifacts and staining artifacts were manually annotated and excluded from analysis. Quantification was performed within selected tissue regions of interest (ROIs).

### Cell culture

All cell lines used in this study were purchased from ATCC or similar repositories and collaborators. All cell lines were confirmed via STR profiling. Primary mouse HCC cell lines were cultured in plates pre-treated with collagen (50 μg/mL) in Dulbecco’s modified Eagle medium (DMEM) supplemented with 2 mM L-glutamine, 500 U/mL each of penicillin and streptomycin, 10% FBS, and plasmocin (5 μg/mL), and maintained in a humidified incubator at 37°C with 5% CO2. All reagents were obtained from Sigma-Aldrich or Fisher Scientific unless otherwise stated.

Lentiviral transductions were performed using virus produced in HEK293T cells transfected with psPAX2, pVSV.G, and the target construct, followed by transduction of HepG2 cells in the presence of polybrene (4 µg/mL). pKLV vector was used to generate shRNA cell lines targeting *DJ-1* (5’-GGGTAGCCGTGATGTGGTCAT-3’) or a non-targeting sequence (shCtrl) (5’-GGCCTAAGGTTAAGTCGCCCTCG-3’).

Fatty acid/BSA conjugates for *in vitro* assays were prepared based on established methods described previously^85^. Briefly, fatty acids were initially dissolved in 0.1 M NaOH to prepare 100 mM stock solutions. The mixture was warmed at 85 °C and vortexed at 2000 rpm until fully solubilized and clear. Stocks were then added dropwise to 5% fatty acid–free BSA to achieve a final concentration of 5 mM and incubated at 37 °C for 30 min to allow efficient conjugation of fatty acids to BSA. The resulting solutions were filtered through a 0.2 µm membrane and stored at – 20 °C until further use. For lipid treatments in HepG2 cells, an equimolar mixture of OL, PA, and LA (50 µM each) was applied for 24 h. When indicated, cells were co-treated with etomoxir (5 µM) or DJ-1 inhibitor (5 µM) for the same duration. For LA uptake assays, HepG2 cells were incubated with 2.5 µM Linoleate-Bodipy (Cayman Chemical) for 1 h at 37 °C, followed by quantification by flow cytometry as described above.

For the Click-based uptake assay, cells were incubated with alkyne-LA at 100 µM for 24 h. Click chemistry was performed using a modified protocol of the Click-iT™ HPG Alexa Fluor™ 594 Protein Synthesis Assay Kit (Thermo Fisher Scientific). Briefly, after alkyne incubation, cells were harvested, fixed in 3.7% paraformaldehyde for 15 min at RT (RT), and permeabilized with 0.5% Triton X-100 for 20 min at RT. The click reaction cocktail was freshly prepared by mixing 1X Click-iT HPG reaction buffer, copper sulfate, Alexa Fluor 488, and 10X Click-iT HPG buffer additive at a ratio of 2000:43:0.5:10, respectively. Cells were incubated with the cocktail for 30 min at RT in the dark and washed once with reaction rinse buffer and subsequently washed twice with PBS supplemented with 3% FBS. All washes were performed with PBS supplemented with 3% FBS. Samples were analyzed by flow cytometry.

Conditioned media (CM) from alkyne-LA-treated HepG2 cells was generated as follows. HepG2 cells were treated with 100 µM alkyne-LA for 24 h, then trypsinized and washed five times with PBS before replating in new culture plates. After 48 h of incubation to allow secretion of alkyne-LA–derived metabolites, the CM was collected, centrifuged to remove debris, and the supernatant was stored at –80°C. *In vitro*-stimulated CD8^+^ T cells were incubated with CM for 48 h, after which Click chemistry was performed as described above.

### CD8^+^ T cell isolation and *in vitro* stimulation

Naïve CD8^+^ T cells were isolated from the spleen and lymph nodes of 8-10-week-old female wild-type C57BL/6 mice by negative enrichment using the MagniSort™ Mouse CD8 T Cell Enrichment Kit (Thermo Fisher Scientific) according to the manufacturer’s instructions. A total of 1 × 10^5 CD8^+^ T cells per well were plated in 96-well plates in RPMI 1640 \emented with 10% charcoal-stripped FBS, 2 mM L-glutamine, 500 U/mL penicillin, 500 U/mL streptomycin, 1 mM sodium pyruvate, non-essential amino acids, 50 µM β-mercaptoethanol, and 50 ng/mL IL-2 (BioLegend). For T cell stimulation, cells were co-cultured with pre-washed anti-CD3/anti-CD28–coated magnetic beads (Dynabeads™ Mouse T-Activator, Thermo Fisher Scientific) at a bead-to-cell ratio of 1:1 and incubated at 37 °C for 72 h. Non-stimulated controls were maintained in the same culture medium without bead addition. Fatty acid treatments were performed at the indicated concentrations for 72 h. Where indicated, etomoxir (5 µM) or thioridazine (2 µM) was added during the last 48 h of activation. C11-Bodipy and JC-1 staining were performed at 2 µM for 30 min at 37 °C. For experiments with alkyne-LA-treated HepG2 CM, CD8^+^ T cells were stimulated for 24 h and then incubated with CM diluted 1:1 with culture medium for the following 48 h. Click chemistry was performed as described above. For PMA/ionomycin stimulation, CD8^+^ T cells were pre-treated for 48 h with the indicated fatty acids and subsequently stimulated with 50 ng/mL PMA and 500 ng/mL ionomycin for 5 h. Surface and intracellular markers, as well as dye stainings, were analyzed by flow cytometry as described previously. Total glutathione ratio was measured using the luminescent-based GSH/GSSG-Glo Assay (Promega, #V6611) following the manufacturer’s instructions.

### Human CD8⁺ T Cell Isolation and C11-Bodipy Staining

Cryopreserved patient-derived PBMCs were thawed, washed with 1× PBS, and resuspended in RPMI medium supplemented with glutamine, 10% FBS, and 10 mM HEPES for cell counting. CD8⁺ T cells were isolated by negative selection using the Human CD8⁺ T Cell Isolation Kit (Miltenyi Biotec, Cat. no. 130-096-495) according to the manufacturer’s instructions. The enriched CD8⁺ T-cell fraction was stained with 2 µM C11-Bodipy 581/591 C11 (Invitrogen, D3861) for 45 min at 37°C. Following staining, cells were washed once with 1× PBS and resuspended in cold FACS buffer (PBS supplemented with 1% FBS). Unstained control samples were prepared in parallel for each patient sample to establish autofluorescence baselines. Flow cytometric acquisition was performed using a CytoFLEX SRT flow cytometer (Beckman Coulter). Fluorescence was detected in the B525 (FITC) and Y585 (PE) channels, corresponding to the oxidized and reduced forms of C11-Bodipy 581/591, respectively. Samples were maintained on ice and protected from light prior to acquisition.

## Metabolomics analysis

### Untargeted metabolomics on murine MP53 and MPn HCCs

Untargeted metabolomic profiling was performed on mouse liver tumor homogenates using high-resolution accurate-mass (HRAM) spectrometry, as described previously^86^. Metabolites were extracted from tissue samples spiked with an internal standard cocktail by two-step cold homogenization and protein precipitation using methanol/water (1:3, v/v) and methanol/acetonitrile (1:1, v/v) at a final 1:10 (w/v) ratio. Ten microliters of extract were analyzed on a Vanquish™ Duo UHPLC system coupled to a Q Exactive HF-X Orbitrap mass spectrometer (Thermo Fisher Scientific). Chromatographic separation was conducted in both positive (HILIC) and negative (RP) ion modes using Waters BEH Amide and Higgins Targa C18 columns, respectively, under optimized gradient conditions at 60 °C. HRMS data were acquired in full-scan and data-dependent MS2 modes at resolutions of 120,000 and 7,500, respectively, with standard source parameters for both polarities. Raw data were processed in Compound Discoverer (v3.3.1) with <5 ppm mass error, S/N ≥ 3, and adaptive retention time alignment. Features were annotated using mzVault, mzCloud, and ChemSpider against HMDB, KEGG, and LipidMAPS databases.

### Untargeted lipidomics on murine MP53 HCCs

Lipid extraction was adapted from Matyash *et al*.^87^. All solvents were LC-MS grade. Fresh-frozen liver tumor tissue was homogenized in ice-cold methanol containing 0.1% (w/v) butylated hydroxytoluene using ceramic beads (Revvity Soft Tissue Homogenizing Mix, 1.4 mm) and a Revvity Omni Bead Ruptor Homogenizer at a 1:20 tissue-to-solvent ratio (mg/µL). Homogenates (300 µL) were transferred to glass tubes containing 1 µL SPLASH Lipidomics Internal Standard Mix (Avanti Polar Lipids). Lipids were extracted with 1 mL ice-cold methyl tert-butyl ether (MTBE), vortexed, and incubated overnight at −20°C. Phase separation was induced by addition of 250 µL ice-cold water followed by centrifugation. The upper organic phase was collected, dried under nitrogen, and stored at −80°C until analysis. Prior to LC-MS analysis, samples were reconstituted in isopropanol:acetonitrile:water (4:3:1, v/v/v) and centrifuged to remove particulates. Quality control (QC) samples were generated by pooling 10 µL from each sample.

LC-MS analysis was adapted from Tian *et al.*^88^. Samples were analyzed on a SYNAPT XS mass spectrometer coupled to an ACQUITY Premier UPLC system (Waters Corp). Lipids were separated on an ACQUITY Premier Peptide CSH C18 column (1.7 µm, 2.1 × 100 mm) using a 20-min gradient with solvent A (60:40 acetonitrile:water containing 10 mM ammonium acetate and 0.1% acetic acid) and solvent B (85:10:5 isopropanol:acetonitrile:water containing 10 mM ammonium acetate and 0.1% acetic acid). The gradient progressed from 40% to 99% solvent B, followed by column re-equilibration. Column temperature was maintained at 65°C and flow rate at 400 µL/min. Injection volumes were 3 µL (positive mode) and 6 µL (negative mode). Samples were analyzed in both positive and negative electrospray ionization modes. Source parameters included capillary voltage of 2.2 kV, sampling cone voltage of 30 V, source temperature of 120°C, and desolvation temperature of 500°C. Nitrogen was used as desolvation gas at 900 L/h. Leucine enkephalin was used as lock mass, and sodium formate was used for mass calibration. Data were acquired in data-independent acquisition (MSE) mode across m/z 50–1200 with a scan time of 0.1 s. Low collision energy was set to 4 eV and high collision energy was ramped from 25–60 eV. Fatty acid features were annotated based on accurate mass and retention time and validated against the LIPID MAPS database^89^ (using [M−H]⁻ adducts with a 5 ppm mass tolerance. Relative fatty acid abundance was quantified by peak area integration using QuanLynx software within MassLynx (Waters Corp).

Targeted ω-6 lipidomics on TIFs, TIF-treated CD8^+^ T cells, and isolated tumor-infiltrating CD8^+^ T cells; and untargeted lipidomics on TIF-treated CD8^+^ T cells were performed by Creative Proteomics following internal standard protocols.

### Targeted ω-6 PUFA lipidomics on frozen patient HCC tumors

Tissues were ground in a mortar and pestle pre-cooled with liquid nitrogen. Approximately 30 mg of powdered tissue was weighed into 2 mL bead tubes pre-filled with 1.4 mm ceramic beads (OMNI International). Metabolites were extracted using methanol:water (60:40, v/v) containing 0.5 µM palmitic acid-d31 (Cayman Chemical) as an internal standard, with extraction solvent added at a ratio of 30 µL per mg of tissue. Samples were further homogenized using a Biotage Bead Ruptor at 4°C. Following homogenization, 500 µL of the homogenate was transferred to a new Eppendorf tube, and 600 µL of chloroform was added. Samples were vortexed, incubated on ice, and centrifuged at 14,000 g for 5 min at 4°C. The lower, chloroform layer was collected, transferred to a clean glass vial, and dried under nitrogen. Dried extracts were reconstituted in 100 µL of acetonitrile:isopropanol (1:1, v/v) prior to LC–MS analysis. Fatty acid profiling was performed using an Agilent 6546 Q-TOF mass spectrometer operated in negative ion mode, coupled to an ACQUITY UPLC BEH C18 column (100 mm × 2.1 mm, 1.7 µm; Waters). Mobile phase A consisted of 10 mM ammonium acetate in 60:40 acetonitrile:water, and mobile phase B consisted of 10 mM ammonium acetate in 85:10:5 isopropanol:acetonitrile:water. The LC gradient was as follows: 0 min, 10% B; 2.0 min, 10% B; 16 min, 60% B; 21 min, 100% B; 24.1 min, 10% B; 27.6 min, 10% B. Other LC parameters included a flow rate of 0.3 mL/min, a column temperature of 45 °C, and an injection volume of 5 µL. MS source parameters included: gas temperature: 200°C; gas flow: 11 L/min; nebulizer pressure: 50 psig; sheath gas temperature: 300°C; sheath gas flow: 12 L/min; VCap: 3000 V; nozzle voltage: 0 V; fragmentor: 150V. Data were acquired from m/z 50–1700 with active reference mass correction (m/z 301.9981 and 980.0163). Data processing, including peak identification and integration, was carried out using Skyline.

### Targeted ω-6 PUFA lipidomics on human plasma biospecimens

Plasma samples (10 µL) were combined with 100 µL of butanol:methanol (1:1, v/v) containing 10 mM ammonium formate and a fatty acid internal standard (FA 18:1-d9; Avanti Polar Lipids). Samples were vortexed thoroughly and subjected to sonication in a water bath for 60 minutes at room temperature to facilitate lipid extraction. Following sonication, samples were centrifuged at 14,000 × g for 10 minutes at 4°C. The resulting supernatants were carefully transferred into glass insert vials for subsequent chromatographic analysis. Targeted lipidomics was performed using an Agilent 1290 ultra-high-performance liquid chromatography (UHPLC) system coupled to an Agilent 6495C triple quadrupole (QqQ) mass spectrometer. Chromatographic separation was achieved on a ZORBAX Eclipse Plus C18 column (2.1×100mm 1.8mm, Agilent) with the thermostat set at 45°C. Data were acquired using Agilent MassHunter Acquisition software (version 12.1). The mobile phase comprised (A) water/acetonitrile/isopropanol (50:30:20, v/v/v) and (B) water/acetonitrile/isopropanol (1:9:90, v/v/v), both containing 10 mM ammonium formate. Following a 1 uL sample injection, a stepped linear gradient was applied at a constant flow rate of 0.4 mL/min. The elution profile was programmed as follows: 15% B (0 min); 50% B (2.5 min); 57% B (2.6 min); 70% B (9 min); 93% B (9.1 min); 96% B (11 min); 100% B (11.1–12.0 min); and re-equilibration at 15% B from 12.2 to 16 min. To prevent carryover and maintain system integrity, a 1:1 isopropanol/water (v/v) seal wash and a 1:1 butanol/methanol (v/v) needle wash were employed.

Mass spectrometry was conducted in positive electrospray ionization (ESI+) mode using multiple reaction monitoring (MRM) mode. Source parameters were optimized as follows: gas temperature 150 °C, gas flow 17 L/min, nebulizer 20 psi, sheath gas temperature 200 °C, sheath gas flow 10 L/min, dwell time 100 ms, capillary voltage 3500 V, and Nozzle voltage 1000 V. The relative abundance in each sample was quantified by using Agilent Quantitative software 12.1.

## Spatial lipidomics by Raman spectroscopy

The lipid standards PA (C16:0) (CAS # 57-10-3), OA (C18:1) (CAS # 112-80-1), LA (C18:2) (CAS # 60-33-3), GLA (C18:3) (CAS # 506-26-3), DGLA (C20:3) (CAS # 1783-84-2), AA (C20:4) (CAS # 506-32-1), DTA (C22:4) (CAS # 122068-08-0), DPA (C22:5) (CAS # 24880-45-3), and DHA (C22:6) (CAS # 6217-54-5), were purchased from Cayman Chemical. All the Raman experiments were performed using Renishaw inVia Reflex spectrometer (785 nm laser, 50X long-working distance (LWD) air objective, 1200 grating). All samples were mounted on stainless-steel slides to minimize background Raman signals. Raman spectra were recorded from the lipid standards at 10% power (18 mW), 10 s exposure, and 100-3200 cm^-1^ range using extended mode. Raman spectra were preprocessed according to previously reported protocols^58^, including the removal of cosmic-ray artifacts and the subtraction of background signals. Principal component analysis (PCA) was then applied to the spectral datasets to enable multivariate analysis. This approach facilitated clustering of lipid standards based on spectral similarity, confirming that their Raman signatures were sufficiently distinct to support accurate mapping with minimal overlap. Flash-frozen samples (mice and patients) were cryosectioned to 20 μm and placed on stainless-steel slides without any additional sample preparation. To enable direct correspondence between histopathology/ immunophenotyping and Raman-derived biochemical maps, each Raman section was collected as an adjacent serial section to the corresponding 5 μm section used for H&E and IHC. Sections were stored at –80 °C prior to imaging to minimize lipid degradation. Raman imaging of tumor and adjacent normal tissues was performed using a raster-scanning approach (Stream^HR^ mode) at 100% laser power (180 mW) with a 7 s integration time and a spectral center at 1200 cm⁻¹ in static mode. Data were acquired across the entire sample area with a spatial step size of 100 μm (mouse tissues) and 200 μm (patient tissues). The total number of spectra collected per sample varied with sample dimensions, typically exceeding 6,000 spectra per specimen. To perform targeted analysis on specific regions of interest (e.g., only the tumor region or only the adjacent healthy region) within a sample, we used the “mask” feature of Renishaw WiRE 5.2. Lipid distributions within the mapped regions were quantified using Direct Classical Least Squares (DCLS) regression based on the curated fatty acid reference library described above. The resulting regression coefficients were mapped back onto the original spatial coordinates to reconstruct two-dimensional component images for each molecular species. These coefficients were treated as semi-quantitative indicators of the relative abundance of the lipid species. DCLS, a supervised multivariate method, uses Beer–Lambert Law and models the measured signal (D) as a linear combination of known reference spectra (S), their corresponding concentration-related coefficients (C), and residual error (E), expressed as D = C·Sᵀ + E.

## Mass spectrometry

### AlkMGO chemoproteomics

AlkMGO was synthesized and freshly deprotected as described previously^90^. HepG2 and MCF7 cells were treated with 3 mM of AlkMG for 5 h at 37°C, trypsinized, and washed twice with cold phosphate buffered saline (PBS). The cell pellets were resuspended in RIPA buffer (50 mM HEPES, pH7.4, 150 mM NaCl, 1% sodium deoxycholate, 1% NP-40, 0.5% SDS, 2.5 mM MgCl2, 10 mM sodium glycerophosphate, 10 mM sodium biphosphate) containing protease inhibitor cocktail and 25 unit/mL benzonase. Cell lysates were incubated on ice for 15 min, sonicated on ice twice, and centrifuged at 13000 x *g* for 10 min at 4°C. The concentration of the supernatant was measured by BCA assay, and equal amount of lysate was subjected to CHCl_3_/MeOH precipitation to remove any remaining AlkMG in the cell lysate. The white protein dish was then resuspended in 2% SDS (50 mM HEPES, pH 7.4, 150 mM NaCl, 2.5 mM TCEP) and sonicated once. The resuspended lysate was clicked with biotin alkyne (200 µM biotin-PEG3-azide, 500 µM CuSO_4_, 1 mM THPTA, 5 mM NaAsc, and 5% DMSO) for 3 h at RT with rocking. This was followed by CHCl_3_/MeOH precipitation again to remove excess biotin-PEG3-azide, and the lysate was resuspended in 200 µL 2% SDS (50 mM HEPES, pH 7.4, 150 mM NaCl, 2.5 mM TCEP), sonicated, and diluted with RIPA (final SDS < 0.5%). Meanwhile, 20 µL of high-capacity streptavidin agarose beads multiplied by the number of samples were washed with 1 mL RIPA and distributed to Eppendorf tubes. The lysates were added to the beads and incubated overnight at RT on a rotator. The beads were washed with RIPA ×2, 1 M KCl, 0.1 M Na _2_CO_3_, 2 M urea in HEPES buffer ×2, RIPA ×2, and water x3. After the final wash with water, the samples were quantified using a BCA assay. For this experiment, 200 µg of proteins were digested by trypsin. After protein digestion, approximately 100 µg of peptides were labeled using a TMT chemical labeling approach with a TMT 11 plex kit. The TMT incorporation efficiency was evaluated via ratio-check. Peptides were fractionated into 8 fractions using high-pH fractionation and combined into 4 fractions. The samples were run on an Eclipse mass spectrometer using a TMT pro SPS MS3 method, coupled to a Neo Vanquish liquid chromatography system, using a 4-hour gradient. For quality control, standards were run before and after the project to check instrument conditions. The mass spectrometry runs, along with the number of peptides and proteins identified and quantified, were consistent with expectations for these types of experiments. The data analyses were performed with Proteome Discoverer (version 2.4) and the data was searched against the SwissProt Human database (downloaded on 2/20/2024). A 1% FDR threshold was applied.

For AlkMGO immunoblot, cell lysis, protein precipitation, and click chemistry were performed as described above for the proteomics workflow. Proteins were subsequently resolved by SDS–PAGE and processed for immunoblotting as described below.

### FABP5-FLAG immunoprecipitation and mass spectroscopy for glycation site identification

10 cm plates of 70% confluent HEK293T cells were transfected with 8 ug of FABP5-MYC-FLAG plasmid and 20 uL of Lipofectamine 2000 for 16 hours. Cells were then treated with 0 mM or 3 mM MGO in DMEM for 4 hours. Cell pellets were lysed and sonicated (2 x 25A for 10 s) in lysis buffer (10 mM HEPES, 150 mM KCl, 1% Triton X-100, protease inhibitor). 50 uL of Anti-FLAG M2 Affinity Gel (Sigma, A2220) and 1500 ug of protein lysate were used to isolate FABP5-MYC-FLAG as described by the manufacturer. The beads were washed 3x with PBS and were run via western blot or digested and analyzed via mass spectrometry.

For mass spectrometry analysis of FABP5 glycated residues, on-bead tryptic digestion was performed on the bead-bound sample using a 3-stage trypsin strategy. Proteins were reduced with DTT (30 mins, 37°C) and alkylated with IAA (45 min, 25°C) in a 2 M urea / 50 mM ammonium bicarbonate digestion buffer. An initial trypsin digestion was performed (1 hr, 37°C), followed by a second overnight trypsin incubation (37°C) to maximize sequence coverage. A third addition of trypsin the following day was included as a standard step to ensure complete digestion. The resulting peptide mixture was acidified, quantified by NanoDrop, and desalted via C18 StageTip solid-phase extraction using LC/MS-grade solvents throughout. Purified peptides are dried by SpeedVac, reconstituted in 0.1% formic acid, and transferred to LC vials for MS analysis.

Raw data files were processed using Spectronaut version 20.5 (Biognosys) and searched with the PULSAR search engine with Uniprot *Homo sapien* protein database downloaded on 2025/06/18 (42,533 entries). cysteine carbamidomethylation was specified as fixed modifications, while Methionine oxidation, acetylation of the protein N-terminus and Deamidation (NQ), CEL/CEA (KR) and MG-H1(R) were set as variable modification. A maximum of three trypsin missed cleavages were permitted. Searches used a reversed sequence decoy strategy to control peptide false discovery rate (FDR) and 1% FDR was set as threshold for identification. PTM localization filter was set as 0.75.

### Structural Analysis and Solvent Accessibility

Visualization and solvent accessible surface area (SASA) calculations were performed with the UCSF ChimeraX (v1.11.1) program using the crystal structure from PDB: 4LKP for the human FABP5 protein^78^. The relative solvent accessibilities (RSA) of the amino acid residues were calculated by normalizing SASA with the maximum allowed solvent accessibilities described by Tien *et. al*.^57^ and plotted against residue number in R.

### Quantification of MGO and AGEs from tumor tissue

For MGO quantification, tumor tissue was homogenized in 80:20 MeOH:H2O (–80 °C) containing 50 pmol ^13^C_3_-MGO. Samples were then spun down at 14,000 x *g* for 10 min at 4 °C. Half the supernatant was transferred to a new 1.7 mL tube for MGO quantification. 10 µL of 5 µM ^13^C-MGO and 10 µL of 1 mM *o*-phenylenediamine (*o*-PD) were added. Samples were vortexed and put in a long-axis rotator to derivatize for 2 h in the dark at room temperature. After derivatization, samples were spun at 14,000 x *g* for 10 min and plated for LC-MS/MS analysis. Samples were separated with a Shimadzu LC system with a 2.1 x 50 mm, 3 µm diameter Atlantis dC18 column (Waters) with a flow rate of 0.5 mL/min, using A (H_2_O with 0.1% formic acid) and B (ACN with 0.1% formic acid) buffers. MRM was performed using a Sciex 6500 QTRAP with the following transitions: m/z 145 –> 77 for MGO, 148 –> 77 for MGO-IS. Area ratios were determined using the IS and samples were normalized to mg of tissue.

For tumor AGE quantification, the tissues were cut and weighed (∼10-30 mg). Protein was homogenized in a lysis buffer containing 100 mM Tris-HCl, 150 mM NaCl, 10 mM NaF, 1% Triton X-100, pH = 7.4 with 1:500 protease and phosphatase inhibitor cocktails (Sigma Aldrich). Samples were then sonicated using a probe sonicator (10x 10 s pulses at 20% power). Samples were spun down at 14,000 x *g* for 10 min and clarified supernatant was used to conduct a BCA. 250 µg of soluble protein was then precipitated in acetone (–80 °C) for 1 h. Samples were again spun down at 14,000 x *g* for 10 min at 4 °C. Acetone was removed and the pellet was allowed to air dry. 65 µL of 50 mM freshly prepared ammonium bicarbonate was added. 10 µL of an internal standard mix (see above) and 5 µL of 0.1 mg/mL sequencing-grade trypsin was added to each sample. Digests were performed at 37 °C for 3 h. Samples were then boiled at 95 °C for 10 min, left at room temperature to cool, and briefly spun down via benchtop centrifuge. Then, 10 µL of 0.15 mg/mL aminopeptidase was added, and samples were left at 37 °C overnight. The next day, samples were again boiled at 95 °C for for 10 min, left at room temperature to cool for 5 min, and briefly spun down via benchtop centrifuge. 15 µL of a 1:1 HFBA:H_2_O solution was added to each sample. Samples were centrifuged for 14,000 x *g* for 10 min and plated. The same chromatographic conditions were used as above. MRM was performed with the following transitions: m/z 175 –> 70 for Arg, 185 –> 75 for Arg-IS, 247 –> 70 for CEA, 248 –> 70 for CEA-IS, 219 –> 84 for CEL, 223 –> 88 for CEL-IS, 132 –> 86 for Leu, 138 –> 91 for Leu-IS, 147 –> 84 for Lys, 155 –> 90 for Lys-IS, 229 –> 70 for MG-H1, 230 –> 70 for MG-H1-IS, 219 –> 84 for LactoylLys. Samples were quantified using the respective ^13^C-IS to analyte ratio and normalized to Leu concentration and µL of tissue.

### Immunoblot and qPCR

Proteins were extracted in ice-cold RIPA lysis buffer (25 mM Tris-HCl, pH 7.6; 150 mM NaCl; 1% NP-40; 1% sodium deoxycholate; 0.1% SDS) supplemented with protease and phosphatase inhibitor cocktails (Thermo Fisher Scientific). Samples were washed twice with ice-cold PBS and lysed directly in the same ice-cold RIPA buffer containing the inhibitors. All lysates were incubated on ice for 30 min with intermittent vortexing, followed by centrifugation at 14,000 × g for 15 min at 4 °C. The supernatants were collected, and protein concentrations were determined using Bradford protein assay (Bio-Rad) according to the manufacturer’s instructions. Equal amounts of protein (30 µg) were separated by SDS-polyacrylamide gel electrophoresis using 10-20% polyacrylamide gels (or 4-20% precast Mini-PROTEAN TGX gels, Bio-Rad; cat. no. 456-1094) and transferred to polyvinylidene difluoride (PVDF) membranes (Millipore; 0.2 µm pore size). Membranes were blocked either in LI-COR Odyssey blocking buffer for 1 h at RT, followed by overnight incubation at 4 °C with primary antibodies against DJ-1 (Cell Signaling Technology, #2134), FABP5 (Cell Signaling Technology, #39926), FABP4 (Cell Signaling Technology, #50699), FABP1 (R&D Systems, MAB2964), CD36 (Cell Signaling Technology, #28109), β-actin (Cell Signaling Technology, #3700), or GAPDH (Cell Signaling Technology, #5174). After three washes with TBST, membranes were incubated with IRDye-conjugated secondary antibodies (LI-COR Biosciences) for fluorescent detection using the Odyssey CLx Imaging System (LI-COR Biosciences).

For total RNA isolation RNeasy Mini Kit (Qiagen, Cat. No. 74106) was used and reverse transcribed using the LunaScript RT SuperMix Kit (NEB, Cat. No. E3010L) following the manufacturer’s protocol. Quantitative PCR was performed using TaqMan Gene Expression Master Mix (Thermo Fisher Scientific) for *Fads2* (Mm00517221_m1) and *Park7* (Mm00498538_m1) probes labeled with VIC or FAM fluorophores (Thermo Fisher Scientific). qPCR reaction was performed employing an Applied Biosystems QuantStudio 5 Real-Time PCR system (Applied Biosystems). For PCR analysis, the CT values were normalized to all housekeeping genes and analyzed using 2−ΔΔCT method.

## scRNAseq analysis

### Sample preparation

Tumors were harvested and processed as described for flow cytometry. Cells were stained with LIVE/DEAD Fixable Dye (Invitrogen, 1:1000) and anti-CD45 (BioLegend, clone 104, 1:1200). Viable CD45⁺ cells were FACS-sorted using a Sony MA900 for the MP53 experiment. For the temporal MP53-lucOS experiment, both viable CD45⁺ and CD45⁻ cells were collected. Fc receptors were blocked with anti-CD16/32 TruStain PLUS FcX (BioLegend, 1:50) for 30 min on ice, and cells from each mouse were barcoded using TotalSeq B anti-mouse Hashtag Antibodies (BioLegend, clones M1/42 and 30-F11; 1:250) following the cell hashing protocol^91^. All washes and incubation were done with PBS supplemented with 10% FBS. For the MP53 –lucOS experiment, CD45⁺ cells were spiked with 20% of CD45⁻ cells. Cell suspensions were adjusted to 1000 cells/µl after confirming >80% viability by Trypan Blue staining. For the MP53 and MPn experiment, four mice were multiplexed per lane, while for the MP53-lucOS experiment, two mice were multiplexed per lane. Approximately 20,000 cells were sequenced per lane. Droplet capture was performed using the Chromium Controller (10x Genomics) according to the manufacturer’s instructions, and libraries were sequenced on the Illumina NovaSeq 6000 platform at the Columbia University Single Cell Core.

### Data cleanup and quality control

Sequence reads from each sample were aligned to the *mm10* mouse reference genome and processed with CellRanger using default parameters. Count matrices were imported into Scanpy^92^ and merged into a single object. Cell hashtag demultiplexing was performed with HashSolo Demux (integrated in Scanpy), and barcodes classified as negatives or doublets were removed. Additional doublet detection and removal were carried out with Scrublet^93^. For the MP53 scRNA-seq experiment, genes detected in fewer than four cells were excluded. Cells with fewer than 500 reads or with mitochondrial content >40% were also filtered out, yielding a final dataset of 20,867 cells and 18,765 genes. For the temporal MP53-lucOS experiment, cells with fewer than 200 detected genes or with mitochondrial content >35% were removed, and genes expressed in fewer than three cells were excluded, resulting in a final dataset of 128,525 cells and 23,866 genes.

Following quality control, count matrices were normalized by total read count and log-transformed with a pseudocount of 0.1. The top 4,000 highly variable genes (HVG) were selected for principal component analysis (PCA, 50 components). A k-nearest neighbor graph was computed on the PCA space (k=30). For the MP53-lucOS dataset, batch correction was applied using Harmony^94^ with default settings. Data visualization was performed using UMAP with default parameters. Clustering was conducted using Leiden algorithm at resolution of 0.6. Differential gene expression analysis was carried out on log-normalized data using the Wilcoxon rank-sum test. Major immune cell populations were annotated based on the average expression of established marker genes.

### T cell analysis

T cell clusters from the MP53 and MPn experiment; and CD8⁺ T cell clusters from the MP53-lucOS experiment, were independently isolated and reclustered using the same parameters described above (HVG, PCA, UMAP, Leiden clustering, and differential gene expression analysis). CD8⁺ T cell subtypes were annotated based on the average expression of established markers for each subcluster, according to previous studies^95^. For the MP53 and MPn dataset, differential abundance of T cells between diets was assessed using MiloR^96^. In this approach, cells are grouped into “neighborhoods” based on a k-nearest neighbor graph, and counts are modeled with a negative binomial generalized linear model to test for differences in cell proportions. In the resulting plot, each dot represents a neighborhood, with color indicating enrichment in SD (red) or WD (blue) at FDR ≤ 0.05. To mitigate the effects of sparse gene expression, MAGIC^97^ was applied to denoise and impute the expression data with parameters k = 30, ka = 10, and t = 3.

## Differential expression and GSEA on immunotherapy-treated HCC patients

Differential gene expression (DGE) analysis was performed on transcriptomic data (n=247) from the GO30140 and IMbrave150 trials^61–63^, comparing responders (CR/PR) to atezolizumab + bevacizumab immunotherapy against non-responders (SD/PD). The analysis used the limma-voom pipeline implemented in the edgeR and limma R packages (versions 4.4.2). Lowly expressed genes were removed using the filterByExpr function and library sizes were normalized using the trimmed mean of M-values (TMM) method. DEGs were considered significant at an adjusted p-valued (FDR) < 0.2 and absolute log2 fold-change > log2(1.5).

Gene Set Enrichment Analysis (GSEA) was performed using the fgsea R package (version 4.4.3) on the pre-ranked gene list. Gene sets from the Molecular Signatures Database (MSigDB) were retrieved using the msigdbr package (version 4.4.3), including Hallmark (H) and Canonical Pathways (C2) collections. Enrichment scores were calculated using 10,000 permutations, considering gene sets with a minimum size of 15 and a maximum size of 500 genes. Pathways were considered significantly enriched at FDR < 0.05. Negative normalized enrichment scores (NES) indicate enrichment in non-responders. In addition, over-representation analysis (ORA) was performed on significantly upregulated genes in non-responders using the clusterProfiler package (version 4.4.2)^98^. Gene Ontology (GO) Biological Process enrichment analysis was conducted after converting gene symbols to Entrez IDs using the org.Hs.eg.db annotation database. Enriched GO terms were identified using a Benjamini–Hochberg adjusted p-value cutoff of 0.2 and a q-value cutoff of 0.3.

### Analysis of human transcriptomic data using publicly available signatures

Hallmarks, GO and C2 pathways derived from the GSEA analyses were used as input for the single-sample Gene Set Enrichment Analysis (ssGSEA) module of GenePattern to calculate enrichment scores per sample for each gene signature in the atezo+bev treated cohorts (n=247) [IMbrave150 phase III and GO30140 phase I trials], in addition to 53 MASH-HCC tumors, and compared them to those from other non-MASH etiologies (n=184)^34,35^. Furthermore, molecular and immune HCC subclasses were evaluated using the Nearest Template Prediction (NTP) module from GenePattern, categorizing each sample into presence or absence of each single – derived HCC gene signature^99^. Significant prediction was defined using Benjamini-Hochberg-adjusted FDR <0.2.

## CIBERSORT analysis

Immune cell composition in human HCC tumors was inferred using CIBERSORT^100^ on TIMER2 platform. Gene expression data from TCGA HCC samples were obtained, and patients were stratified into high or low expression groups based on either *PARK7* or *FADS2* gene expression. CIBERSORT was applied to each group to estimate the relative abundance of immune cell types, allowing comparison of immune infiltration patterns between high and low gene expression cohorts.

No unique code was developed for this study.

---

**Figure S1:**
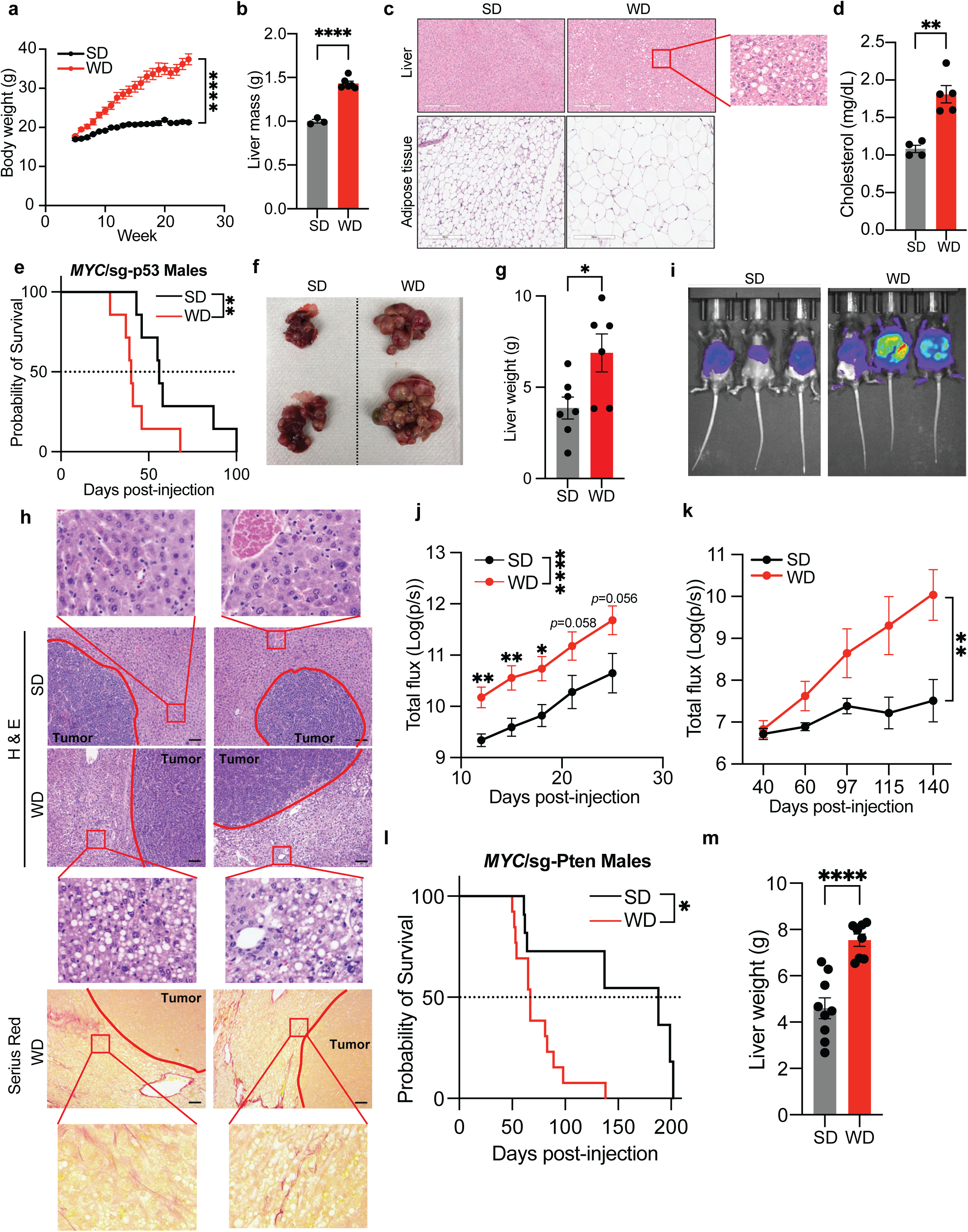
Overnutrition induces MASLD-associated liver dysfunction and hepatocarcinogenesis. **(a)** Body weight of female C57BL/6 mice fed SD (n=7) or WD (n=9) *ad libitum*; **(b)** Liver weight of mice from (a) after 24 weeks on SD (n=3) or WD (n=6) *ad libitum*; **(c)** Representative H&E-stained sections of liver (top) and adipose tissue (bottom) from 24-week SD– or WD-fed mice; **(d)** Serum cholesterol levels in SD– (n=4) and WD-fed (n=5) mice; **(e)** Kaplan–Meier survival curves of male mice bearing MP53 HCCs fed SD or WD (n=7); **(f)** Representative images of time-matched livers from MP53 HCC-bearing mice under SD or WD; **(g)** Liver weights of SD-(n=7) or WD-fed (n=6) MP HCCs; **(h)** Representative H&E (top) and Sirius Red (bottom) staining of tumour-bearing livers highlighting steatosis and incipient fibrosis in tumour-adjacent tissue; scale bar = 200 µm; **(i)** Representative IVIS bioluminescence images from mice bearing luciferase-expressing MP53 HCCs and fed indicated diet; **(j)** IVIS quantification of luciferase-expressing multifocal MP53 HCCs generated by HDTV injection (n≥5); **(k)** IVIS quantification of luciferase-expressing unifocal MP53 tumours generated by electroporation (n≥5); **(l)** Kaplan–Meier survival curves of male mice bearing MPn HCCs (SD: n=11; WD: n=13); **(m)** Liver weights of time-matched MPn HCCs (SD: n=9; WD: n=8). Results are shown as mean ± SEM. ns: not significant, *p < 0.05, **p < 0.01, ****p < 0.0001.

**Figure S2:**
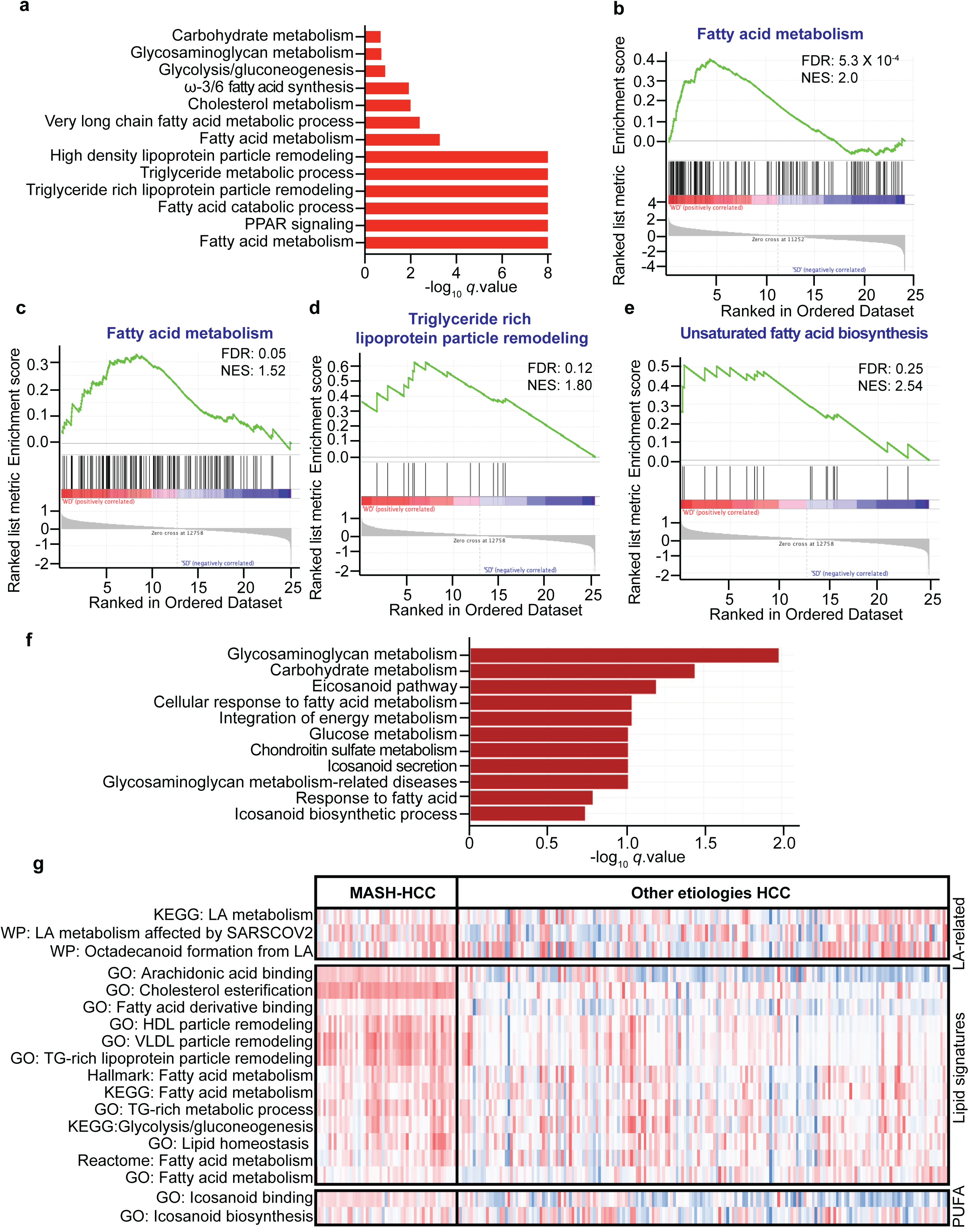
WD-driven HCCs exhibit enrichment of fatty acid– and PUFA-related metabolic pathways. **(a)** Pathway enrichment analysis of bulk transcriptomic data from WD-fed MP53 HCCs; **(b)** GSEA of KEGG fatty acid metabolism pathway in WD-fed MP53 tumours; **(c – e)** GSEA enrichment plots for fatty acid metabolism (c), triglyceride-lipoprotein metabolism (d), and unsaturated fatty acid biosynthesis (e) pathways in WD-fed MPn HCCs; **(f)** Enriched metabolic pathways in a WD-influenced orthotopic HCC model (Hep53.4 cells); **(g)** Heatmap of linoleic acid, lipid metabolism, and PUFA-related gene signatures in bulk transcriptomic data from HCC patient samples stratified by MASH (n=53) versus other etiologies (n=184). All p-values <0.01.

**Figure S3:**
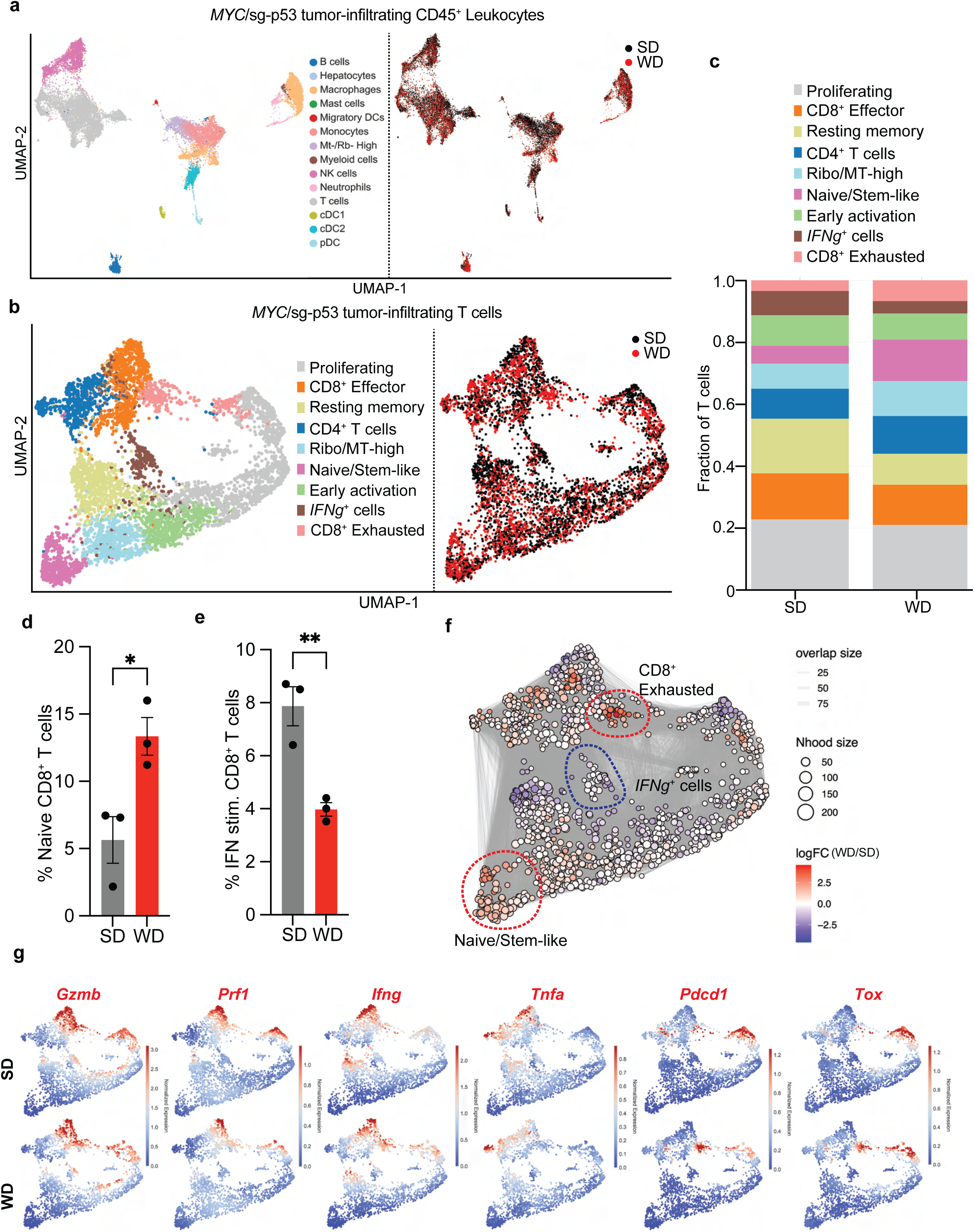
scRNA-seq analysis of tumour-infiltrating T cells from SD– and WD-fed MP53 HCCs. **(a)** UMAP projection of tumour-infiltrating CD45⁺ cells from MP53 HCCs colored by cell type (left) and dietary regimen (right) (SD: 12,979 cells; WD: 12,037 cells; n=3 mice). DCs: Dendritic cells; Mt: mitochondrial genes; Rb: ribosomal genes; NK: Natural killer; cDC: Conventional dendritic cells; pDC: plasmacytoid dendritic cells; **(b)** UMAP projection of tumour-infiltrating T cells colored by cell type (left) and dietary regimen (right) (SD: 4,933 cells; WD: 4,084 cells; n=3 mice); **(c)** Relative abundance of intratumoral T cell subsets in MP53 HCCs by dietary condition (n=3); **(d & e)** Percentage of naïve (d) and IFNγ⁺ (e) CD8⁺ T cells in MP53 HCCs under SD or WD (n=3); **(f)** K-nearest neighbor (Milo) analysis of T cell subsets enriched under SD (blue) or WD (red); **(g)** UMAPs showing expression of representative marker genes across T cell subsets stratified by diet. Results are shown as mean ± SEM. ns: not significant, *p < 0.05, **p < 0.01.

**Figure S4:**
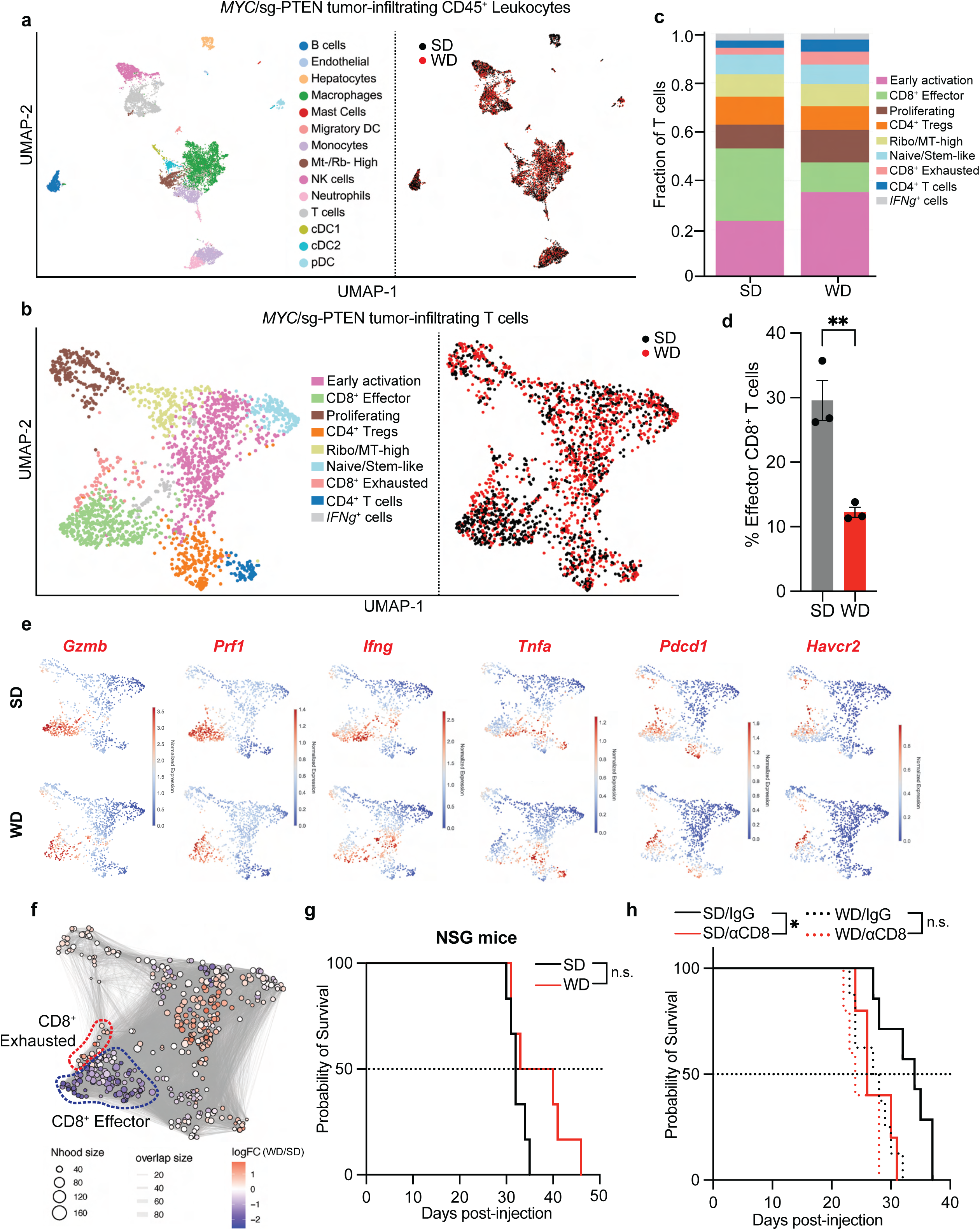
scRNA-seq analysis of tumour-infiltrating T cells from SD– and WD-fed MPn HCCs. **(a)** UMAP projection of MPn tumour-infiltrating CD45^+^ cells colored by cell type (left) and dietary regimen (right) (SD: 7,717 cells; WD: 5,965 cells; n=3 mice). DCs: Dendritic cells; Mt: mitochondrial genes; Rb: ribosomal genes; NK: Natural killer; cDC: Conventional dendritic cells; pDC: plasmacytoid dendritic cells; **(b)** UMAP projection of MPn tumour-infiltrating T cells colored by cell type (left) and dietary regimen (right) (SD: 1,128 cells; WD: 1,214; n=3 mice); **(c)** Relative abundance of intratumoral T-cell subsets in MPn HCCs across dietary conditions; **(d)** Percentage of effector CD8^+^ T cells in MPn HCCs under different diets (SD: n=3; WD: n=3); **(e)** UMAPs showing expression of representative marker genes in all T cells stratified by diet; **(f)** K-nearest neighbor (Milo) analysis of T cell types enriched in SD (blue) or WD (red); **(g)** Kaplan–Meier survival of immunodeficient NSG female mice bearing MP53 HCCs under SD or WD feeding (n=6); **(h)** Kaplan–Meier survival of female mice bearing MP53 HCCs under SD or WD feeding and treated with the indicated antibodies (SD/IgG: n=7, SD/αCD8a: n=5, WD/IgG: n=8, WD/αCD8a: n=5). Results are shown as mean ± SEM. ns: not significant, *p < 0.05.

**Figure S5:**
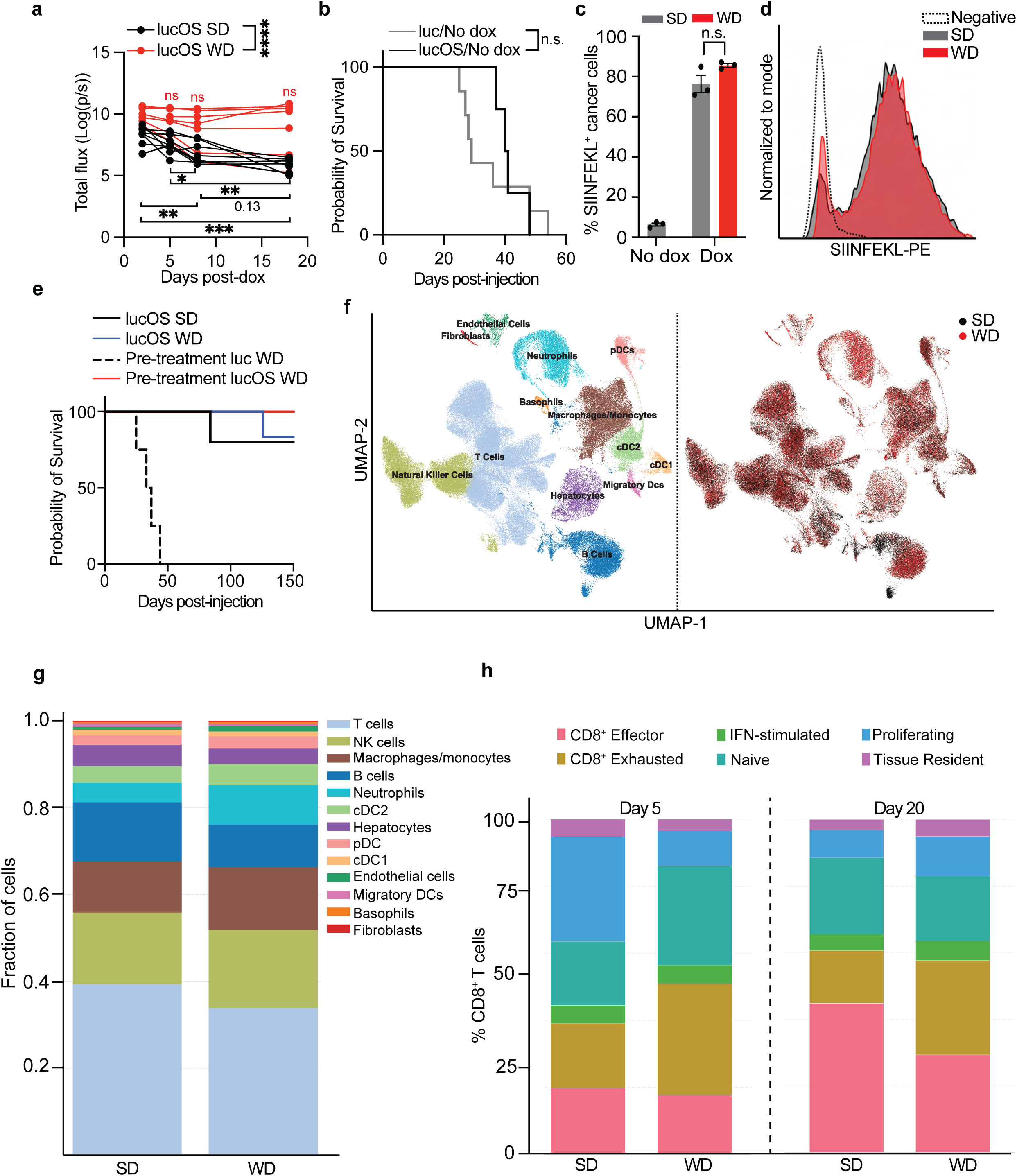
Characterization and scRNA-seq analysis of the dox-inducible immunogenic model MP53-lucOS. **(a)** IVIS quantification of lucOS-expressing MP53 HCCs under SD and WD regimens (n=7); **(b)** Kaplan–Meier survival of female animals bearing MP53-luc or MP53-lucOS HCCs under SD without dox supplementation (luc/No dox: n=7; lucOS/No dox: n=4); **(c & d)** Percentage (c) and representative histograms (d) of MHC-I-bound SIINFEKL^+^ cancer cells in SD-and WD-fed MP53-lucOS HCCs ± dox (n=3); **(e)** Survival of female mice bearing constitutive MP53-luc/lucOS HCCs with SD or WD feeding initiated pre– or post-tumour onset (Pre-treatment luc/WD: n=4; Pre-treatment lucOS/WD: n=10; lucOS/SD: n=5; lucOS/WD: n=6); **(f)** UMAP of MP53-LucOS tumour-infiltrating CD45⁺ cells and CD45⁻ cells colored by cell type (left) and diet (right) (SD: 66,517; WD: 72,008); **(g)** Relative abundance of tumoral cell types in MP53-LucOS HCCs across dietary conditions; **(h)** Relative abundance of intratumoral CD8^+^ T-cell subsets in MP53-LucOS HCCs across dietary conditions. Results are shown as mean ± SEM. ns: not significant, ****p < 0.0001.

**Figure S6:**
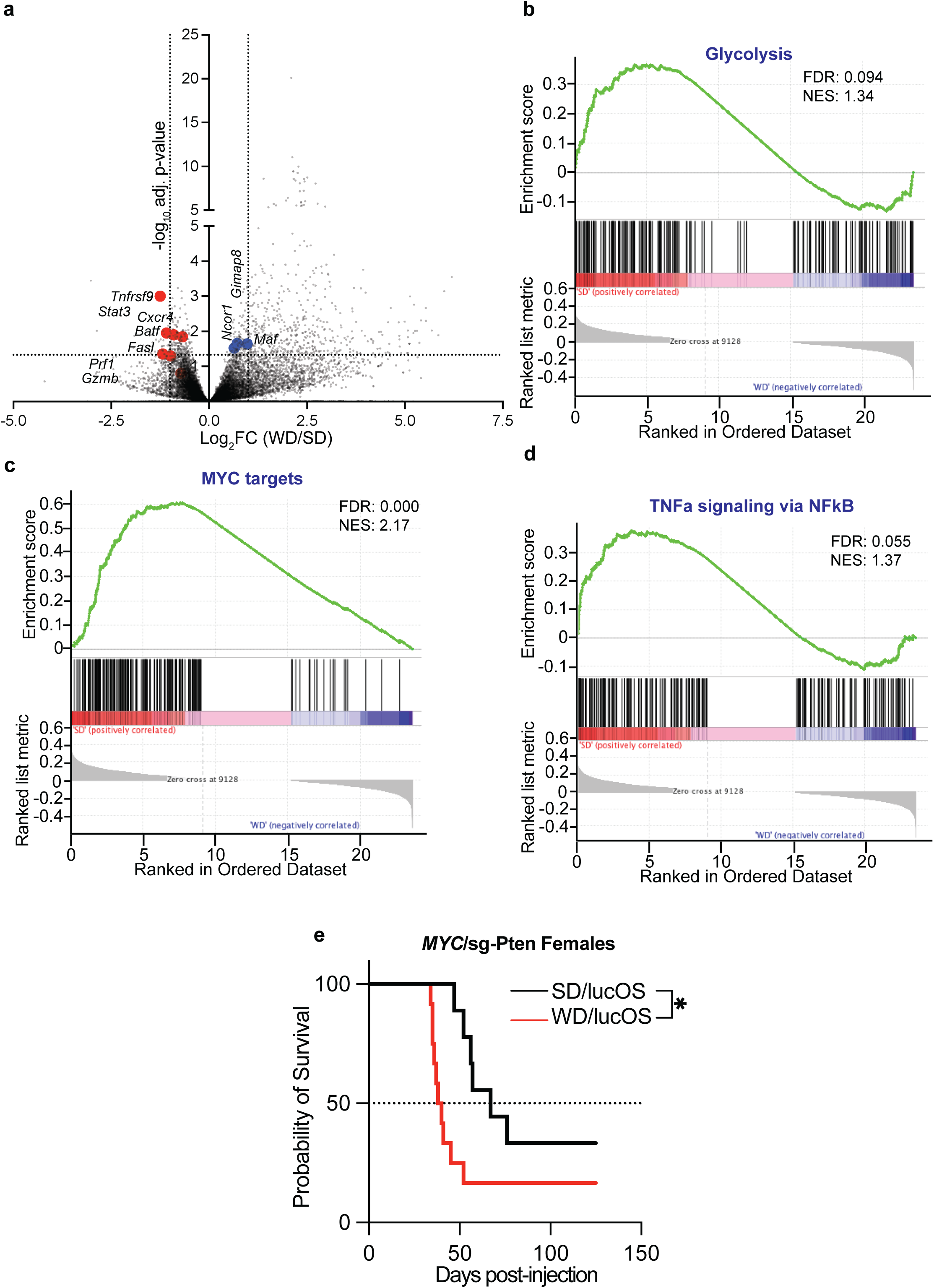
WD impairs antitumor response in tumour-infiltrating CD8^+^ T cells. (**a**) Differentially abundant genes in MP53-LucOS-infiltrating CD8^+^ T cells under SD (n=4) and WD (n=5); effector marker genes are highlighted in red and dysfunction/exhaustion-associated genes in blue; **(b-d)** GSEA of the CD8⁺ T cell transcriptome showing enrichment of metabolism (b), proliferation (c), and activation (d) gene programs in tumour-infiltrating CD8⁺ T cells from Mp53-lucOS HCCs (SD: n=4; WD: n=5); **(e)** Kaplan-Meier survival curves of mice bearing MPn-lucOS HCCs under SD or WD *ad libitum* (SD/lucOS: n=9; WD/lucOS: n=10). *p < 0.05.

**Figure S7:**
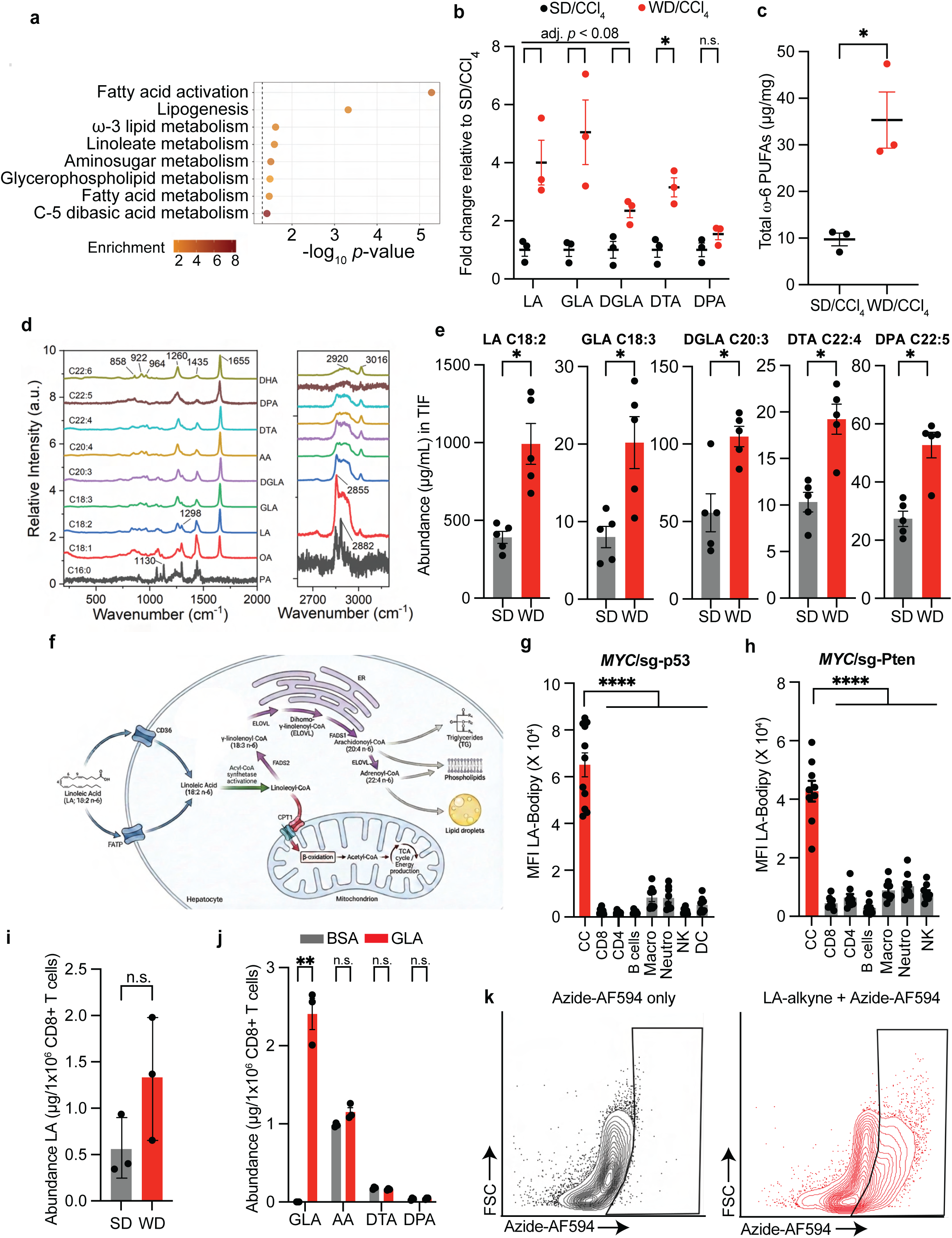
Dietary LA fuels tumour-derived ω-6 PUFA production and CD8⁺ T cell suppression. (**a**) Untargeted metabolomics-based enrichment in WD-fed MPn HCCs (n=3); **(b & c)** Relative abundance of LA pathway fatty acids (b) and total ω-6 PUFA content (c) in the carcinomas from WD/CCl₄ treated mice relative to adenomas from SD/CCl₄ group (n=3); **(d)** Raman vibrational spectra signatures used for the quantification of indicated fatty acids; **(e)** Abundance of indicated ω-6 PUFAs in SD– or WD-derived TIFs (n=5); **(f)** LA metabolic pathway in cancer cells; FADS1: Fatty acid desaturase 1, ELOVL: Very long chain fatty acid elongase; FATP: Fatty Acid Transport Protein; CPT1: Carnitine palmitoyltransferase I; **(g & h)** LA-Bodipy uptake by the TME cellular subtypes of WD-fed MP53 (n=11 tumours from 4 mice) (g) and MPn (n=9 tumours from 3 mice) (h) HCCs. CC: Cancer cells, Macro: Macrophages, Neutro: Neutrophils, NK: Natural killer cells, DC: Dendritic cells; **(i)** LA abundance in tumour-infiltrating CD8^+^ T cells isolated from MP53 HCCs under SD or WD regimen (n=3), determined by targeted lipidomics; **(i)** Abundance of indicated ω-6 fatty acids in CD8⁺ T cells stimulated in the presence of BSA or GLA (100 µM, 24 h) (n=3); **(k)** Representative dot plots showing LA-alkyne-derived metabolite uptake by CD8⁺ T to conditioned media from LA-alkyne–treated HepG2 cells (100 µM, 24 h). Results are shown as mean ± SEM. ns: not significant, *p < 0.05, **p < 0.01, ****p < 0.0001.

**Figure S8:**
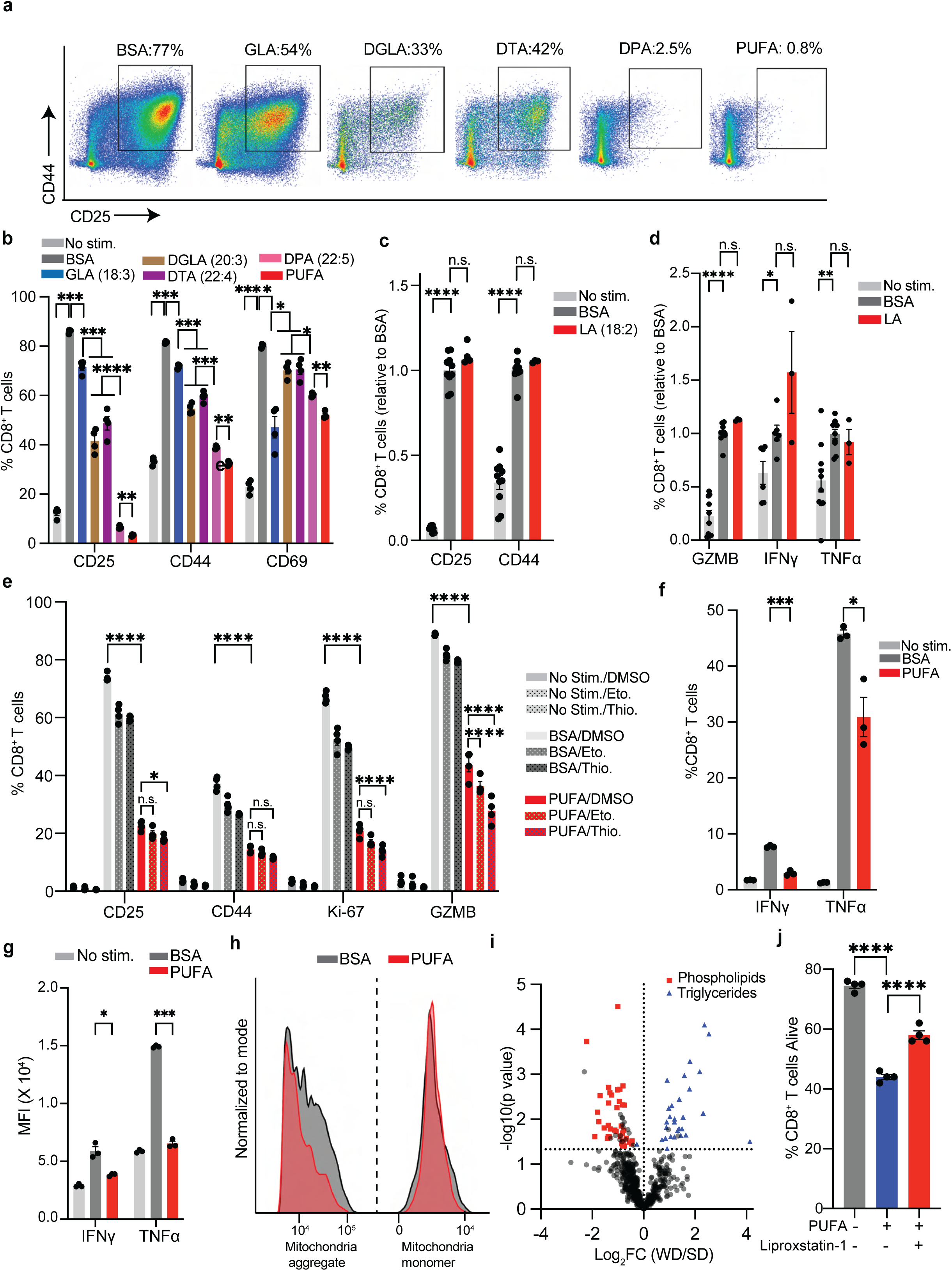
Effect of LA-derived ω-6 lipid intermediates on CD8^+^ T cell metabolic fitness and function. (**a**) Representative dot plots of CD44⁺/CD25⁺ frequency in CD8⁺ T cells stimulated in the presence of BSA or indicated ω-6 lipids (200 µM, 72 h). PUFA: equimolar mix of ω-6 lipids (50 µM); **(b)** CD8⁺ T cell activation markers after stimulation in the presence of indicated ω-6 lipids (200 µM, 72 h, n=4); **(c & d)** CD8⁺ T cell activation marker expression after stimulation with indicated BSA or LA (100 µM, 72 h, n≥3); **(e)** Frequency of marker-positive CD8⁺ T cells stimulated in the presence or absence of combined ω-6 lipids (PUFA) (50 µM each, 72 h) ± etomoxir (5 µM, 48 h), or ± thioridazine (2 µM, 48 h) (n=4); **(f & g)** Positive percentage (d) and levels (e) of indicated cytokines in CD8⁺ T cells stimulated with PMA/I ± combined ω-6 lipids (All) (50 µM each, 72 h) (n=3); PMA: Phorbol 12-myristate 13-acetate; I: Ionomycin; **(h)** Representative histograms of JC-1 staining showing aggregated (left) and monomeric (right) mitochondrial states in stimulated CD8⁺ T cells ± combined ω-6 lipids (PUFA) (50 µM each, 72 h); **(i)** Differentially abundant lipids in MP53 HCCs under SD or WD, determined by untargeted lipidomics (n=4); **(j)** %live CD8⁺ T cells stimulated in the presence or absence of ω-6 PUFA (50 µM each, 72 h) ± liproxstatin-1 (500 nM, 72 h) (n=4). Results are shown as mean ± SEM. ns: not significant, *p < 0.05, **p < 0.01, ***p < 0.001, ****p < 0.0001.

**Figure S9:**
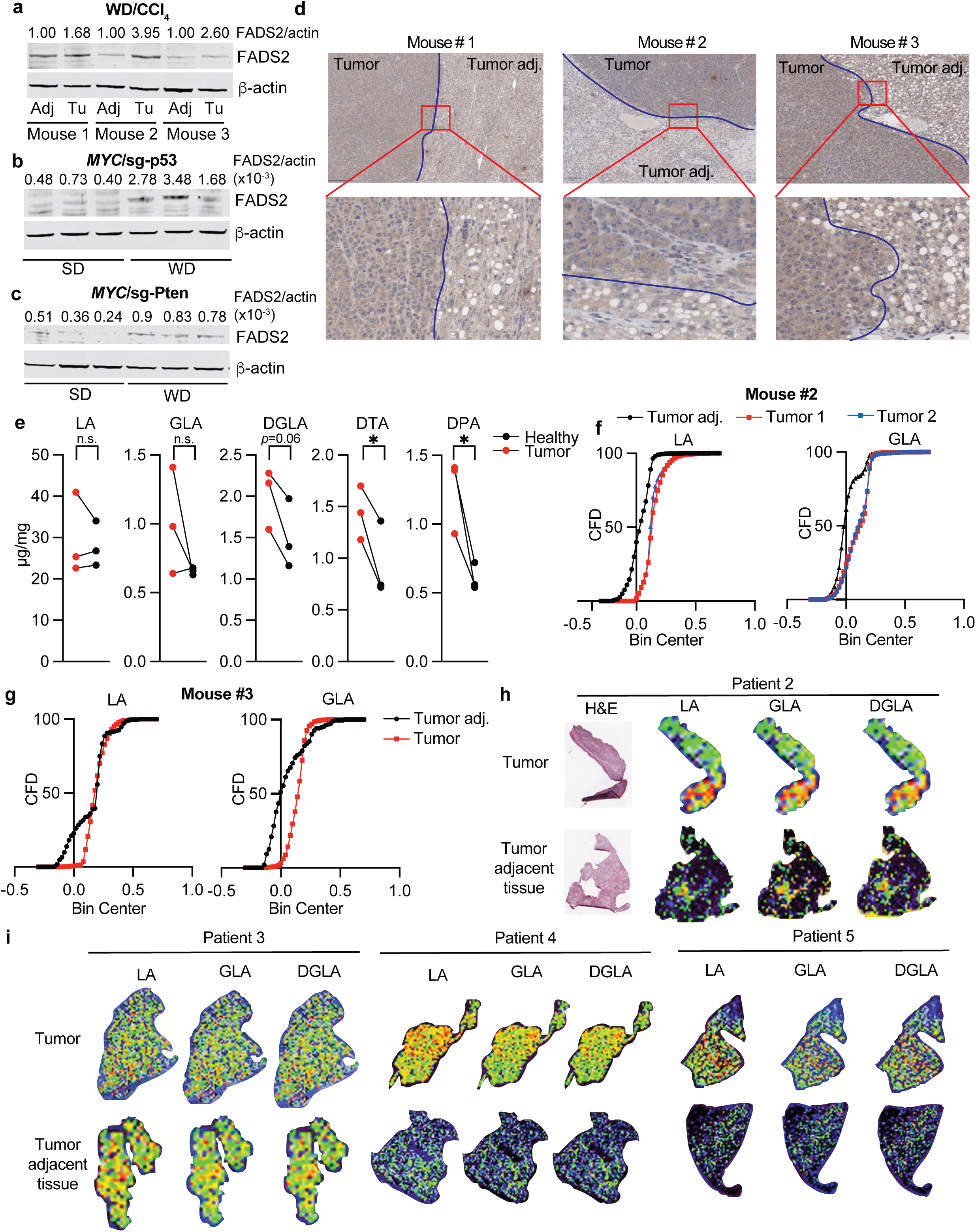
FADS2 and LA-related PUFAs are enriched in tumour tissue in murine and human HCC. **(a – c)** Immunoblot analysis of FADS2 levels in tumour and adjacent tissue in the WD/CCl_4_ (a), MP53 (b), and MPn (c) tumours under SD or WD (n=3); **(d)** FADS2 expression in tumour and adjacent tissue of WD-fed MP53 HCCs; blue lines indicate tumour boundaries; **(e)** Abundance of indicated LA-related PUFAs in tumour and adjacent non-tumor tissue in the WD/CCl_4_ murine model (n=3); **(f & g)** Cumulative frequency distribution (CFD) of Raman spectroscopy signal intensity for LA and GLA in Mp53-lucOS tumours and adjacent liver tissue; **(h & i)** Raman spectroscopy images showing ω-6 PUFA distribution in MASLD-associated HCC patient samples (Columbia University cohort). ns: not significant, *p < 0.05.

**Figure S10:**
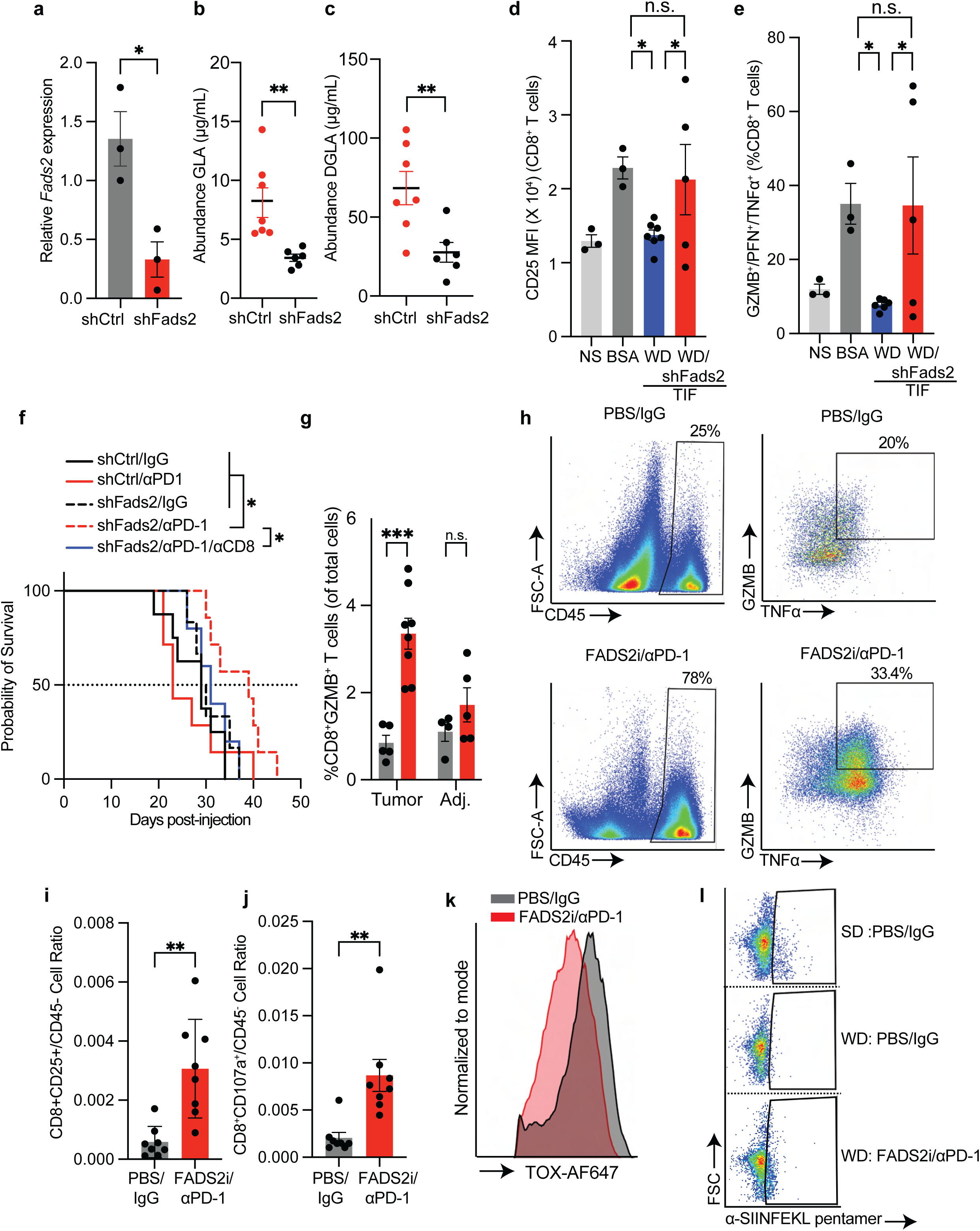
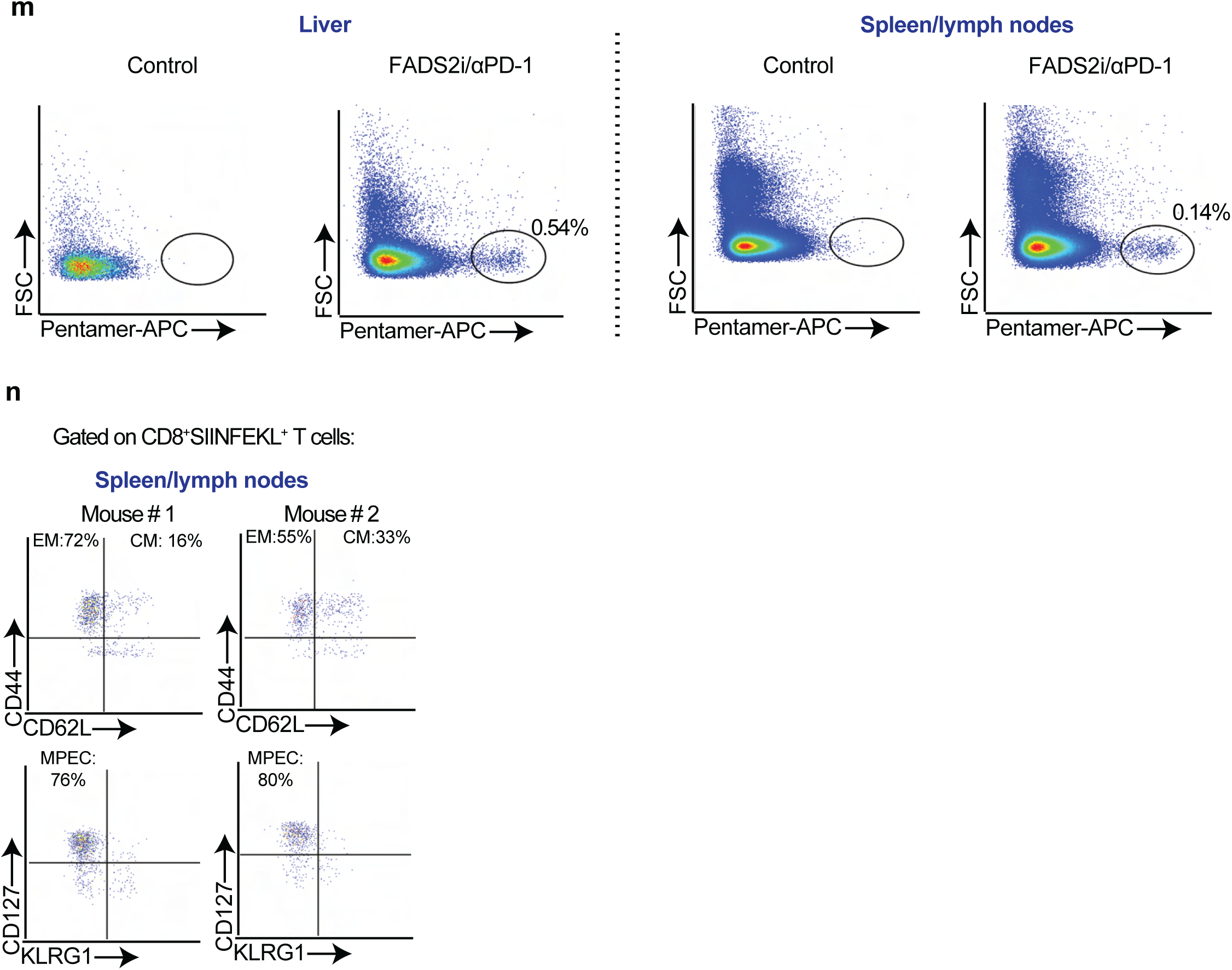
FADS2/PD-1 co-inhibition restores CD8^+^-driven antitumor immunity and promotes immunologic memory. (**a**) Relative *Fads2* expression in indicated MP53 HCCs (n=3); **(b & c)** Abundance of GLA (b) and DGLA (c) in TIF from WD-fed Fads2-proficeint and silenced MP53 tumours (shCtrl: n=7; shFads2: n=6); **(d & e)** Frequency of CD8⁺ T cells expressing CD25 (d) or GZMB, PRF1, and TNFα (e) following stimulation in the presence of TIF derived from WD-fed *Fads2*-proficient or *Fads2*-silenced MP53-lucOS tumours (NS: n=3; BSA: n=3; WD/shC-TIF: n=7; WD/shFads2-TIF: n=5); **(f)** Kaplan–Meier survival curves of WD-fed female mice bearing *Fads2*-proficient or –silenced MP53 HCCs treated as indicated (shCtrl/IgG: n=8; shCtrl/αPD-1: n=7; shFads2/IgG: n=6; shFads2/αPD-1: n=7; shFads2/αPD-1/αCD8: n=5); **(g)** Frequency of GZMB⁺ CD8⁺ T cells among total cells in tumour and adjacent liver tissue from WD-fed MP53-lucOS mice receiving the indicated treatments (Tumour: PBS/IgG: n=5 tumours from 3 mice, FADS2i/αPD-1: n=8 from 3 mice; Adjacent: PBS/IgG: n=4 from 2 mice, FADS2i/αPD-1: n=5 from 3 mice); **(h)** Representative dot plots of infiltrating CD45^+^ cells (left) and GZMB^+^/TNFα^+^ CD8^+^ T cells (right) in WD-fed MP53-lucOS HCCs treated with either PBS/IgG (top) or FADS2i/αPD-1 (bottom); **(i & j)** Relative infiltration of CD25^+^ (i) and CD107a^+^ (h) CD8⁺ T cells in WD-fed MP53-lucOS HCCs under indicated treatments (n=8 tumours from 3 mice); **(k)** Representative histogram of TOX expression in CD8⁺ T cells from WD-treated MP53-lucOS tumours treated as indicated; **(l)** Abundance of SIINFEKL pentamer⁺ tumour-infiltrating CD8⁺ T cells from Mp53-lucOS tumours under the indicated conditions; **(m)** Abundance of pentamer-SIINFEKL⁺ CD8⁺ T cells in liver (left) and lymphoid organs (right) from healthy controls or tumor-free FADS2i/αPD-1–treated mice (MP53-lucOS) (related to Fig. 3a); **(n)** Pentamer-SIINFEKL⁺ CD8⁺ T cells expressing effector or central memory (top) and precursor effector memory (bottom) markers in lymphoid organs of FADS2i/αPD-1–treated mice after tumour clearance (related to Fig. 3a). Results are shown as mean ± SEM. ns: not significant, *p < 0.05, **p < 0.01, ***p < 0.001.

**Figure S11:**
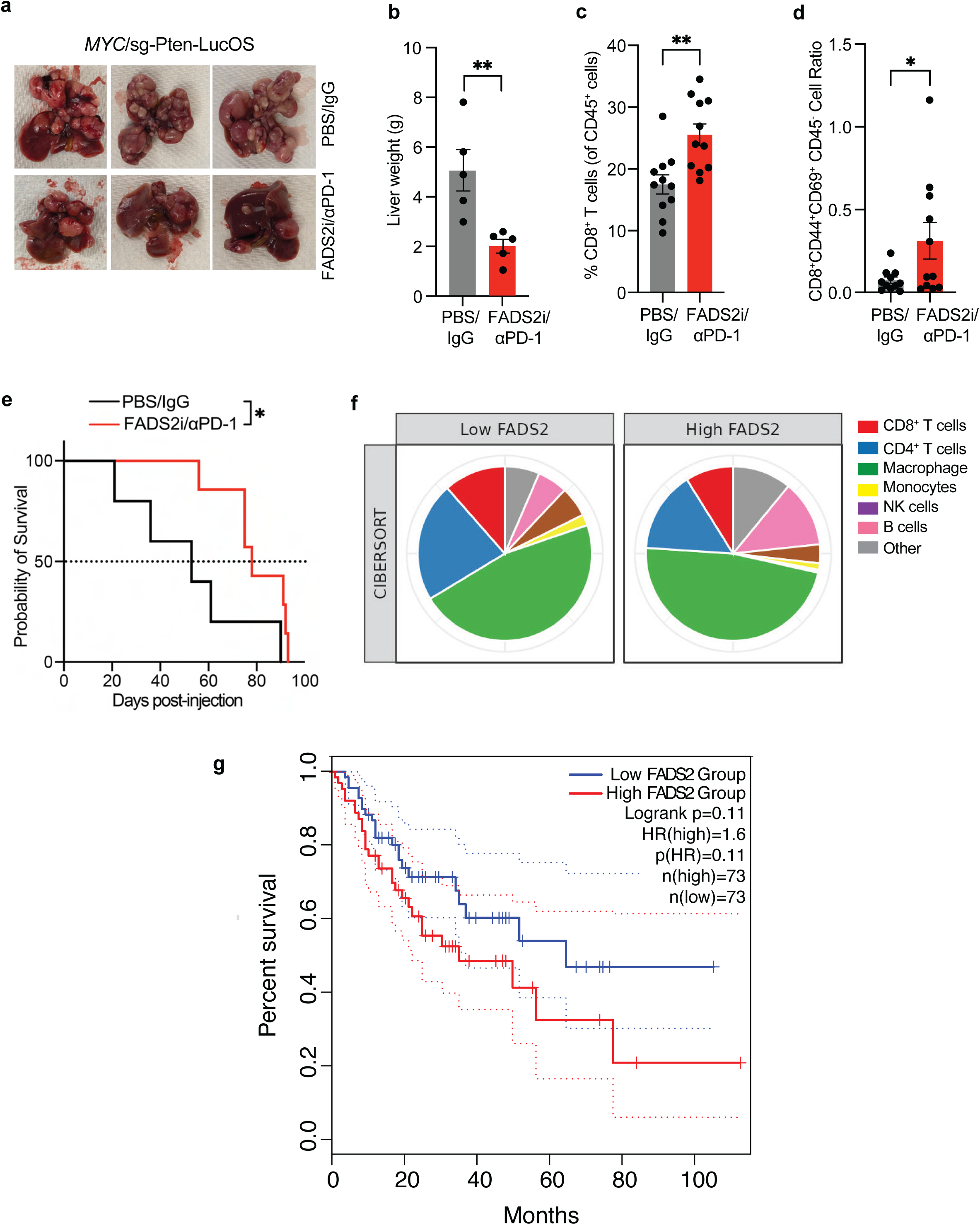
Protective effect of FADS2i/αPD-1 combination in MPn-lucOS and β-catenin-driven HCC murine models. Survival correlation of *FADS2* levels in human HCC. **(a & b)** *Ex vivo* images (a) and weights (b) of time-matched livers from WD-fed female mice bearing MPn-lucOS HCCs and treated as indicated (n=5); **(c & d)** Frequency of CD8^+^ T cells (c) and relative infiltration of CD44^+^CD69^+^ CD8^+^ T cells (d) in WD-fed female mice bearing MPn-lucOS HCCs treated as indicated (PBS/IgG: n=11; FADS2i/αPD-1: n=11); **(e)** Kaplan–Meier survival analysis of WD-preconditioned male mice intrasplenically injected with MYC/sg-p53/β-catenin^τι90^ cells and treated as indicated (n=7); **(f)** CIBERSORT analysis of immune cell abundance in the TCGA-HCC dataset stratified by *FADS2* expression (top and bottom quartile); **(g)** Overall survival of HCC patients stratified by tumoral *FADS2* expression. Results are shown as mean ± SEM. ns: not significant, *p < 0.05, **p < 0.01.

**Figure S12:**
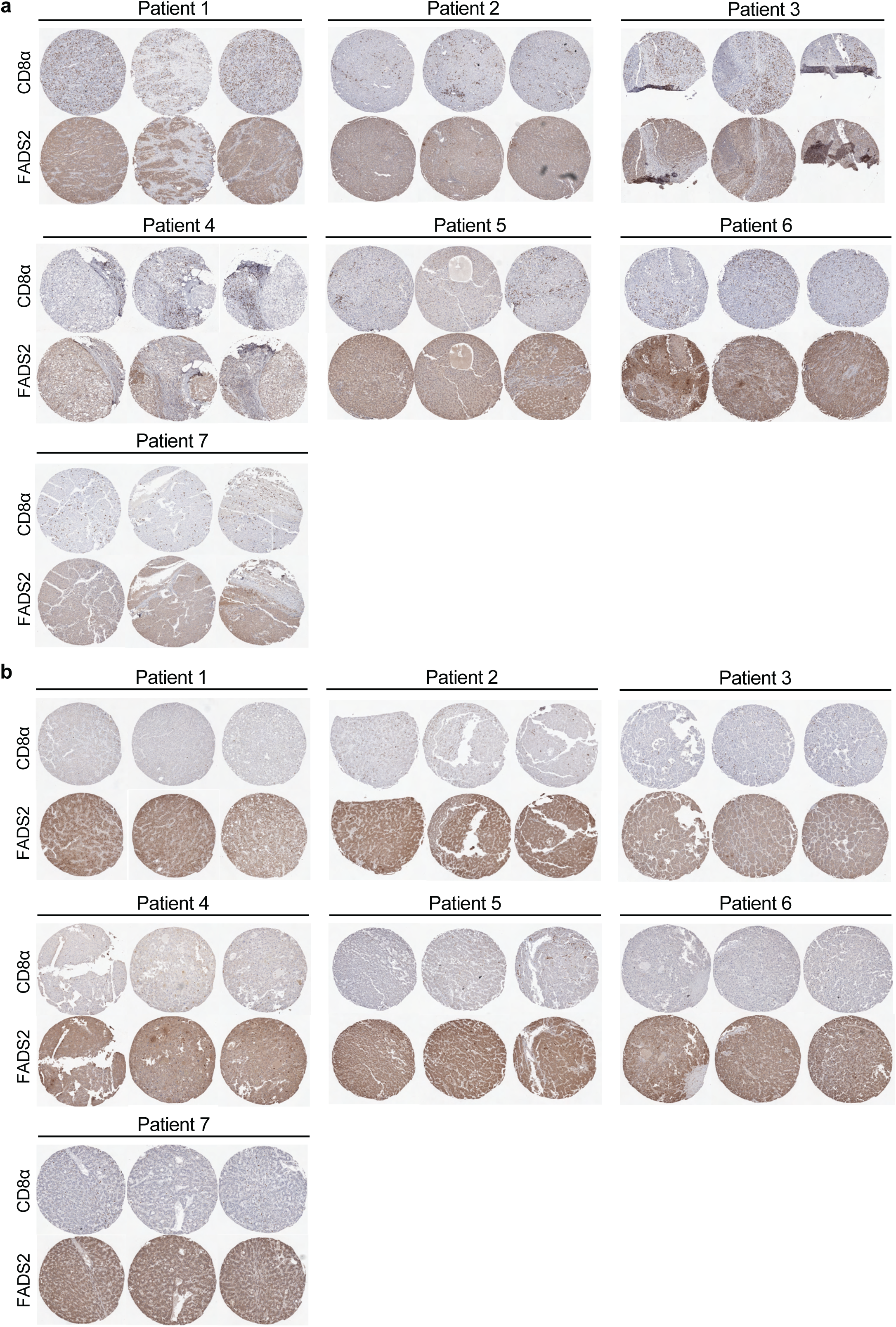
High CD8^+^ T cell infiltration correlates with low FADS2 levels in human HCCs. **(a & b)** TMA images of HCC patient tumours with high CD8⁺ T cell infiltration and low FADS2 expression (n=7) (a), or low CD8⁺ T cell infiltration and high FADS2 expression (n=7) (b).

**Figure S13:**
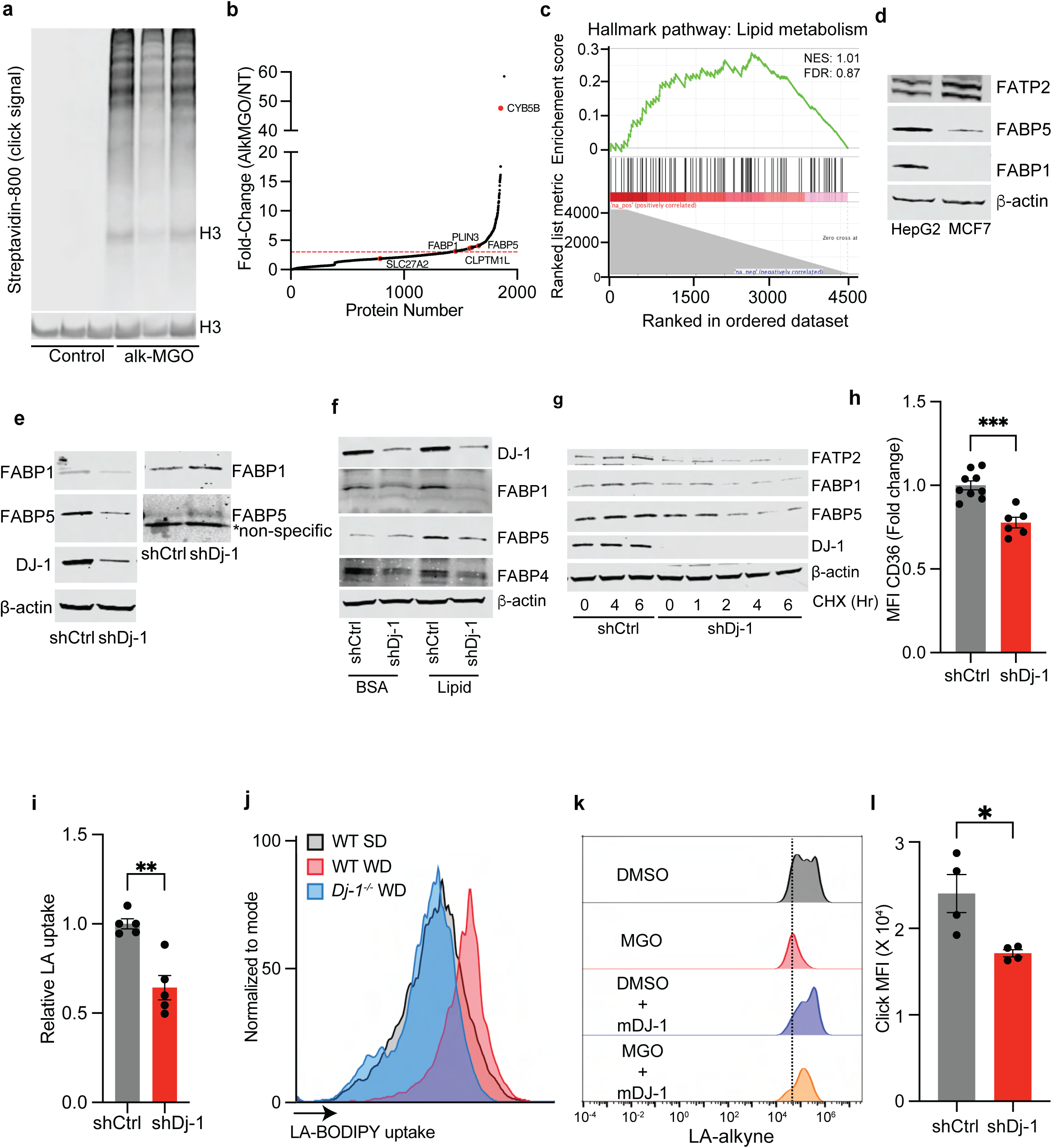
DJ-1 loss promotes MGO-driven hyperglycation of lipid metabolism proteins. **(a)** Immunoblot of click-labelled proteins in DMSO or alkMGO-treated HepG2 cells (3 mM, 5 h); **(b)** Waterfall plot of protein FC (alkMGO-modified/untreated) in HepG2 cells (n=3); lipid metabolism-associated proteins are highlighted; red dashed line indicates FC >2; **(c)** GSEA of lipid metabolism pathway among glycated protein in MCF7 cells (not significant); **(d)** Immunoblot of FATP2, FABP1, and FABP5 proteins in HepG2 and MCF7 cells; **(e)** Immunoblot of streptavidin-bound fractions from *DJ-1*–proficient or –silenced alkMGO treated HepG2 cells (3 mM, 5 h) following azido-biotin Click; **(f)** Immunoblot of indicated lipid-handling proteins in *DJ-1*-proficient or –silenced HepG2 cells ± lipid mixture (LA, OL, PA, 50 µM each, 24 h); **(g)** Immunoblot of indicated lipid-handling proteins in *DJ-1*-sufficient or –deficient HepG2 cells treated with cycloheximide (50 µg/mL, at indicated timepoints); **(h)** Surface expression of CD36 in *DJ-1*-sufficient (n=9) or silenced (n=6) HepG2 cells; **(i)** LA-Bodipy uptake in *DJ-1*-proficient or –silenced HepG2 cells (n=5); **(j)** LA-Bodipy uptake in cancer cells from *Dj-1*-proficient or deficient MP53 tumours under SD or WD regimen; **(k)** LA-alkyne uptake in HepG2 cells ± mDj-1 overexpression following DMSO or MGO treatment (3 mM, 5 h); **(l)** LA-alkyne-derived metabolite levels in CD8⁺ T cells incubated with conditioned media from *DJ-1*–proficient or –silenced HepG2 cells (100 µM, 24 h) (n=4). Results are shown as mean ± SEM. ns: not significant, *p < 0.05, **p < 0.01, ***p < 0.001.

**Figure S14:**
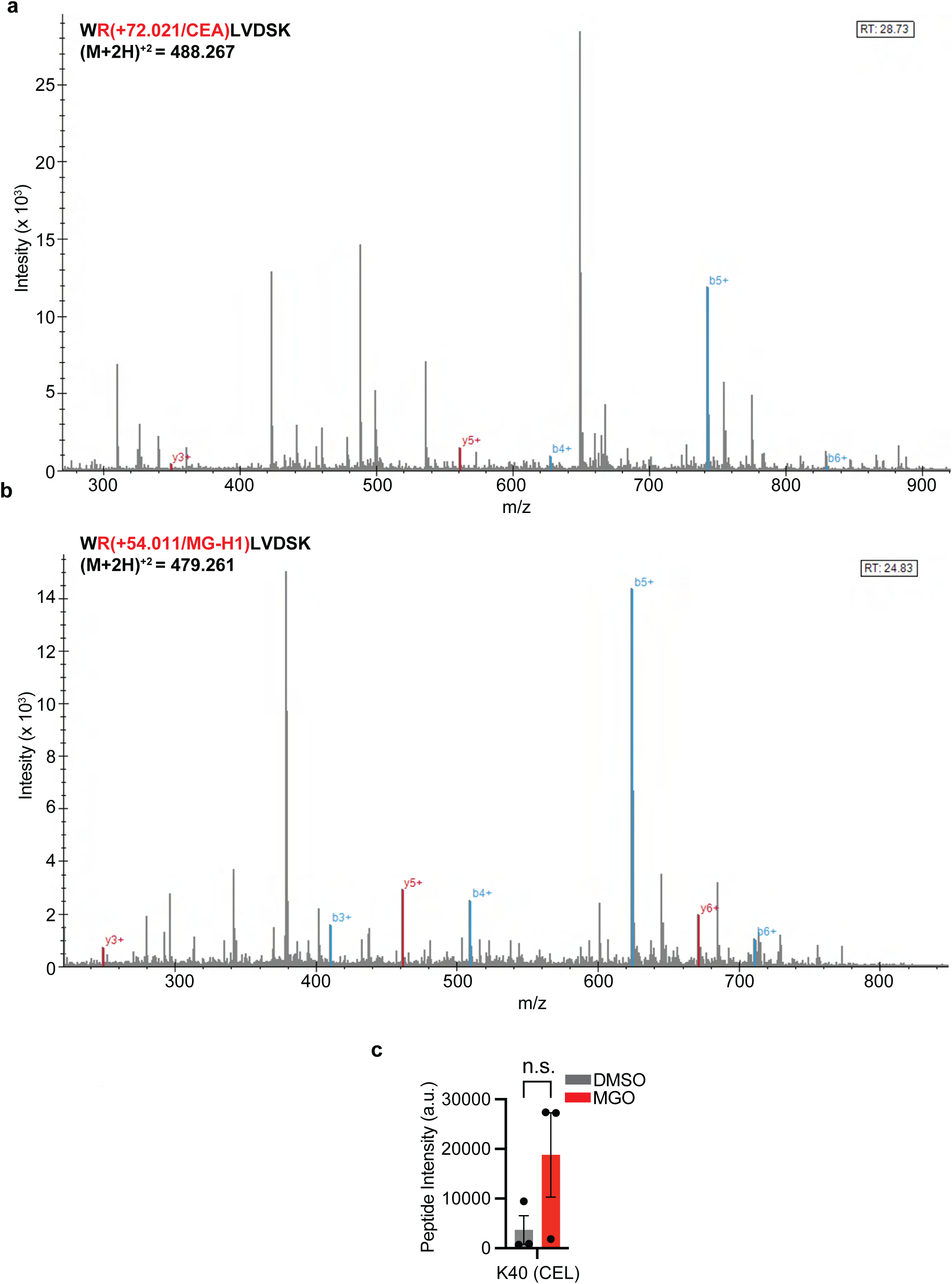
Spectral profiles of the glycation-prone R12 residue in FABP5. **(a & b)** Spectral files of R12-containing peptide with CEA (a) or MG-H1 (b) adducts; **(c)** Relative intensity of K40-CEL (carboxyethyl-lysine) in DMSO– or MGO-treated HepG2 cells (n=3).

**Figure S15:**
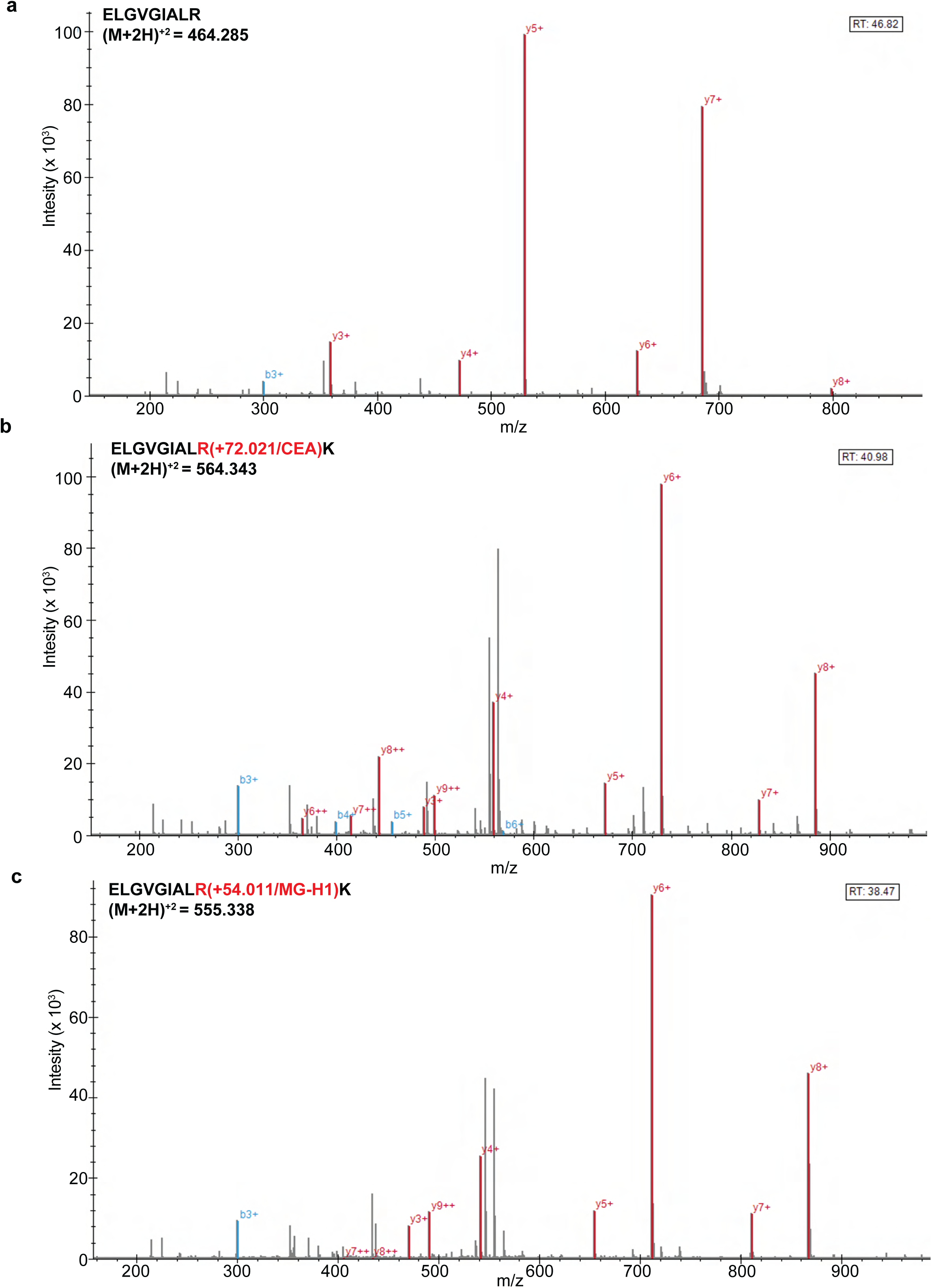
Spectral profiles of the glycation-prone R33 residue in FABP5. **(a – c)** Spectral files of R33-containing unmodified peptide (a), or with CEA (b) or MG-H1 (c) adducts.

**Figure S16:**
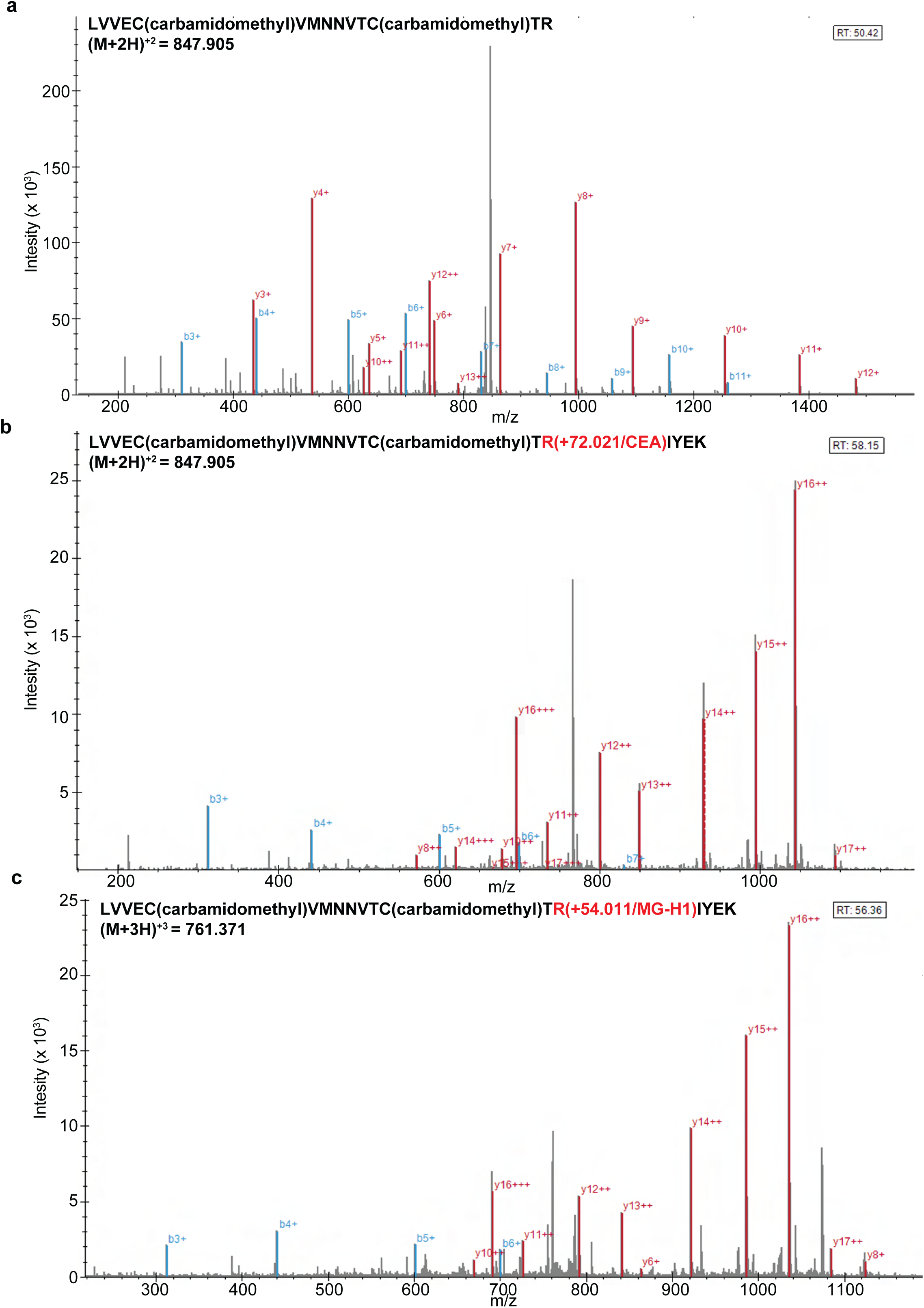
Spectral profiles of the glycation-prone R129 residue in FABP5. **(a – c)** Spectral files of R129-containing unmodified peptide (a), or with CEA (b) or MG-H1 (c) adducts.

**Figure S17:**
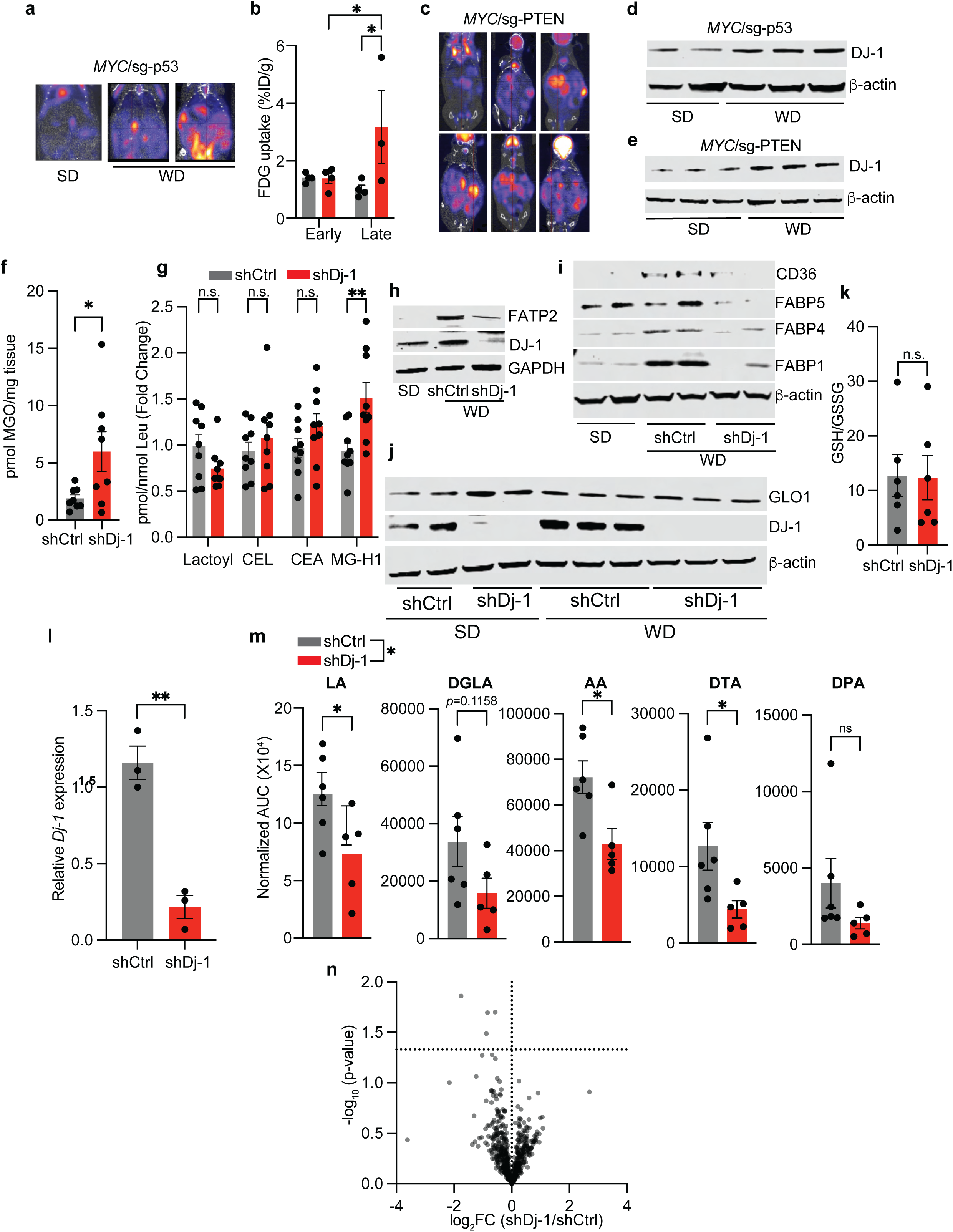
DJ-1 regulates MGO-driven glycation stress and lipid remodeling in WD-driven HCC. **(a, b)** Representative PET images (a) and quantification of FDG uptake (b) in mice bearing Mp53-lucOS HCCs under SD or WD feeding (n ≥ 3); **(c)** FDG uptake in SD– or WD-fed mice harboring MPn HCC, **(d, e)** DJ-1 protein abundance in MP53 (d) and MPn (e) tumours from mice maintained on SD or WD; **(f, g)** Levels of MGO (f) and indicated glycation adducts (g) in *Dj-1*-proficient or – silenced WD-MP53 HCCs (n ≥ 3); CEL: carboxyethyl-lysine; **(h, i)** Immunoblots of indicated lipid-handling proteins in *Dj-1*-proficient or –deficient SD– or WD-fed MP53 HCC; **(j)** GLO1 protein expression in indicated MP53 HCC; **(k)** Reduced-to-oxidized glutathione in WD-fed *Dj-1*-proficient or –silenced MP53 tumours (n = 6); **(l)** Relative *Dj-1* expression in indicated MP53 HCCs (n = 3); **(m)** Relative abundance of LA and downstream PUFAs in WD-fed *Dj-1*-proficient (n = 6) or – defcient (n = 5) MP53 HCCs; **(n)** Volcano plot of differentially abundant lipids in *Dj-1*-proficient versus Dj-1-silenced Mp53-lucOS HCCs from WD-fed mice (n = 4). Results are shown as mean ± SEM. ns: not significant; *p < 0.05; **p < 0.01.

**Figure S18:**
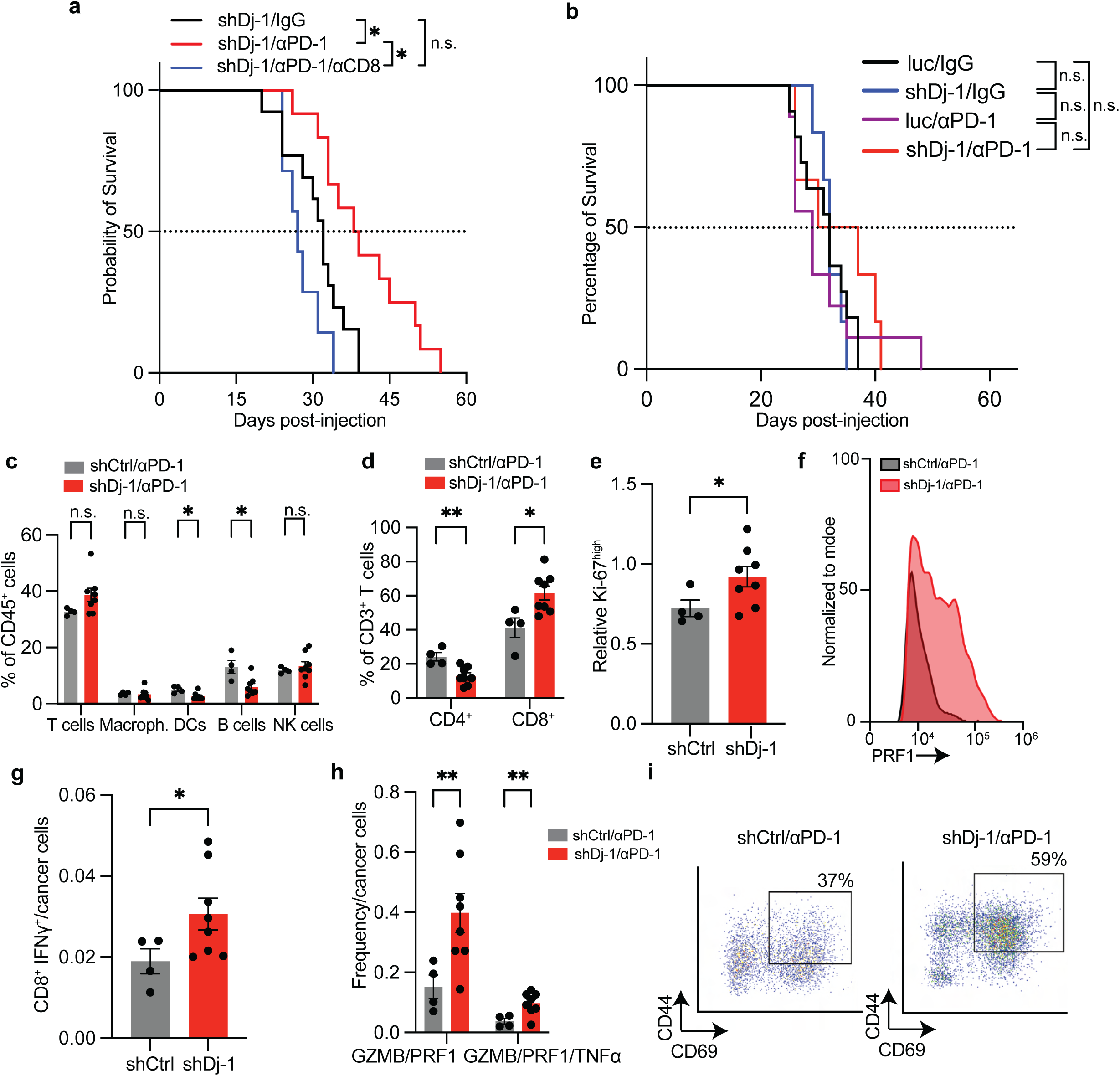
DJ-1 loss restores CD8⁺ T cell function and sensitizes WD-driven HCCs to immunotherapy. (**a**) Kaplan–Meier survival curves of WD-fed female mice bearing *Dj-1*–proficient or –deficient MP53 HCCs treated as indicated (shDj-1/IgG: n=13; shDj-1/αPD-1: n=12, shDj-1/αPD-1/αCD8: n=7); **(b)** Kaplan–Meier survival curves of SD-fed female mice bearing *Dj-1*-proficient or –deficient MP53 HCCs treated with indicated antibodies (luc/IgG: n=11; shDj-1/IgG: n=6; luc/αPD-1: n=9; shDj-1/αPD-1: n=6); **(c & d)** Frequency of indicated immune populations relative to CD45⁺ cells (c) or to CD3⁺ T cells (d) in *Dj-1*–proficient or –deficient MP53 HCCs treated as indicated (shCtrl/αPD-1: n=4 tumours from 2 mice; shDj-1/αPD-1: n=8 tumours from 3 mice); **(e)** Relative levels of Ki-67^high^ in CD8^+^ T cells in *Dj-1*-sufficient (n=4) or –silenced (n=8) MP53 HCCs treated with αPD-1; **(f)** Representative histogram of PRF1 levels in indicated experimental groups**; (g & h)** Relative infiltration of IFNγ⁺ (g) and frequency of GZMB⁺PRF1⁺ or GZMB⁺PRF1⁺TNFα⁺ (h) CD8⁺ T cells in *Dj-1*–proficient (n=4 tumours from 2 mice) or –deficient (n=8 tumours from 3 mice) MP53 HCCs treated with αPD-1; (i) Representative dot plots of CD8^+^ T cells co-expressing CD69 and CD44 in indicated tumours. Results are shown as mean ± SEM. ns: not significant, *p < 0.05, **p < 0.01.

**Figure S19:**
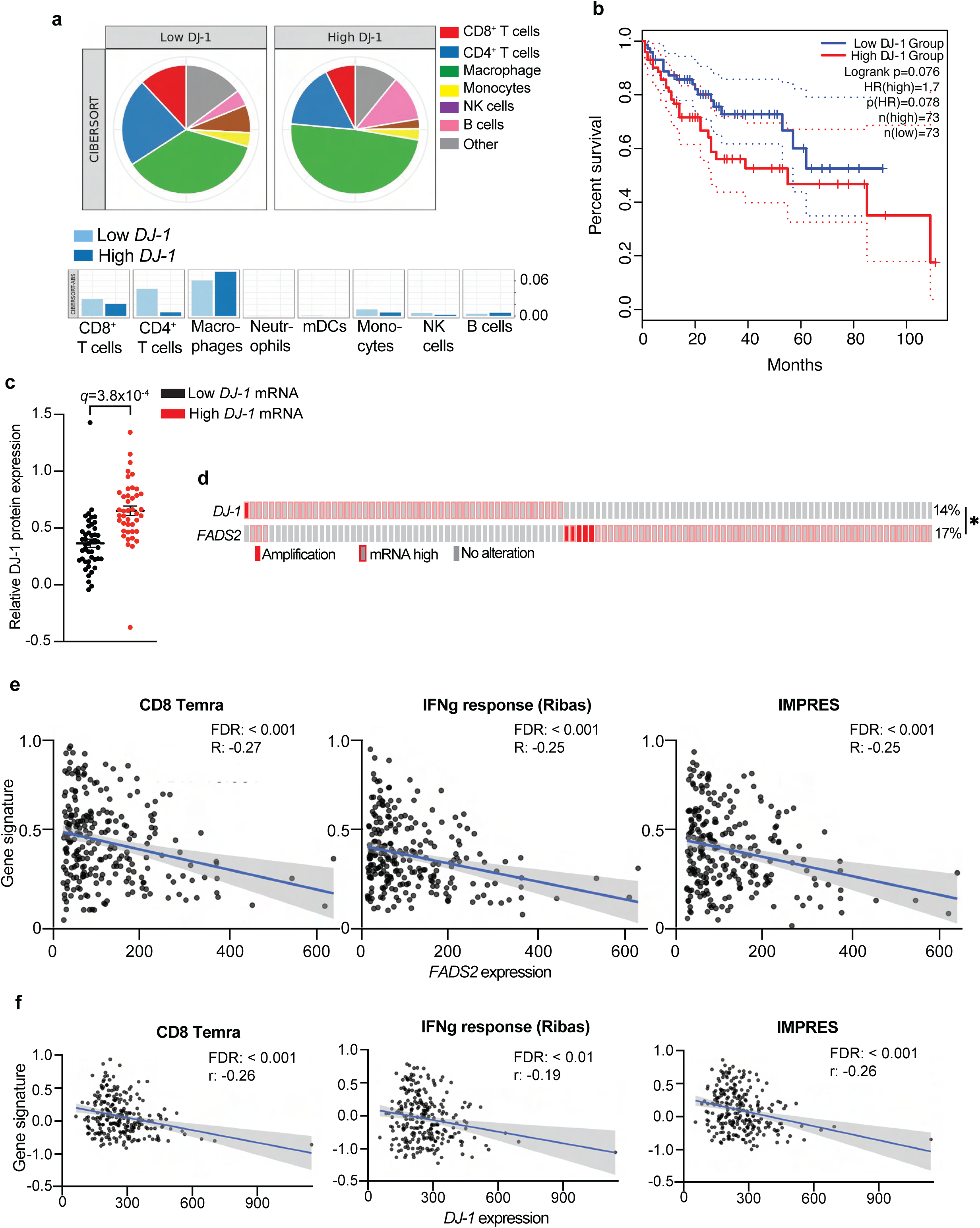
Low *DJ-1* expression correlates with better survival, higher CD8⁺ T cell infiltration, and improved T cell function in human HCC. **(a)** CIBERSORT analysis of immune cell infiltration in TCGA-HCC patients stratified by high and low *DJ-1* (*PARK7*) expression quartiles; **(b)** Overall survival of HCC patients stratified by high *DJ-1* mRNA levels; **(c)** Relative DJ-1 protein levels in HCC patients stratified by low (n=45) and high (n=42) *DJ-1* mRNA expression; **(d)** OncoPrint of TCGA-HCC patients showing mutually exclusive amplification and/or high mRNA expression of *DJ-1* (*PARK7*) and *FADS2*; **(e & f)** Correlation plots showing the association of *DJ-1* (e) or *FADS2* (f) expression with gene signatures predictive of cytotoxicity and immunotherapy-response. *p < 0.05.

